# An integrated multimodal pan-organ atlas of the female reproductive system across the lifespan contextualises gynaecological pathologies

**DOI:** 10.64898/2026.06.10.731198

**Authors:** Celeste E Cohen, Antonio Parraga-Leo, Leticia Rodríguez-Montes, Christina E Kim, Marie Moullet, Ana Paredes, Valentina Lorenzi, Roser Vilarrasa-Blasi, Alexander V Predeus, Krzysztof Polanski, Erick Armingol, Cecilia Icoresi Mazzeo, Brian Rous, Laura Fachal, Bradley T Harris, Carmen Sancho-Serra, Iva Kelava, Magda Marečková, Loren Méar, Nicole D Ulrich, Dilara N Anbarci, Lijiang Fei, Taylor Schissel, Fu Wei, Gustaw Eriksson, Tanja Turunen, Josep Marí Alexandre, Marie-Therese Bammert, Nilay Kuscu, Vicente Pérez-García, Juan Gilabert-Estellés, Frédéric Chalmel, Christian Becker, Krina T Zondervan, Alain Chedotal, Suzannah Williams, Tereza Cindrova-Davies, Nardhy Gomez-Lopez, Semir Beyaz, Antoine D Rolland, Anindita Basu, Aymara Mas, Felipe Vilella, Elisabet Stener-Victorin, Pauliina Damdimopoulou, Carlos Simon, Susana M Chuva de Sousa Lopes, Sarah A Teichmann, Miriam Baumgarten, Cecilia Lindskog, Ariella Shikanov, Saher Sue Hammoud, Carl A Anderson, Luz Garcia-Alonso, Roser Vento-Tormo

## Abstract

Single cell transcriptomics has transformed our knowledge of reproductive tissues, yet studies remain largely organ-specific and temporally limited, leaving an incomplete picture of how cell types are distributed across the reproductive system over a lifetime. Gynaecological conditions affect more than one in four females and frequently span multiple organs and life stages. To advance our understanding and treatment of these conditions, an integrated cellular reference is essential. Here we present the *Human Female Reproductive System Cell Atlas v1*: a single-cell transcriptomic resource integrating more than 2M cells across the ovary, fallopian tube, uterus, cervix and vagina over the lifespan and menstrual cycle, further integrated with spatial transcriptomics and chromatin accessibility profiling to define 210 cell types through community-based annotation. Cross-organ integration resolves shared and organ-specific cellular states, identifying uterine-specific perivascular populations lining uterine spiral arteries, hypoxia-sensing type 3 innate lymphoid cells (ILC3s) enriched in the uterus, and lipid-associated macrophages with distinct subsets in each reproductive organ, including a previously undescribed population shared between the uterus and fallopian tube. Cross-organ integration enables detection of ectopic epithelial populations in otherwise healthy donors, including endometrial-like cells within a paediatric ovary consistent with early endometriosis. Integration with genome-wide association studies (GWAS) reveals that risk variants for major gynecological conditions act in mesenchymal cell states defined by specific transcriptional programmes and spatial or temporal context – for instance, heavy menstrual bleeding risk is enriched in basal fibroblasts (SFRP5⁺) of the regenerative endometrial compartment. An integrated chromatin accessibility atlas provides peak-to-gene maps across reproductive cell types, enabling nomination of disease effector genes and providing the first regulatory evidence linking a Polyendocrine Metabolic Ovarian Syndrome (PMOS) risk locus to *INHBB* in granulosa cells. Together, this resource establishes a cellular and molecular framework for reproductive biology and the pathogenesis of neglected gynaecological conditions.

## Main

The female reproductive system is one of the most dynamically regulated in the human body. Its tissues undergo continuous, hormone-driven change across the lifespan, from puberty through menopause, and throughout the menstrual cycle, alternating between states of growth, differentiation and regeneration each month. This dynamism spans the five organs of the internal genitalia: the ovary, fallopian tube, uterus, cervix and vagina. Each organ is structurally and cellularly distinct, reflecting specialised roles, yet they collectively function as a coordinated whole to achieve reproductive function. Most extreme in this dynamism is the endometrium (the mucosal lining of the uterus) which regenerates with each menstrual cycle, doing so more than 400 times across the reproductive lifespan^1^.

Because common laboratory models do not fully recapitulate human reproductive physiology^2–4^, understanding how these tissues function in humans requires direct study of human samples. Their dynamic regulation further requires a holistic view across multiple temporal scales, from broad life stages (development, childhood, puberty, reproductive age and menopause) to the finer timescale of days, capturing cellular state shifts across the menstrual cycle. Single-cell and spatial genomics have generated comprehensive maps of cell types and states within individual reproductive organs at unprecedented resolution. These efforts have revealed novel insights into reproductive biology, including the dynamism of progenitor populations in the developing reproductive organs^5–9^, the underappreciated cellular complexity of the adult reproductive tissues^10–22^, and the specialised immune programmes central to reproductive function^19,23,24^.

Despite this progress, no integrated atlas spanning all five reproductive organs and their temporal dynamics exists. Such a resource would offer capabilities beyond single-organ studies^25–27^: it would enable comparison of shared versus organ-specific cell states, clarifying the cellular basis of tissue-specific homeostasis, increase the power to detect rare populations that are difficult to characterise in isolation and illuminate the biology of gynaecological conditions that involve multiple organs. Endometriosis, for example, involves the displacement of endometrial-like tissue to ectopic sites including the ovary^28^; high-grade serous ovarian cancer (HGSOC), the most lethal gynaecological malignancy, presents in the ovary but existing evidence indicates fallopian tube origin^29^. More broadly, the cell of origin remains unknown for most malignancies of the reproductive tissues. Understanding such cross-organ relationships demands an integrated reference that spans tissues, development and the menstrual cycle.

A cross-organ atlas is also an essential resource for interpreting the genetic basis of gynaecological conditions. Large-scale human genetics studies have identified hundreds of genomic loci associated with gynaecological conditions, yet translating associations into biological mechanisms remains challenging. Genetic risk for complex disease is mediated by specific cell types: variants affect gene activity within the cell states where they operate, driving disease. Identifying how genetic variants manifest in disease therefore requires knowing which cell states exist across the affected tissues, when they are present, and where. For reproductive disorders, which span multiple organs and fluctuate with the menstrual cycle and developmental stage, this information is only accessible through an integrated, temporally resolved atlas. A further challenge is that approximately 93% of disease-associated variants across complex traits lie in non-coding regions of the genome^30^, where their effects on gene regulation are cell type- and context-dependent. Nominating the specific gene a non-coding variant controls requires direct characterisation of the chromatin regulatory landscape at cell-state resolution.

Here we present the *Human Female Reproductive System Cell Atlas v1*: a single-cell transcriptomic resource spanning the ovary, fallopian tube, uterus, cervix and vagina across the lifespan (from development to menopause, including paediatric stages for the ovaries) and the menstrual cycle (for the uterus and fallopian tubes). The atlas defines a consensus reference of cell types and states across all five organs, with spatial transcriptomics used to ground cell type annotations in their anatomical context, and cell nomenclature established by the Human Cell Atlas Reproductive Network jointly with domain experts. Integrated chromatin accessibility profiling (scATAC-seq) further maps links between regulatory DNA regions and the genes they control at cell state resolution, enabling non-coding disease variant interpretation. Together, this pan-organ view resolves shared and tissue-restricted cellular programmes invisible to single-organ studies, clarifies the cellular basis of reproductive tissue physiology and histopathology, and provides a reference for linking disease-associated genetic variation to the cells and molecular programmes in which it acts.

## Results

### Human Female Reproductive System Cell Atlas v1

We assembled the unified *Human Female Reproductive System Cell Atlas v1* by integrating single-cell transcriptomic datasets from the human ovary, fallopian tube, uterus, cervix and vagina spanning fetal development through postmenopause (**Figure 1a** and **Supplementary Table 1a**). The atlas combines publicly available scRNA-sequencing datasets from multiple studies and centres (**Figure 1b**), supplemented by 1) newly generated fallopian tube data to improve coverage of this organ, and 2) spatial transcriptomics data generated across multiple reproductive tissues for this study (**Supplementary Table 1b,c**). In total, we resolved 2,235,448 high-quality cells from 291 donors and 27 datasets after unified re-preprocessing, ambient RNA removal and quality control (**Figure 1a,b** and **Supplementary Note 1.1**). Together these datasets allowed us to define the identities and location of 210 fine cell types within female reproductive tissues across lifespan (**Figure 1c-e, Extended Data Fig. 1a-b** and **Extended Data Fig. 2a-c**).

**Figure 1.**
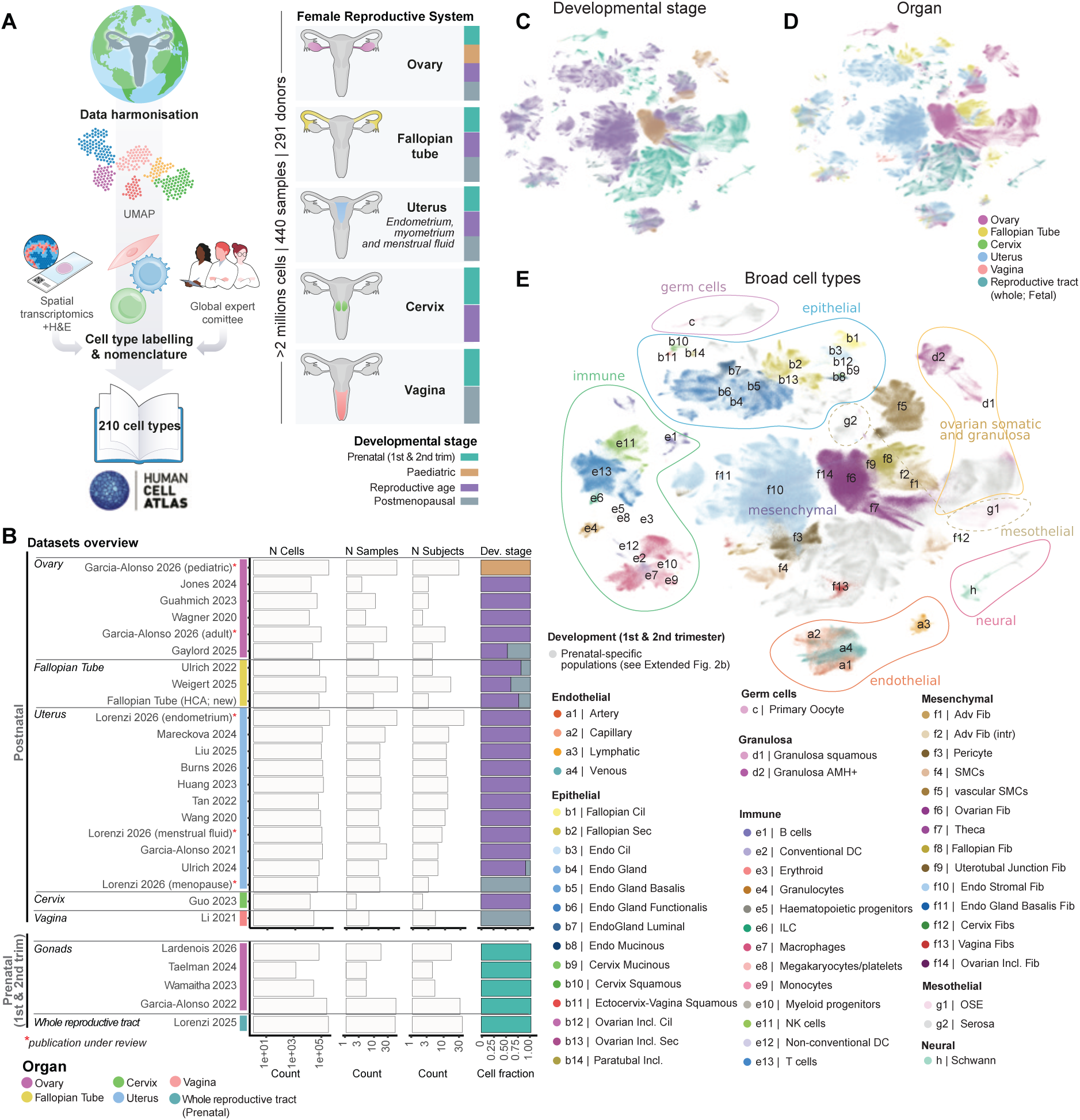
Human Female Reproductive System Cell Atlas v1 overview. **A,** Schematic overview of the organs, developmental stages and samples included in the atlas, together with the data harmonisation and reprocessing workflow, cell type annotation, validation and consultation within the HCA Reproduction Network and domain experts. **B,** Bar plots representing the number of cells, samples, donors and developmental stage proportions (x-axis) contributed by each dataset included in the atlas (y-axis) for the postnatal and prenatal organs of the female reproductive system. **C-E,** Batch-corrected Uniform Manifold Approximation and Projection (UMAP) visualisation manifold of the scRNA-seq dataset (n = 2,235,448 cells; n = 291 donors), coloured by developmental stage (“C“), organ (“D”) and broad cell type annotation in postnatal donors (“E”; "`broad_celltype`" labels). Prenatal-specific broad cell categories are shown in Extended Data Fig. 2b. Cil, ciliated; Sec, secretory; Endo, endometrial; Incl, inclusion; DC, dendritic cell; ILC, innate lymphoid cell; NK, natural killer; Adv, adventitial; Fib, fibroblast; Intr, interstitial; SMC, smooth muscle cell; OSE, ovarian surface epithelium.

To achieve consistent annotation across organs and studies, we developed a hierarchical integration and annotation framework based on scVI integration and iterative lineage-specific re-analysis (**Supplementary Note 1**). Cells were first classified into eight major lineages (epithelial, mesothelial, mesenchymal, endothelial, immune, granulosa/ovarian supporting, germ cell and peripheral nervous system populations) before being resolved into progressively finer pan-organ and tissue-specific states (**Extended Data Fig. 1b** and **Extended Data Fig. 2c**). Cell identities were assigned using canonical marker genes, *de novo* identified markers, sample metadata and spatial localisation with matched Visium or Xenium (for ovaries, fallopian tubes and uterus), with additional spatial data generated for adult fallopian tubes (**Supplementary Table 1c**). All populations were mapped to a structured nomenclature framework developed within the Human Cell Atlas Reproductive Network in consultation with domain experts and pathologists (**Supplementary Table 2** and **Supplementary Note 2.1**). Cellxgene objects are available at https://www.reproductivecellatlas.org/HCAreproductive/v1/.

The following results sections describe the major cell types and transcriptional programmes identified across organs and lineages, with a focus on paediatric/adult stages. For prenatal cross-organ comparison, we refer readers to Lorenzi et al., 2026^9^.

### A pan-reproductive fibroblast hierarchy with uterine-specific perivascular cells

To build a unified mesenchymal reference across the reproductive system, we re-annotated populations that previous atlases had variably grouped as broad stromal or vascular cells. Beyond tissue-specific interstitial fibroblasts (39 cell states), the atlas identified three non-interstitial fibroblast compartments conserved across reproductive tissues: perivascular, smooth muscle cells (SMCs) and adventitial. Perivascular mural cells (pericytes and vascular SMCs (vSMCs)) and visceral SMCs were consistent with populations previously described in the original datasets (**Figure 2a,b** and **Extended Data Fig. 3a**). In contrast, adventitial fibroblasts emerged as a distinct transcriptomics compartment that had been largely overlooked in prior atlases, likely owing to their scarcity and their annotation as generic stromal fibroblasts (**Extended Data Fig. 3a** and **Supplementary Note 2.1**).

**Figure 2.**
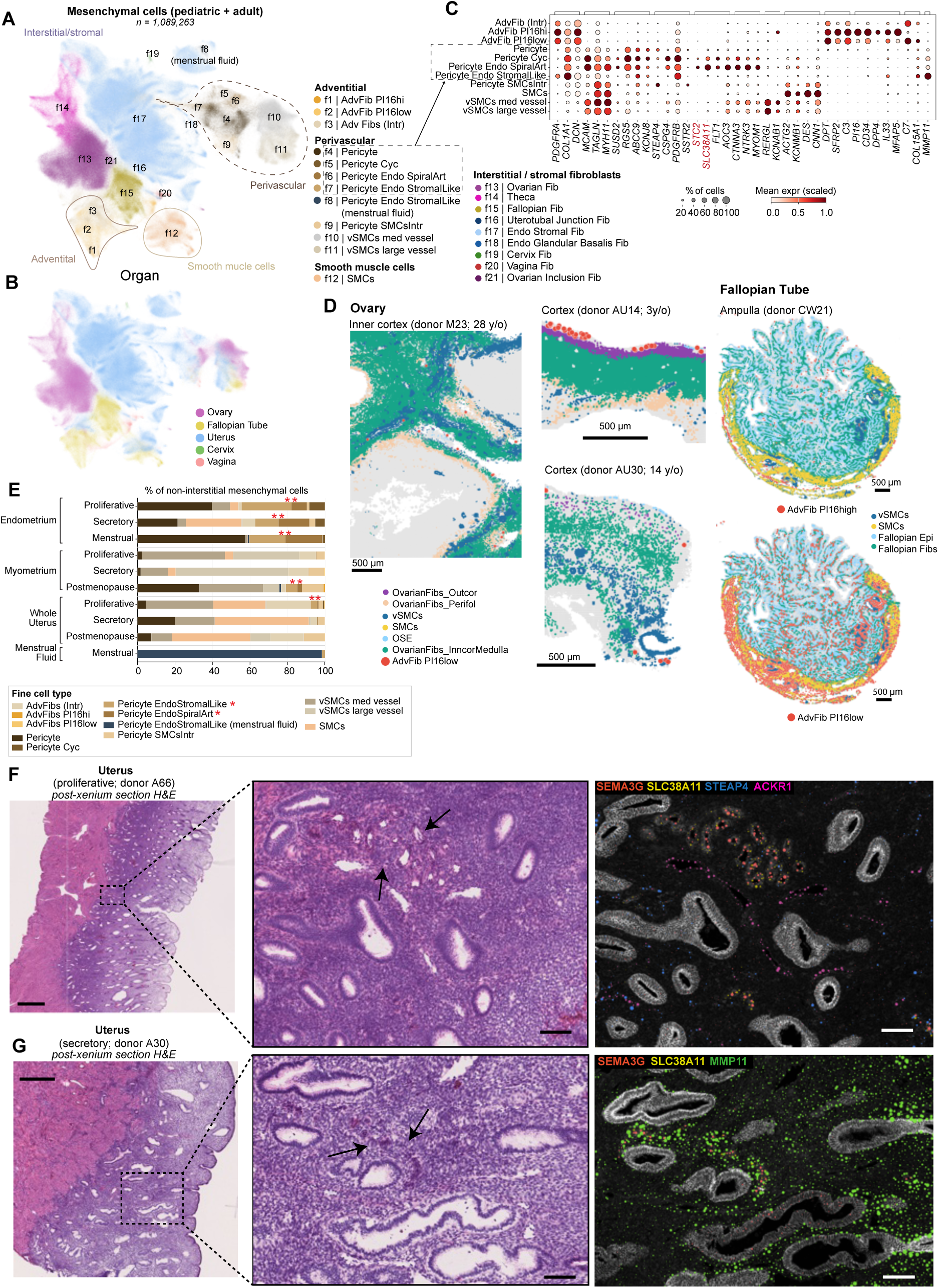
Mesenchymal lineage hierarchy across postnatal female reproductive organs. **A-B,** Batch-corrected UMAP visualisation manifold of the scRNA-seq dataset of all mesenchymal cells (n=1,089,263; "`broad_celltype`" labels for interstitial fibroblasts and “fine_celltype” labels for non-interstitial fibroblasts) from the postnatal donors in the Human Female Reproductive System Cell Atlas v1, coloured by broad cell state (“A”) and organ (“B”). **C,** Dot plot showing log-transformed, min–max-normalised expression of selected marker genes (x-axis) across postnatal mesenchymal cell type categories (y-axis). Labels correspond to `fine_celltype`, except for interstitial populations, which are grouped at the `broad_celltype` level. **D.** Visualisation of predicted adventitial and selected neighbouring cells on one fallopian tube section (ampulla; donor CW21, reproductive age; showing both adventitial PI16-high and adventitial PI16-low) and three ovarian cortex sections (donors M23, 28 years old; AU30, 14 years old; and AU14, 3 years old; showing adventitial PI16-low) profiled using Visium HD. Selected adventitial and selected broad cell-type labels were transferred from the Human Female Reproductive System Cell Atlas v1 onto Visium HD 8 µm bins using the TACCO tool. Scale bar = 500µm. **E.** Bar plot showing the cellular composition of uterine mesenchymal populations ("`fine_celltype`" labels) according to sampling source (endometrium, myometrium, whole uterus or menstrual fluid) and menstrual phase or menopausal status. **F-G**. Visualisation of selected vascular marker transcripts in two representative whole uterine sections from adult donors A66 (reproductive age, proliferative phase), and A30 (reproductive age, mid-secretory phase), profiled using a bespoke Xenium 480-gene panel. The left panels show the corresponding haematoxylin and eosin (H&E) histology. **F**. Xenium visualisation of marker genes for arterial endothelium (*SEMA3G*; red), venous/vessel endothelium (*ACKR1*; pink), spiral artery pericytes (*SLC38A11*; yellow), and classical pericytes (*STEAP4*; blue). **G**. Xenium visualisation of arterial endothelium (*SEMA3G*; red), spiral artery pericytes (*SLC38A11*; yellow) and EndoStromalLike pericytes (*MMP11*; green). Arrows point to Spiral Arteries in the tissues. Scale bar = 1mm in full thickness H&E images and 100µm in magnified insets. Adv, adventitial; Fib, fibroblast; Intr, interstitial; Cyc, cycling; Endo, endometrial; Art, arteries; SMC, smooth muscle cell; med, medium; expr, expression; Outcor, outer cortex; Perifol, perifollicular; vSMC, vascular smooth muscle cell; OSE, ovarian surface epithelium; Inncor, inner cortex; Epi, epithelium.

Pan-reproductive adventitial fibroblasts are defined by expression of *DPT*, *SFRP2* and *C3*, forming a transcriptional continuum from *PI16*^hi^ (named “*AdvFib PI16hi*”) to *C7*/*COL15A1*^hi^ states (labeled “*AdvFib PI16low*” and “*Adv Fib (Intr)*”; **Figure 2a,c**), consistent with the universal fibroblast populations previously described^31,32^. *AdvFib PI16*^hi^ states are mostly found in the fallopian tube ampulla and isthmus (99% of *PI16*^hi^ cells; **Extended Data Fig. 3b**), where spatial transcriptomics localise them to its characteristic subserosal (adventitial) connective tissue in the ampulla (**Figure 2d**). In contrast, *AdvFib PI16low* and *Adv Fib (Intr)* (i.e. *C7*/*COL15A1*^hi^ states) are distributed across tissues occupying the mucosal interstitium of the fallopian tube, the subserosa region beneath the ovarian surface epithelium and adventitial positions around vasculature in the ovary, fallopian tube and the myometrium in the uterus (**Figure 2d** and **Extended Data Fig. 3c**).

Perivascular mural cells included shared and tissue-specialised states. Conserved populations included canonical pericytes (*RGS5*^+^/*STEAP4*^+^), associated with capillaries and small vessels, and vSMCs (*RERGL*^+^), associated with medium/large vessels (**Figure 2c,d**), both of which are shared with other non-reproductive tissues^27^. Although both compartments were detected across organs, the endometrium and ovarian cortex showed few large-vessel vSMCs and abundant cycling, pericyte-associated microvasculature (**Extended Data Fig. 3b**). This pattern matches the regenerative demands of these tissues, with the endometrium undergoing repeated monthly rebuilding, and the ovarian cortex supporting continuous follicle growth and atresia.

The uterus harbours a specialised vasculature characterised by spiral arteries, small coiled vessels that extend into the functional endometrial layer and undergo cyclical remodeling with each menstrual cycle. Cross-organ analysis identified two additional perivascular cell populations (*RGS5*^+^) restricted to the uterus (**Figure 2c,e)**. Spiral artery-associated pericytes (called “*Pericyte EndoSpiralArt*”) were present throughout the uterus but enriched in the endometrium (**Figure 2e)**, co-expressed canonical pericyte markers alongside *SLC38A11*, *STC2* and *FLT1*, and lacked the vSMC marker *RERGL* (**Figure 2c**). Spatially, they surround *SEMA3G*^+^ arterial endothelial cells within spiral arteries, further establishing their specialised role in the uterine arterial vasculature (**Figure 2f** and **Extended Data Fig. 3d;** spiral arteries indicated with arrows).

The endometrial perivascular-stromal population (called “*Pericyte EndoStromalLike*”) co-expresses pericyte (*RGS5*) and stromal (*PDGFRA*, *MMP11*) markers (**Figure 2c**). Its intermediate transcriptional profile, together with its localisation around small vessels and spiral arteries across both the proliferative and secretory phases in the endometrium (**Figure 2f,g** and **Extended Data Fig. 3d**), supports a perivascular-stromal identity specific to the endometrium. Its perivascular localisation is consistent with earlier descriptions of endometrial stromal precursors, supporting a possible link to stromal fibroblast intermediates of pericyte origin^33–36^.

Altogether, pan-organ integration defined the adventitial and perivascular compartments in healthy organs, including uterine-specific pericyte subsets adapted to the specialised demands of the uterine vasculature.

### Reproductive mucosal tissues are enriched in tissue-resident innate immune lineages

Tissue-resident immune cells cooperate with the mesenchymal compartment to maintain reproductive homeostasis, with macrophage and innate lymphocyte subsets acquiring organ-specific transcriptional identities that underpin menstrual remodelling, ovarian follicle turnover, and placentation^37–39^. By jointly profiling prenatal and postnatal immune cells from the major reproductive organs, we resolved 54 cell states spanning the progenitor, lymphoid and myeloid lineages, uncovering tissue-specific subsets and transcriptomic signatures not described in previous single-organ studies **(Figure 3a** and **Extended Data Fig. 4a-b)**. The lower cellular representation of the cervix and vagina in our dataset precluded a comprehensive characterisation of immune subsets in these organs.

**Figure 3.**
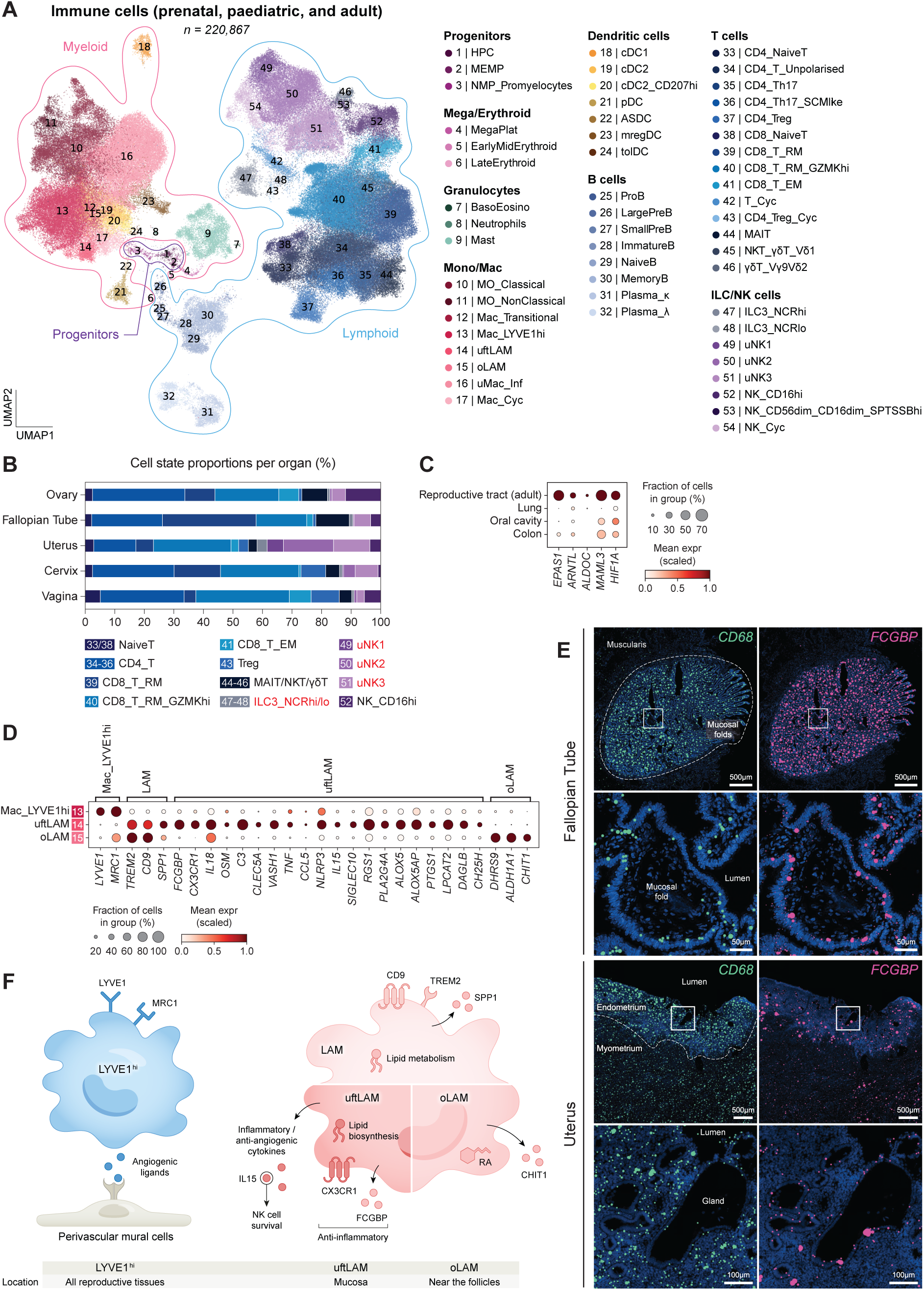
Immune cell profiling across reproductive organs reveals organ-specific innate immune subsets. **A,** Batch-corrected UMAP visualisation manifold of scRNA-seq data for all 220,867 immune cells from donors not taking exogenous hormones in the Human Female Reproductive System Cell Atlas v1, coloured by cell state. **B,** Bar plot displaying the proportion of select lymphocyte populations within each organ, excluding cycling, fetal, and menstrual fluid-derived cells. Cell states with names highlighted in red are enriched in the uterus. **C,** Dot plot showing the log-transformed, min-max normalised expression of selected genes upregulated by adult reproductive tract NCR^hi^ ILC3s compared to NCR^hi^ ILC3s derived from the lung^1,2^, oral cavity^3^, and colon^4^. *EPAS1*, *ARNTL*, and *ALDOC* were among the genes identified by differential gene expression analysis between reproductive tract- and non-reproductive tract-derived NCR^hi^ ILC3s. **D,** Dot plot showing the log-transformed, min-max normalised expression of selected marker genes (x-axis) for identified LYVE1^hi^ and LYVE1^lo^ macrophage subsets (y-axis) in the scRNA-seq data. **E,** (Top) Spatial distribution of transcripts for *CD68* (left column, pan-macrophage marker) and *FCGBP* (right column, uftLAM marker) within a fallopian tube cross-section stained by the Xenium 5k panel. Bottom two images are a zoom in of the inset indicated by the white square. Dotted shape indicates the boundary between the inner mucosal folds and the smooth muscle layer of the tube. (Bottom) visualisation of the same genes within an endometrial tissue section taken during day 5 of menstruation. Dotted line indicates the endometrial-myometrial border. Generated by 10x Genomics Xenium Explorer 4.1.1. **F,** Schematic summarising the potential signalling mechanisms of the LYVE1^hi^, uftLAM, and oLAM subsets, alongside their identified tissue location. HPC, haematopoietic progenitor cell; MEMP, megakaryocyte-erythroid-mast progenitor; NMP, neutrophil-myeloid progenitor; Baso, basophil; Eosino, eosinophil; MegaPlat, megakaryocyte/platelet; MO, monocyte; Mac, macrophage; uftLAM, uterine and fallopian tube lipid-associated macrophage; oLAM, ovary lipid-associated macrophage; Inf, inflammatory; Cyc, cycling; cDC, conventional dendritic cell; pDC, plasmacytoid dendritic cell; ASDC, AXL/SIGLEC6+ dendritic cell; mregDC, mature regulatory dendritic cell; tolDC, tolerizing dendritic cell; SCM, stem cell memory; Treg, regulatory T cell; MAIT, mucosal-associated invariant T cell; NK, natural killer; ILC3, type 3 innate lymphoid cell; RA, retinoic acid; expr, expression.

We identified a broad adaptive immune repertoire encompassing innate lymphocytes, T and B cell subsets previously undescribed in reproductive tissues **(Figure 3a, Extended Data Fig. 4b** and **Supplementary Note 2.1)**. Three previously described uterine natural killer (uNK) cell subsets^23^ were enriched in the uterus, as were type 3 innate lymphoid cells (ILC3s) (**Figure 3b**). We identified two ILC3 subsets that expressed *TOX2,* which is required for the metabolic adaptation to hypoxia in gut-resident ILC3^40^, but were distinguished on the basis of *NCR* expression, consistent with what has previously been described in the uterus^41^ (**Extended Data Fig. 4b**) Both subsets were localised to the functionalis layer of the endometrium (**Extended Data Fig. 5c**). When integrated with corresponding populations from the colon, lung, and oral cavity mucosa^42–45^ (**Extended Data Fig. 5a,b** and **Supplementary Note 1.2**), adult reproductive tract NCR^hi^ ILC3s exhibited differential expression of genes associated with hypoxia sensing and metabolic adaptation (*EPAS1*, *ARNTL*, *ALDOC*), suggesting an enhanced capacity to adapt to fluctuations in their local microenvironment (**Figure 3c** and **Supplementary Table 3**). Reproductive tract NCR^hi^ ILC3s also showed higher expression of *MAML3*, a co-activator of NOTCH signalling inducible under hypoxia^46^, and *HIF1A*, the key regulator of hypoxia-induced intestinal ILC3 responses in mice^47^. Notably, *HIF1A*, *ARNTL*, and *ALDOC* were reported to form a glycolysis-promoting transcriptional axis in colorectal cancer cells^48^, suggesting a similar axis may operate in ILC3s to support adaptation to local oxygen levels.

Within the myeloid compartment, we resolved conventional and non-conventional dendritic cell (DC) subsets, classical and non-classical monocytes, and a transitional macrophage population that co-expressed monocyte and macrophage markers **(Figure 3a, Extended Data Fig. 4b, Extended Data Fig. 5d** and **Supplementary Note 2.1)**. We also identified two broader macrophage subtypes distinguished by *LYVE1* expression **(Figure 3d)**. LYVE1^hi^ macrophages (called “*Mac_LYVE1hi*”) were detected across all reproductive organs **(Extended Data Fig. 4c)**, consistent with observations in mice^49,50^, and their transcriptomic profile pointed to a perivascular localisation and a non-inflammatory, tissue remodelling programme **(Extended Data Fig. 5e** and **Supplementary Note 2.1)**. By contrast, LYVE1^lo^ macrophages resolved into three organ-enriched subsets, two of which shared a canonical lipid-associated macrophage (LAM) transcriptional programme^51^ **(Figure 3d** and **Extended Data Fig. 5f),** and a population of inflammatory macrophages constrained to the menstruating endometrium (called “*uMac_Inf*”) (**Extended Data Fig. 5g)** (Lorenzi et al., biorxiv 2026). The two LAM populations were distinct between organs: uterine and fallopian tubes LAMs (uftLAMs) were defined by expression of *FCGBP* and *CX3CR1*, whereas ovary LAMs (oLAMs), which we described previously (Garcia-Alonso et al., biorxiv 2026), were confined to the ovary, exhibited a retinoic acid signature (e.g., *DHRS9 and ALDH1A1*), and upregulated *CHIT1* **(Figure 3d** and **Extended Data Fig. 4c)**.

Spatial transcriptomics localised uftLAMs to the mucosal epithelium in both the uterus (endometrium) and the fallopian tube **(Figure 3e),** situating them at a lipid-rich interface consistent with their lipid-associated signature. Compared with LYVE1^hi^ macrophages and oLAMs, uftLAMs are transcriptionally poised toward pro-inflammatory lipid mediator production, upregulating inflammatory genes (*IL18, OSM, C3, CLEC5A*) and anti-angiogenic mediators (*VASH1*) alongside *CX3CR1*, which is a hallmark of mucosal macrophage motility **(Figure 3d** and **Supplementary Table 3)**. Relative to oLAMs, uftLAMs display a markedly more inflammatory transcriptional state (*TNF, CCL5, NLRP3)* and distinctive upregulation of *IL15*, which activates uNK cells enriched in the endometrium **(Figure 3d** and **Supplementary Table 3).** However, concurrent expression of anti-inflammatory mediators (*FCGBP*, *SIGLEC10*, *RGS1*) places uftLAMs outside a canonical M1 classification **(Figure 3d)**, pointing instead to a tissue-adapted, mixed inflammatory state. Beyond lipid accumulation, which is characteristic of LAMs broadly, uftLAMs also deploy a dedicated biosynthetic programme: upregulation of *PLA2G4A*, *ALOX5*, *ALOX5AP*, *PTGS1*, *LPCAT2*, *DAGLB* and *CH25H* **(Figure 3d** and **Supplementary Table 3)** equips these cells to produce leukotrienes, prostaglandins, platelet-activating factor, and endocannabinoids, which are bioactive lipids that can act on neighbouring stromal and immune cells^52–56^.

Taken together, our cross-organ immune atlas reveals ILC3 and LAM subsets across reproductive organs, including a previously undescribed uftLAM population located in the reproductive mucosa **(Figure 3f)**.

### Mucinous epithelial specialisation across the reproductive tract

Epithelial and mesothelial cells constitute the structural lining of reproductive organs, forming continuous interfaces with the stromal compartments described above. Pan-organ integration showed that, despite organ-specific transcriptional differences, epithelial cells grouped into four sublineages shared across tissues: mucinous, squamous, ciliated and non-ciliated secretory (**Figure 4a,b** and **Supplementary Note 2.1**). The ovarian surface epithelium and the reproductive tract serosa separated from conventional epithelial subtypes, consistent with their mesothelial lineage.

**Figure 4.**
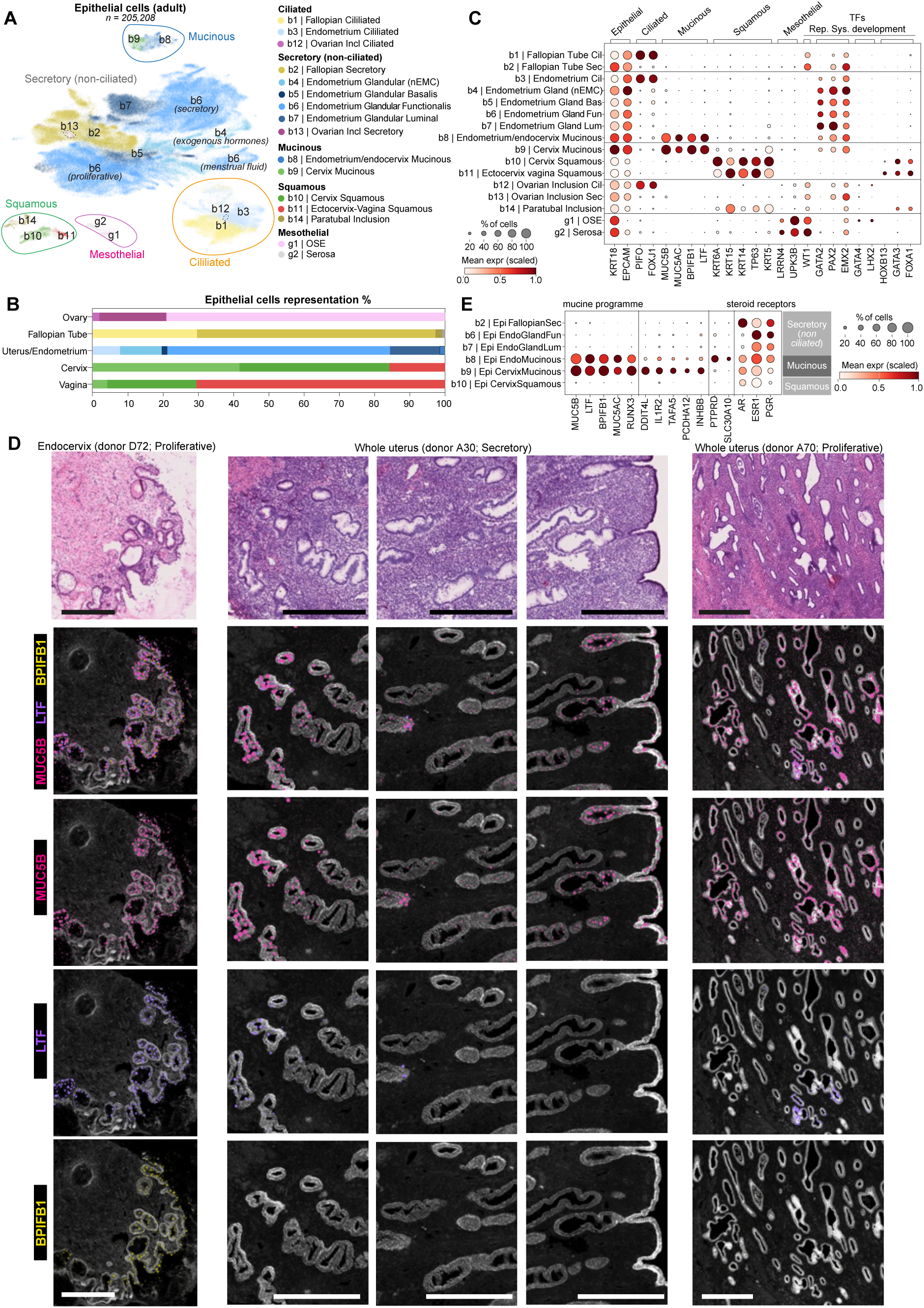
Epithelial lineage hierarchy across postnatal female reproductive organs. **A,** Batch-corrected UMAP visualisation manifold of the scRNA-seq dataset of all epithelial cells (n=205,208; "`broad_celltype`" labels) from the postnatal donors in the Human Female Reproductive System Cell Atlas v1, coloured by broad cell state. **B,** Bar plot showing the cellular composition of epithelial populations ("`broad_celltype`" labels) according to the sampled organ. **C,** Dot plot showing the log-transformed, min-max normalised expression of selected marker genes (x-axis) for the epithelial broad cell type categories identified in postnatal samples (y-axis). **D,** Visualisation of selected mucinous marker transcripts in representative whole endocervix and uterine sections profiled using a bespoke Xenium 480-gene panel. The panels show one endocervical section from donor D72 (reproductive age, proliferative phase) with a magnified view of the endocervical glands (left column); one whole uterine section from donor A30 (reproductive age, secretory phase) with three magnified regions showing MUC5B detection in the basalis, functionalis and lumen (middle columns); and one whole uterine section from donor A66 (reproductive age, proliferative phase) with one magnified region showing *MUC5B* detection in the basalis (right column). Colours indicate mucinous markers: *MUC5B* (pink), *LTF* (purple) and *BPIFB1* (yellow). Scale bars = 200 µm. Generated by 10x Genomics Xenium Explorer 4.1.1. **E,** Dot plot showing the log-transformed, min-max normalised expression of selected mucinous genes (x-axis) for the mucinous, non-ciliated secretory and squamous broad cell type epithelial categories (y-axis). Incl, inclusion; nEMC, non-endogenous menstrual cycle; OSE, ovarian surface epithelium; expr, expression.

Cross-organ integration resolved the identity of the *MUC5B*^+^ and the *AR*^hi^ epithelial population previously observed in endometrial datasets^19,22,57,58^ and represented in the HCA atlas by cells from multiple donors (**Extended Data Fig. 6a** and **Supplementary Note 2.2**). In the integrated atlas, these cells consistently aligned with the mucinous epithelial population of the cervix (**Figure 4a,b** and **Extended Data Fig. 6b**), an epithelium specialised in biochemical and immune barrier protection against ascending pathogens. Both endometrial and cervical mucinous cells expressed canonical mucinous markers (*MUC5AC*, *BPIFB1*, *LTF*) in addition to *MUC5B,* and overexpressed *AR*^58^ (**Figure 4c,e**). To determine the anatomical distribution of the endometrial mucinous populations, we interrogated high-resolution Xenium images.

This revealed that the full mucinous programme (*MUC5B*^+^/*BPIFB1*^+^/*LTF*^+^) was confined to the endocervix (**Figure 4d**, first column, and **Extended Data Fig. 6c**). Xenium imaging also showed that a small subset of endometrial glands, predominantly in the basalis but also in the lumen, expressed mucinous-associated markers (*MUC5B* and *LTF*) but lacked other mucinous markers (*BPIFB1*), indicating partial activation of the mucinous programme by endometrial glands (**Figure 4d**). Comparing cervical and endocervix mucinous cells revealed differences in the expression of cell adhesion regulators (*TAFA5, PCDHA12, PTPRD*), ion transport (*SLC30A10*) and cell-cell signalling (*INHBB*, *IL1R2*), suggesting tissue-specific mucinous specialisation between endocervix and cervix (**Figure 4e**).

Altogether this suggests that the mucinous programme is not restricted to the cervix barrier specialisation, but extends across reproductive epithelia with tissue-specific states: a full programme in the endocervix and cervix (*MUC5B*^+^/*BPIFB1*^+^/*LTF*^+^/*MUC5AC*^+^), and a partial programme (*MUC5B*^+^/*LTF*^+^) activated within a subset of glandular epithelial cells in the endometrium.

### Unconventional epithelial states sampled in reference tissues

The strong tissue-specific identities of reproductive epithelial populations (**Figure 4a** and **Extended Data Fig. 6b**) allowed us to identify rare epithelial cells whose transcriptional programmes matched that of different reproductive organs. We identified three epithelial populations outside their usual anatomical location: two ovarian cortical epithelial inclusions, one of which was compatible with focal ovarian endometriosis, and one paratubal inclusion consistent with Walthard cell rests.

In ovarian samples, beyond the mesothelial cells of the ovarian surface epithelium, we detected rare *EPCAM*^+^/*PAX8*^+^ epithelial cells annotated as “*Ovarian Inclusion Ciliated*” and “*Ovarian Inclusion Secretory*” (n = 202 and n = 20 cells, respectively; **Figure 4b**), which mapped to reproductive tract epithelial states (**Figure 4a**). Spatial transcriptomics localised these cells to discrete intraovarian inclusions, supporting their interpretation as tissue-resident epithelial populations rather than sampling contaminants. One structure contained *PAX8*^+^/*OVGP1*^+^ secretory cells, consistent with an ovarian cortical inclusion cyst lined by tubal-type epithelium, long proposed as a site of origin of epithelial ovarian carcinoma^59^ (**Figure 5a,b**). A second structure contained *PAX8*^+^/*WNT7A*^+^/*OVGP1*^−^ endometrial-like epithelium, surrounded by *HOXA9*^+^/*HOXA10*^+^ endometrial-type stroma (**Figure 5c,d**), compatible with focal endometriosis in the ovary.

**Figure 5.**
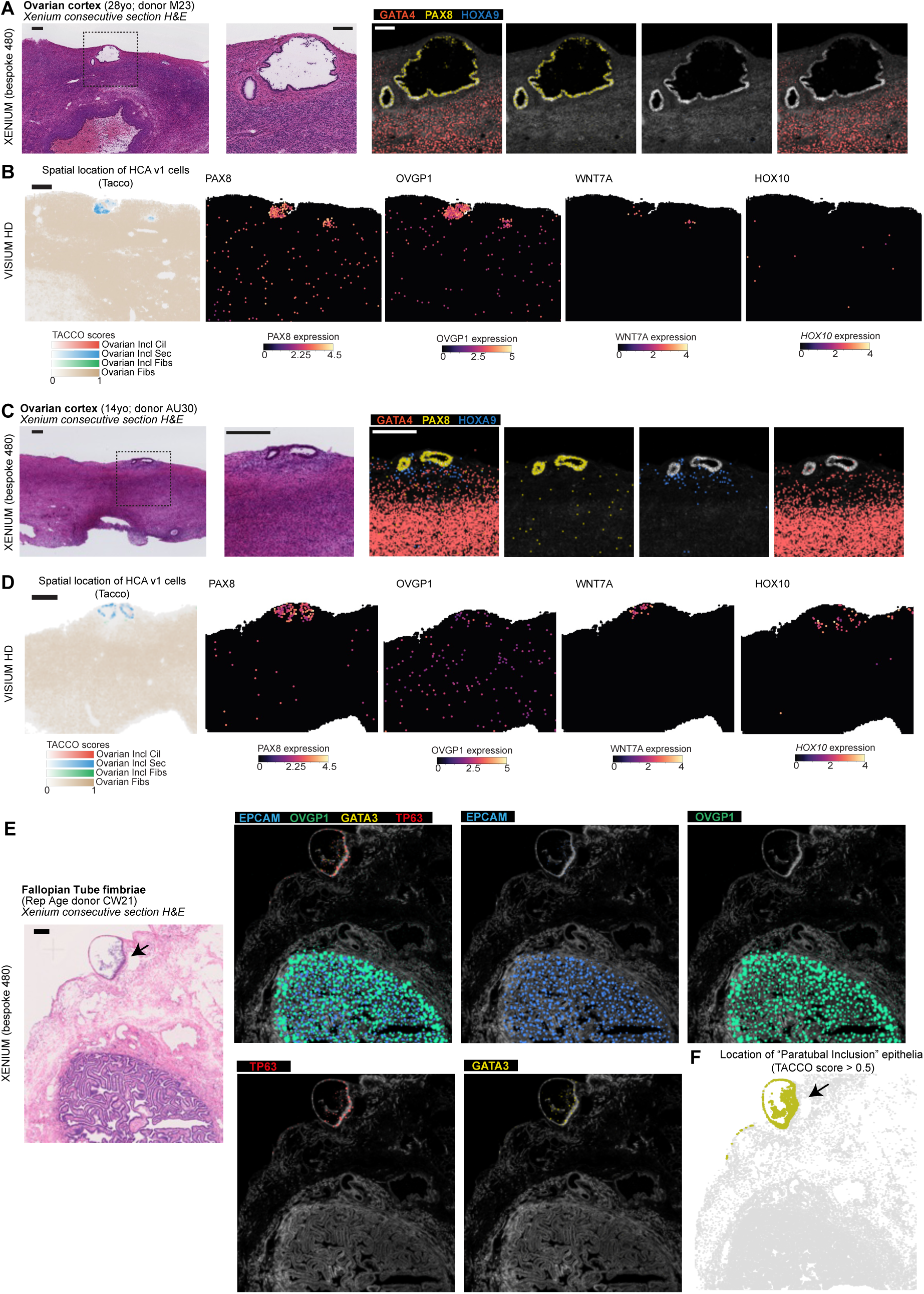
Unconventional epithelial states sampled in reference tissues. **A.** Visualisation of selected marker transcripts in a representative ovarian cortex section from donor M23 (28 years old), containing an ovarian inclusion cyst and profiled using a bespoke Xenium 480-gene panel. The left panels show the corresponding haematoxylin and eosin (H&E) section and a magnified view of the inset indicated by the black dashed square. Colours indicate *GATA4* (red; ovarian interstitial fibroblasts), *PAX8* (yellow; reproductive tract epithelium) and *HOXA9* (blue; lumbosacral HOX factor characteristic of the uterine segment). Scale bar = 200 µm. Images were generated using 10x Genomics Xenium Explorer 4.1.1. **B.** Visualisation of selected marker transcripts in an ovarian cortex section adjacent to that shown in A, profiled using Visium HD. The left panels show broad cell-type labels transferred from the Human Female Reproductive System Cell Atlas v1 onto Visium HD 8 µm bins using the TACCO tool. Gene panels show log-transformed expression of *PAX8* (reproductive tract epithelium), *OVGP1* (fallopian tube-like epithelium), *WNT7A* (endometrial-like epithelium) and *HOXA10* (lumbosacral HOX factor characteristic of the uterine segment). Scale bar = 200 µm. **C.** Visualisation of selected marker transcripts as in A, in an ovarian cortex section from donor AU30 (14 years old) containing a lesion histologically consistent with an ovarian endometrioma. Scale bar = 200 µm. Generated by 10x Genomics Xenium Explorer 4.1.1. **D.** Visualisation of selected marker transcripts as in B, in an ovarian cortex section adjacent to that shown in C, profiled using Visium HD. Scale bar = 200 µm. **E.** Visualisation of selected marker transcripts in a representative fallopian tube section from donor CW21 (fimbriae; reproductive age), containing a paratubal inclusion consistent with Walthard cell rests and profiled using a bespoke Xenium 480-gene panel. The left panel shows the corresponding haematoxylin and eosin (H&E) section. Colours indicate *EPCAM* (blue; canonical epithelial marker), *OVGP1* (green; fallopian tube epithelium), *GATA3* (yellow; urothelial epithelium) and *TP63* (red; squamous epithelium). Scale bar = 200 µm. Generated by 10x Genomics Xenium Explorer 4.1.1. **F.** Visualisation of predicted paratubal inclusion epithelial cells in the same Xenium section shown in E. Cell labels from the Human Female Reproductive System Cell Atlas v1, including the paratubal inclusion epithelial label, were transferred onto Xenium data using the TACCO tool. Incl, inclusion; Cil, ciliated; Sec, secretory; Fib, fibroblast.

The same principle extended beyond the ovary. We detected squamous/basal *KRT5*^+^/*GATA3*^+^ cells in fallopian tube samples from 9 of 22 donors, which localised to paratubal inclusions consistent with Walthard cell rests, a common benign histopathological finding with squamous/urothelial-like features historically described as transitional epithelium^60^ (**Figure 5e**). These interpretations were reviewed with expert histopathologists, who supported the diagnosis of Walthard cell rests, ovarian cortical inclusion cysts, and focal ovarian endometriosis in the corresponding regions.

Altogether, our pan-reproductive atlas allowed rare epithelial populations to be interpreted not only by their anatomical location, but by the epithelial programme they carried. These findings highlight the value of large-scale cross-tissue atlases for detecting microscopic abnormalities in control tissues that may represent early or preclinical stages of disease.

### Spatiotemporal dynamics of mesenchymal cells contextualise genetic risk for gynaecological conditions

The cell type resolution of our atlas provides a resource for interpreting the genetic architecture of common reproductive conditions at cellular and temporal precision. The highly polygenic nature of these conditions complicates identification of pathogenetic mechanisms, yet offers an opportunity: by harnessing the aggregate signal across many loci, we can begin to disentangle the core cell types and pathways underlying disease. We integrated genome-wide association study (GWAS) signals for major gynaecological conditions^61–68^ **(Supplementary Table 4a)** with cell-type-specific expression profiles from our HCA atlas, asking whether higher gene expression specificity per cell type was associated with disease association within and near each gene using LD score regression^69^. We followed up through leave-one-out gene decomposition to identify the pathways driving each association **(Figure 6a, Extended Data Fig. 7a-b, Supplementary Note 2.3, Supplementary Tables 4b-d)**.

**Figure 6.**
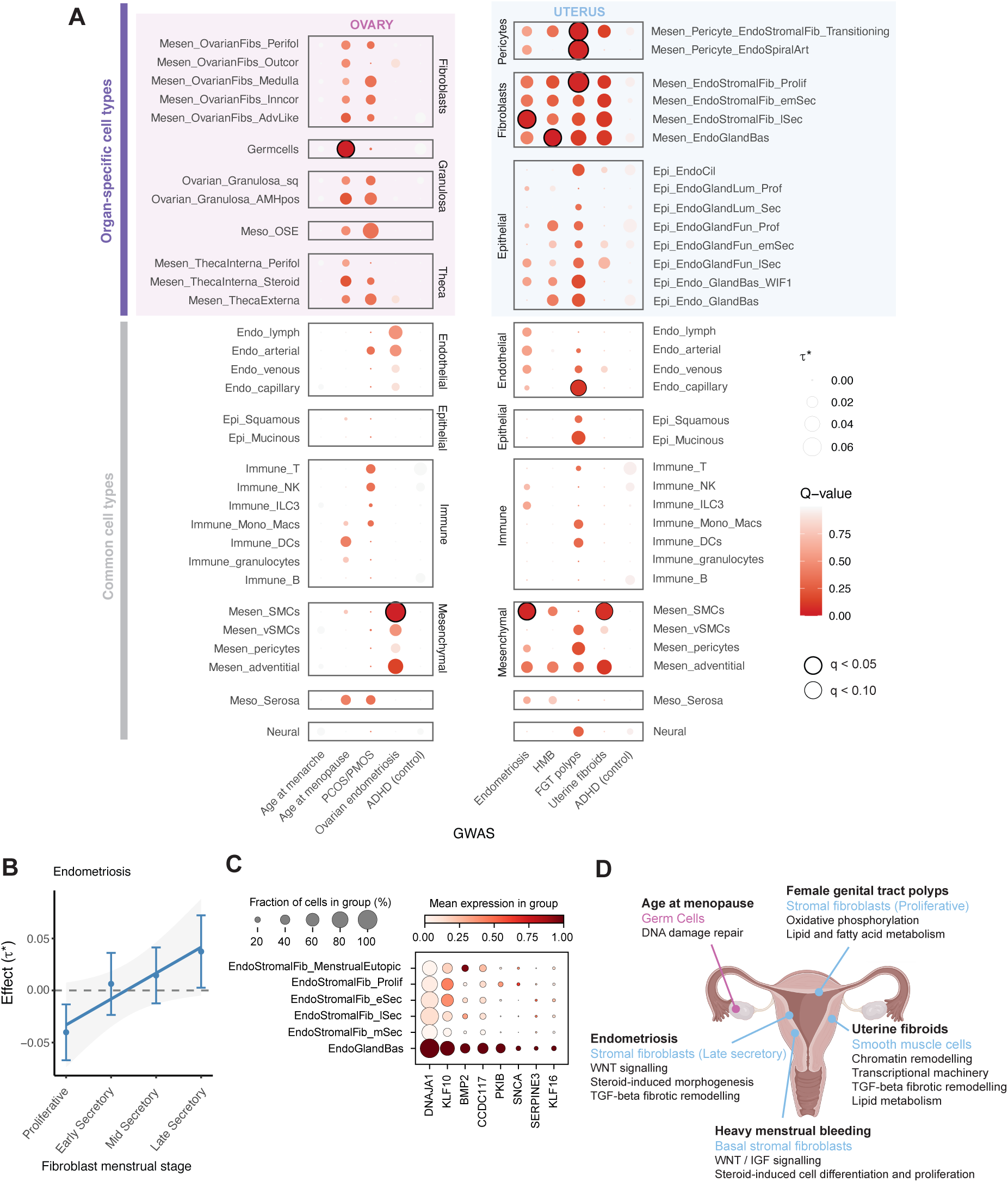
Integrating GWAS with the Human Female Reproductive System Cell Atlas v1. **A,** Cell type enrichment of disease heritability in the specifically expressed genes per cell type. The dot plots show enrichment for both uterine and ovary specific traits (x-axis) computed using LDSC, with ovary-specific and uterus-specific cell types respectively highlighted in pink and blue and non-specific cell types not highlighted. Cell types (y-axis) are grouped by lineage and enrichment is shown in dots where the size represents the standardised effect **τ**\* – the additive change in heritability explained by a 1 standard deviation increase in the gene expression specificity (CELLEX) annotation – and the colour represents significance q-value as -log10(q), corrected for multiple testing. Bold black circles indicate enrichment at q<0.05 and light black circles indicate enrichment at q<0.10. Data points with **τ**\*<0 were set to 0 and corresponding q-values set to 1. **B,** Endometriosis enrichment in endometrial fibroblasts across the menstrual cycle. The plot shows the within-stromal fibroblast enrichment effect **τ**\* and corresponding standard error (SE) for endometriosis in fibroblasts ranked by menstrual cycle phase. The line is the weighted least squares association between cell state rank and **τ**\*, weighted by **τ**\* SE (β=0.0249, p=0.045). **C,** Dot plot of normalised, log-transformed expression of genes driving basal stromal fibroblast enrichment for heavy menstrual bleeding (HMB) heritability across endometrial stromal fibroblasts. Here we show genes that have high gene expression specificity scores (CELLEX) for basal fibroblasts computed against the other fibroblasts shown. The size of the dot represents the number of cells in the cell type (y-axis) expressing each gene (x-axis) and the colour represents the gene expression. **D,** Summary of results from cell type enrichment and follow-up analyses, with each trait in bold, the top cell type in blue (uterus) or pink (ovary) and the associated pathways annotated below. Mesen, mesenchymal; Epi; epithelial; Endo, endothelial/endometrial; Meso, mesothelial; Fib, fibroblast; Perifol, perifollicular; Outcor, outer cortex; Inncor, inner cortex; Adv, adventitial; sq, squamous; OSE, ovarian surface epithelium; Art, arteries; Prof/Prolif, proliferative; Sec, secretory; emSec, early/mid secretory; lSec, late secretory; Cil, ciliated; Lum, luminal; Bas, basalis; NK, natural killer; ILC3, type 3 innate lymphoid cell; Mono, monocyte; Mac, macrophage; DC, dendritic cell; SMC, smooth muscle cell; vSMC, vascular smooth muscle cell; PCOS, polycystic ovary syndrome; PMOS, polyendocrine metabolic ovarian syndrome; ADHD, attention deficit hyperactivity disorder; HMB, heavy menstrual bleeding; FGT, female genital tract.

Genetic risk for age at menopause was strongly associated with germ cells in paediatric and adult ovaries, consistent with the central role of the oocyte pool in determining reproductive lifespan **(Figure 6a, Supplementary Note 2.3, Supplementary Table 4b)**. Analyses of the prenatal subsets further identified enrichment across germ cells and the ovarian somatic precursor populations **(Extended Data Fig. 7a, Supplementary Table 4b)**, pointing to the coupled differentiation of germ cells and somatic progenitors during primordial follicle establishment and illustrating the utility of our cross-lifespan atlas for identifying cell types relevant to both prenatal and adult traits. In the adult endometrium, mesenchymal cells were broadly enriched across multiple uterine pathologies, including endometriosis, heavy menstrual bleeding (HMB), female genital tract (FGT) polyps and uterine fibroids **(Figure 6a, Supplementary Table 4b)**. Despite the ovary being a major site of endometriosis lesions, ovary-specific cell types were not enriched for ovarian endometriosis, in line with with recent work using somatic mutations to suggest that the cells in endometriosis lesions are derived from the endometrium^70^.

Within endometrial stromal fibroblasts, different conditions implicate cell states defined not only by transcriptional identity but by menstrual cycle stage and spatial location **(Figure 6a, Supplementary Table 4b)**. Endometriosis heritability was enriched in late secretory stromal fibroblasts and smooth muscle cells **(Figure 6a)**. Given its pattern of enrichment in fibroblasts, we performed higher resolution enrichment analyses within endometrial stromal fibroblasts and determined that endometriosis risk enrichment increased as a function of menstrual stage **(Figure 6b, Supplementary Table 4c)**. Enrichment in late secretory stromal fibroblasts was driven by WNT signalling (*WNT4*) and TGF-β-driven fibrotic remodelling (*TGFB1I1*) **(Supplementary Table 4d, Extended Data Fig. 7b)**, which has been well-characterised in endometriosis pathogenesis^71^. FGT polyps heritability was significantly enriched in proliferative-phase endometrial stromal fibroblasts, pericyte-like transitioning stromal fibroblasts, spiral artery pericytes and capillary endothelial cells **(Figure 6a)**, illustrating the importance of characterising tissue-specific cell populations. This is consistent with single-cell transcriptomic studies of endometrial polyps which have described altered perivascular, stromal, epithelial and endothelial crosstalk^66,72^. Genes driving enrichment in proliferative-phase endometrial stromal fibroblasts pointed to altered lipid and fatty acid metabolism (*ZFYVE1*, *PLA2G4C*, *APOB*, *CTHRC1)* and mitochondrial (*SIRT3*, *NDUFS1, TMEM135*^73^), consistent with altered metabolic activity in polyp-associated fibroblasts **(Extended Data Fig. 7b, Supplementary Table 4d)**.

We identify, for the first time, a population of endometrial fibroblasts co-localising adjacent to the basalis glandular epithelium (annotated as “Mesen_EndoGlandBas”) that is significantly enriched for genetic risk of HMB, a multifactorial condition characterised by excessive blood loss during menstruation **(Figure 6a, Supplementary Table 4b)**. Whilst multiple conditions can cause HMB, many cases have no known cause identified^74^ and the convergence of genetic risk in basal fibroblasts suggests a common, dominant or convergent mechanism in the HMB GWAS cohort. These fibroblasts reside in the basalis layer, the regenerative compartment of the endometrium that is retained after menstrual shedding and drives tissue renewal each cycle (Lorenzi et al. biorxiv 2026). Their enrichment for HMB heritability was driven by genes involved in intercellular signalling and tissue remodelling, including pathways identified in the HMB GWAS such as WNT (*WNT4*, *WNT11)* and IGF pathways (*IGF1*, *IGFBP3*, *IGF1R*, *IGFBP2* and *IGFBP6*) and re-epithelialisation factors (*FGF1*, *PRDM1*, *BMP2)*^75^ **(Extended Data Fig. 7b, Supplementary Table 4d)** alongside genes specific to these basal fibroblasts, including *DNAJA1*, *KLF10*, *BMP2* and *CCDC117*, all of which had HMB risk loci at *p*<1e-5 in or near the gene body **(Figure 6c, Extended Data Fig. 7c)**. These point to two converging mechanisms: steroid-responsive regulation of cell proliferation through the *BMP2*–*KLF10* axis^76^ and *CCDC117*^77^ as well as control of matrix metalloproteinase activity though *DNAJA1*^78^ which could alter tissue breakdown in the basalis **(Supplementary Note 2.3)**. Together, these signals suggest that dysregulation of the regenerative niche in basal fibroblasts, through impaired proliferative control or excessive matrix degradation, may contribute to the pathogenesis of HMB.

These analyses establish a spatiotemporally resolved map of genetic risk across gynaecological conditions, identifying cell states and cycle stages through which disease-associated variants are likely to act and providing a framework for prioritising candidate genes and pathways underlying cell type associations **(Figure 6d)**.

### A chromatin accessibility atlas links non-coding variants to cell-type specific regulatory elements

While transcriptomic profiles identify disease-enriched cell types at the polygenic level, nominating the specific genes regulated by non-coding variants requires direct characterisation of the accessible regulatory landscape. To provide this, we assembled a harmonised scATAC-seq atlas across the ovary, fallopian tube and endometrium, supplementing existing data with newly generated profiles covering the proliferative phase of the menstrual cycle, which was underrepresented in prior datasets. ScATAC-seq data only covered the functional layer of the endometrium present in Pipelle biopsies, precluding the study of basal cell types which we showed are relevant for HMB. The resulting atlas comprises 112,798 high quality cells across 43 donors **(Figure 7a-c)**, with identities transferred from our transcriptomic atlas, and faithfully recapitulates subtle cell-state changes across the menstrual cycle **(Extended Data Fig. 8a-d** and **Supplementary Note 2.4)**

**Figure 7.**
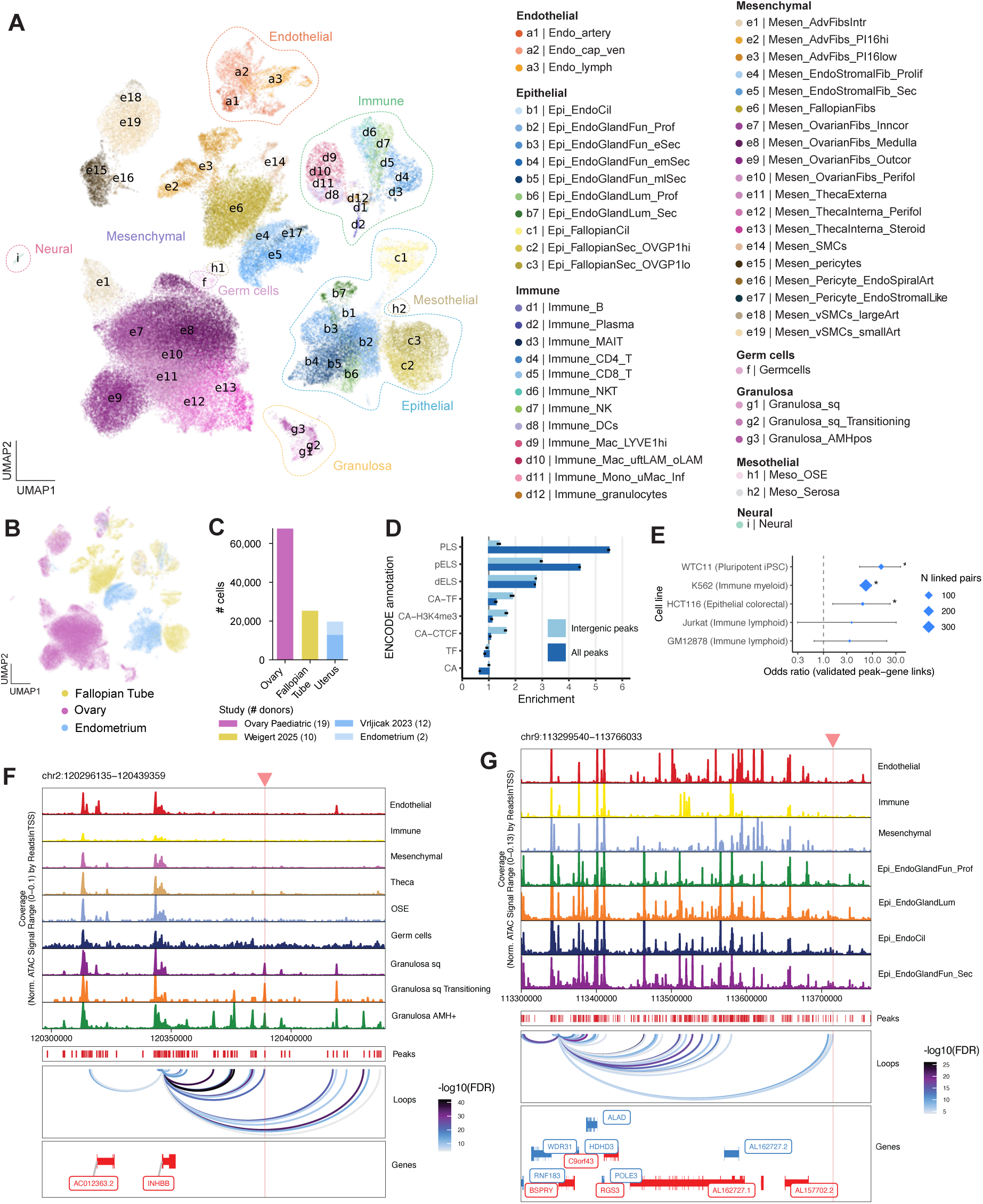
Single-cell chromatin accessibility atlas identifies regulatory relationships. **A,** Batch-corrected UMAP visualisation manifold of the cells in the scATAC-seq atlas, coloured by cell type and grouped by lineage. **B,** Same UMAP manifold, coloured by contributing organ. **C,** Bar plot of the number of cells per organ, coloured by dataset with the number of contributing donors in the legend. “Ovary Paediatric” is from (Garcia-Alonso et al., under review) and “Endometrium” is our in-house generated endometrium scATAC-seq data. **D,** Enrichment of open chromatin peaks for ENCODE functional annotations. Enrichment for all peaks is shown in dark blue and enrichment for intergenic peaks (removing peaks overlapping a gene body) is shown in light blue. The annotations are promoter-like signatures (PLS), proximal enhancer-like signatures (pELS), distal enhancer-like signatures (dELS), the intersection of ENCODE chromatin accessibility and predicted transcription factor binding sites (CA-TF), the intersection of ENCODE chromatin accessibility and H3K4me3 sites (CA-H3K4me3), the intersection of ENCODE chromatin accessibility and CTCF sites (CA-CTCF), predicted transcription factor binding sites (TF) and ENCODE chromatin accessibility (CA). All enrichments (>0) and depletions (<0) shown were significant after multiple testing except the intergenic peak enrichment for ENCODE chromatin accessibility. **E,** Enrichment of peak–gene links for experimentally validated enhancer–gene links. Enrichment was computed for enhancer–gene links validated from multiple cell lines (y-axis) and the odds ratio shown on the x-axis, with significance indicated by an asterisk. The diamond size represents the number of peak–gene links recapitulated in each dataset. **F,** Chromatin accessibility track plot for peaks associated with *INHBB* expression. Chromatin accessibility is shown in different colours for different cell populations as the fragment coverage across the window normalised by total number of reads in transcription start sites. 500bp peaks are shown in red in the “Peaks” track, and peak-gene links are shown in the “Loops” track, coloured by -log10 FDR of the association. Genes from the window are shown in the bottom track, including the target gene. The red arrow highlighted region pointed to by the red triangle is a peak overlapping a PCOS disease risk locus index SNP. **G,** Chromatin accessibility track plot for peaks associated with *BSPRY* expression. Plot structure is the same as **F**, with the highlighted peak containing a female genital tract disease locus index SNP. Endo, endothelial/endometrial; Epi, epithelial; Mesen, mesenchymal; Meso, mesothelial; Cil, ciliated; Fun, functionalis; Lum, luminal; Prof/Prolif, proliferative; Sec, secretory; eSec, early secretory; emSec, early/mid secretory; mlSec, mid/late secretory; MAIT, mucosal-associated invariant T cell; NKT, natural killer T cell; NK, natural killer; DC, dendritic cell; Mac, macrophage; uftLAM, uterine and fallopian tube lipid-associated macrophage; oLAM, ovary lipid-associated macrophage; Inf, inflammatory; Adv, adventitial; Fib, fibroblast; Intr, interstitial; Inncor, inner cortex; Outcor, outer cortex; Perifol, perifollicular; SMC, smooth muscle cell; Art, arteries; vSMC, vascular smooth muscle cell; sq, squamous; OSE, ovarian surface epithelium.

We identified 718,670 reproducible accessible chromatin peaks contributed by 50 cell types **(Extended Data Fig. 9a)**, strongly enriched for regulatory annotations from the ENCODE project, particularly promoter-like signatures (PLS), proximal enhancer-like signatures (pELS) and distal enhancer-like signatures (dELS), confirming that they capture genuine regulatory elements **(Figure 7d)**. We computed peak-to-gene links connecting accessible chromatin peaks to putative target genes **(Extended Data Fig. 9b,c)**, retaining 322,981 high-confidence links after stringent filtering. These links showed significant enrichment for experimentally confirmed enhancer-gene pairs across multiple cell lines (OR=6.3-15.1), highest in induced pluripotent stem cells (WTC11) and lowest in lymphoid lines (Jurkat, GM12878), consistent with the limited lymphoid representation in our dataset **(Figure 6e)**. Peaks linked to a gene were also significantly enriched for ENCODE functional annotations compared with unlinked peaks **(Extended Data Fig. 9d)**. Together, these analyses demonstrate that our peak–gene links reflect genuine regulatory relationships.

To nominate disease effector genes, we overlapped linked peaks with GWAS loci for major gynaecological conditions, identifying 290 peaks associated with 36 disorders and focusing on those pointing unambiguously to a single effector gene **(Supplementary Table 5a)**. The approach recovered established disease genes, including *NGF* in endometriosis^65^, *WT1*^68^ and *PLSCR3*^79^ in uterine fibroids and *NEBL* in epithelial ovarian tumours^80^. Among novel findings, we provide the first regulatory evidence linking a PMOS (formerly PCOS^81^) risk locus (lead SNP in peak chr2:120388859-120389359) to *INHBB* expression **(Figure 7f)**. *INHBB* encodes Inhibin B, a glycoprotein that suppresses follicle-stimulating hormone (FSH) in a negative feedback loop and promotes androgen release from ovarian theca cells^82^. Although *INHBB* has been proposed as a PMOS candidate gene on the basis of proximity to the risk signal^63,64^, it has not been prioritised due to the absence of functional regulatory evidence. Here we show that a PMOS-associated chromatin peak is linked to *INHBB* expression, preferentially open in transitioning and AMH^+^ granulosa cells **(Figure 7f)** which were the second most enriched cell type for PMOS heritability **(Figure 6a)**. *INHBB* has primarily been studied as a biomarker for folliculogenesis and IVF outcome^83^ but is elevated in the peripheral blood of PMOS patients^84^; our data provide the first regulatory evidence for its direct involvement in PMOS and support it as a candidate therapeutic target.

We additionally identify *BSPRY* as a novel candidate effector gene for an FGT polyps risk locus^85^, with the associated peak overlapping a putative enhancer 394,800bp downstream of the gene, preferentially accessible in ciliated and glandular secretory epithelial cells **(Figure 7g)**. *BSPRY* has been implicated in calcium signalling in epithelial tumours^86^. Although its precise mechanistic role in polyp formation remains to be established, the convergence of genetic and regulatory evidence positions it as a strong candidate for further investigation.

Together, this chromatin accessibility atlas complements our transcriptomic reference, providing a cell-type-resolved regulatory map that bridges genetic associations and biological mechanisms, and identifying potential effector genes across a spectrum of gynaecological conditions.

## Discussion

We present the *Human Female Reproductive System Cell Atlas v1*, integrating data from more than two million cells across 291 donors and 27 datasets, spanning the ovary, fallopian tube, uterus, cervix and vagina, to define 210 cell types and states through community consensus annotation. The atlas accounts for the spatial organisation of the distinct organs and temporal dynamics at two scales: the lifespan, from fetal development through menopause, and the menstrual cycle, driven by monthly hormonal fluctuations. Beyond serving as a reference for shared and organ-specific cellular programmes, the atlas makes three conceptually distinct contributions: it defines the tissue-resident cellular landscape across reproductive organs at a resolution inaccessible to single-organ studies; it demonstrates that large-cohort, cross-tissue references can detect structures in otherwise healthy donors that represent early or preclinical disease signatures; and it provides a framework for translating genetic disease associations into the specific cell types and hormonal contexts in which they act.

Cross-organ integration of the mesenchymal compartment revealed a conserved hierarchy of non-interstitial fibroblasts – adventitial fibroblasts, perivascular mural cells and smooth muscle cells – consistent with the universal fibroblast architecture described in mice^31^ and extending it to the human reproductive system. Unlike interstitial fibroblasts, which are highly organ-specific and temporally regulated by hormonal cycling, this non-interstitial compartment is relatively stable across the menstrual cycle, suggesting a structural rather than hormone-responsive role. While the core mural compartment is broadly conserved across the body^27,87^, organ-specific perivascular populations have been described in tissues with specialised vascular demands, including the brain^88^ and lung^89^. The uterus represents a further example of this, harbouring two perivascular populations absent from other organs. Spiral artery-associated pericytes express *FLT1*, which encodes the primary VEGF receptor and a key regulator of endometrial angiogenesis^90^, and *STC2*, a hypoxia-inducible survival factor, suggesting these cells are molecularly adapted to support spiral artery remodelling and to persist within the hypoxic environment of menstruation^91^. The endometrial perivascular-stromal population, co-expressing pericyte and stromal markers, is consistent with a proposed vascular niche for endometrial stromal progenitors. Collectively, the definition of this non-interstitial compartment in healthy reproductive tissues provides a reference for pathological processes including endometriosis, pelvic adhesions and fibrotic disease of the reproductive organs, in which adventitial and perivascular cells are thought to play a central but poorly understood role.

The immune subsets of the integrated atlas reveal the depth of tissue-resident immune specialisation across reproductive organs. Uterine ILC3s had been identified in prior studies^19,41,92^, but we now localise them to the functionalis layer of the endometrium and, through integration with ILC3s from other organs, resolve their distinctive tissue-resident programme. We find two ILC3 subsets distinguished by NCR expression, where the NCR^hi^ subset upregulates genes indicative of adaptation to the hypoxic uterine environment and shows elevated NOTCH signalling, a pathway independently linked to hypoxia^46^, which may promote the acquisition of mature NCR^hi^ ILC3 identity in the reproductive tract, consistent with findings in the mouse intestine^93^. In line with the role of ILC3 in tissue regeneration and recovery^94^ and their peak in abundance during endometrial repair^92^, our results suggest that NCR^hi^ ILC3 represent a locally adapted mucosal population that may contribute to tissue homeostasis within the endometrium, particularly during stages of transient hypoxia such as menstruation.

Within the myeloid compartment, macrophages divide into LYVE1^hi^ and LAMs across all reproductive organs, consistent with their distribution in non-reproductive tissues^25,49^. Cross-organ integration further resolves organ-specific LAM subsets, indicating that reproductive tissues may establish distinct immune niches tailored to the physiological demands of each organ. Amongst them, uftLAMs, shared between the uterine endometrium and fallopian tube, upregulate *IL15*, which can activate the unique uNK cells characteristic of the endometrial niche^95^, and co-express inflammatory and immunoregulatory markers, including *FCGBP*, reflecting the dual requirement for immune vigilance and tolerance that characterises the uterine microenvironment during pregnancy^96^. LAMs have emerged as pathologically relevant in multiple tissues^97^, and how uftLAMs modulate inflammatory reproductive disorders such as endometriosis is an important open question.

Cross-organ integration also clarified the identity of a *MUC5B*^+^/*AR*^hi^ population repeatedly observed in endometrial datasets^19,22,57,58^. In the integrated atlas, this *MUC5B*^+^ population aligns with the mucinous epithelium of the cervix, sharing some features of a goblet-like mucosal barrier programme seen in tissues such as the lung airway and intestine^98^. Spatial imaging confirms that the full mucinous programme is restricted to the endocervix and cervix, and that endometrial glands activate only a partial version marked by *MUC5B* upregulation. Accordingly, *MUC5B*^+^ cells recovered from endometrial datasets most likely represent endocervical cells captured during tissue sampling, a misidentification resolvable only through integrated spatial and transcriptomic analysis; we cannot exclude, however, that this population is genuinely present at low frequency in the endometrium of a subset of donors. The mucinous programme (*MUC5B*^+^/*LTF*^+^) has been reported in other barrier tissues^98^, and reflects the specialised defence function of the cervix against ascending infection. The programme has also been reported in endometriosis^22,99^ and aligns with the epithelial cells reported in some endometrial cancers^100,101^, underscoring the importance of resolving this mucinous programme and its disease relevance.

A key conceptual contribution of this work is that large-cohort cross-tissue references can reveal the earliest cellular traces of disease in tissues collected from control individuals. The pan-organ design of the atlas provided the transcriptomic reference needed to recognise rare epithelial cells in ovarian and fallopian tube samples whose identity pointed to other organs. These ectopic or lesion-associated structures would be difficult to interpret in single-organ single-cell datasets and may escape histological detection if small or not captured in the section imaged. Here we captured a rare population compatible with an early endometriosis lesion in a 14 years old donor, potentially representing one of the earliest events in endometrioma development, before pathology is clinically established and before the initiating cell states and their surrounding niche have evolved into a progressed pathology. We also captured ovarian cortical inclusion cysts of tubal origin in five donors, which have historically been considered possible precursors of epithelial ovarian carcinoma^29,59^. Disease-focused atlases profile tissues once disease is clinically apparent or advanced stages, when the early cellular and microenvironmental context of initiation has already evolved. By contrast, large-scale control cohorts offer a unique window into these early events, arguing that profiling control tissues may offer a clearer view to investigate how gynaecological pathologies emerge at the cellular level.

Integration with GWAS data reveals that genetic risk for uterine conditions acts predominantly through the mesenchymal compartment, and that the implicated cell states are defined not only by transcriptional identity but by menstrual cycle stage. This spatiotemporal dimension underscores a fundamental challenge in interpreting reproductive disease genetics: studies that do not account for tissue dynamics will systematically miss or misassign the cellular context of risk variants. The atlas was essential to this resolution, providing organ-specific populations such as spiral artery pericytes, implicated in FGT polyps, and rare populations such as basal fibroblasts, implicated in HMB, neither of which had been characterised before. HMB affects approximately one in three women^102,103^, a prevalence exceeding that of asthma^104^, yet remains massively underdiagnosed and undertreated. The convergence of HMB genetic risk identified in basal fibroblasts, made possible by our parallel characterisation of the basalis layer (Lorenzi et al., biorxiv 2026), illustrates how atlas-guided cell-type resolution can generate physiologically grounded, genetically informed mechanistic hypotheses and nominate new avenues for therapeutic intervention.

Because most disease-associated variants lie in non-coding regions, our ability to nominate individual disease effector genes is limited in transcriptomics-based approaches. Our chromatin accessibility atlas provides peak-to-gene linkage maps that connect non-coding variants to target genes at cell type resolution. A key limitation is that the chromatin atlas represents the uterus through endometrial biopsies only, precluding capture of basal cell types relevant to HMB; full-thickness uterine sampling in future versions would substantially extend the utility of these maps. Despite this, the resource enables effector gene nomination at scale, exemplified by the regulatory link between a PMOS risk locus enhancer and *INHBB* expression in granulosa cells, providing the first functional evidence for a nomination previously based on genomic proximity alone. The translational value of this approach is substantial: therapeutic targets supported by GWAS evidence are more than twice as likely to succeed in clinical trials^105^.

Several limitations of our study should be noted. First, access to raw sequencing data was required to reprocess all datasets through a common pipeline and minimise technical variation. This is particularly important in reproductive tissues, where hormone-responsive states introduce substantial biological variation between donors. Together with the limited availability of deeply profiled cervix and vagina datasets, this contributed to the underrepresentation of these tissues in the current atlas^106^. Secondly, metadata completeness varied across studies, especially for the menstrual cycle stage, which was often unavailable in datasets not focused on the uterus. Pregnancy was excluded because it represents a major physiological state that would introduce additional cellular and temporal complexity beyond the scope of this non-pregnant reproductive system reference. Prenatal donors are included in the atlas and their data are available for community exploration — for instance, to trace developmental origins of adult cell types or examine whether disease-associated transcriptional states reflect a return to earlier developmental programmes. For uterus, we retained eutopic uterine samples from donors with endometriosis, a condition defined by the presence of endometrial-like tissue outside the uterus, because prior studies reported no major cell type changes or broad transcriptional shifts in the eutopic endometrium from these donors^19^. Finally, the strong spatial and temporal dynamics of reproductive tissues mean that some rare or transient populations remain undersampled. This is particularly relevant for ovarian follicles that are highly temporarily and spatially localised, including advanced follicles such as ovulatory follicles, corpus luteum and corpus albicans, which are not covered in this atlas. Capturing these cell states will require targeted tissue screening with matched histological assessment to identify the relevant structures before profiling.

This atlas is part of the broader HCA initiative to build integrated reference atlases across the human body through a coordinated cross-community, cross-tissue effort. The hierarchical annotation framework of the HCA Reproductive Atlas, which resolves cell identity progressively from pan-organ lineages to tissue-specific states, is designed to be compatible with other organ atlases and to support future organism-level integration across lifespan. By integrating five organs within a unified analytical framework, the pipeline to build this atlas also provides a template for constructing pan-tissue atlases from single-organ integrated resources being generated across the HCA. Future versions should expand not only in cohort size, but also in their ability to capture rare, transient and spatially restricted cell states, which are a defining feature of the highly dynamic female reproductive system and are easily missed by dissociation-based approaches. Spatial transcriptomics, prospective tissue screening and matched histology will be essential to reach these under-sampled compartments. Coupled with modalities such as genotyping and methylation-based lineage tracing, single cell atlases will provide the foundation for studying gynaecological conditions that remain poorly understood despite their major clinical burden.

## Methods

### Single-cell RNA-seq datasets collection and inclusion criteria

Single-cell RNA-sequencing datasets of the female reproductive system (ovary, fallopian tube, uterus, cervix and vagina) generated using 10x Genomics platforms were systematically identified from public repositories and HCA network studies. Datasets were included if raw sequencing data were available, at least one tissue of interest was represented, and donors lacked active disease affecting the profiled organ. Overtly pathological tissues were excluded; uterine samples from donors with endometriosis were retained as a covariate given their prevalence and previously demonstrated minimal impact on endometrial cell type composition^19^.

### Single-cell RNA-seq profiling of fallopian tubes

#### Fallopian tube sample collection

Adult fallopian tubes were obtained from elective salpingectomies at the Rosie Hospital (Cambridge University Hospitals NHS Foundation Trust) with written informed consent and approval from the Cambridge local Research Ethics Committee (REC Reference 19/YH/0441). Specimens were processed within 24 h of surgery. Anatomical regions (fimbriae, infundibulum, ampulla, or isthmus) were recorded, and neighbouring regions were divided for spatial embedding or cryopreserved for single-cell analysis by mincing in ice-cold CryoStor CS10 solution (Sigma, C2874).

#### Fallopian tube tissue dissociation

Cryopreserved tissue was thawed, washed with HBSS, and digested for 35 min at 37°C with rotation in stromal digestion buffer containing 1 mg/mL Collagenase D (Merck, 11088858001), 1.5 mg/mL Hyaluronidase (Merck, H3506), and 100 µg/mL DNase I (Merck, 11284932001). The suspension was filtered through a 200 µm strainer (pluriStrainer, 43-50200-03) and neutralised with RPMI supplemented with 10% FBS. Residual tissue fragments were further digested in 0.25% Trypsin-EDTA (Thermo Fisher, 25200056) containing 100 µg/mL DNase I for 15 min at 37°C with rotation, filtered through a 100 µm strainer (Merck, CLS352360), and neutralised with RPMI + 10% FBS. Both fractions were pooled, centrifuged at 400g for 5 min at 4°C, and treated with 1x RBC Lysis Buffer (eBioscience, 00-4333-57) according to the manufacturer’s instructions. Cells were stained with FITC anti-human CD45 antibody (BioLegend, 304005) and 7-AAD viability dye (eBioscience, 00-6993-50). Cells were stained with FITC anti-human CD45 antibody (BioLegend, 304005; 1:100), 7-AAD viability dye (eBioscience, 00-6993-50), followed by cell hashing using TotalSeq anti-human antibodies (BioLegend, C0251–C0254) according to the manufacturer’s instructions for 10x Feature Barcoding technology. Cells were then FACS-sorted using a Sony MA900 sorter, and CD45^+^ and CD45^−^ populations were pooled at a 70:30 ratio before immediate processing for single-cell transcriptomics.

#### Libraries preparation and sequencing of 10x Genomics Chromium GEX

Single-cell libraries were prepared using the 10x Genomics Chromium Single Cell Next GEM 3’ v3.1, Next GEM 5’ v2 or GEM-X Single Cell 5’ v3 kits for transcriptomics according to the manufacturer’s instructions and with the aim to attain between 2,000 and 20,000 cells per reaction. For cell hashing workflow (CITE-seq), Chromium Next GEM Single Cell 5′ v2 with Feature Barcoding technology for Cell Surface Protein was used (5’ Feature Barcode kit) according to the manufacturer’s protocol (10x Genomics). All libraries were sequenced on an Illumina-HTP NovaSeq 6000 or Illumina NovaSeq X platforms with S2, S4 or SP flow cells, using paired-end sequencing, and to target a minimum of 50K reads per cell/nucleus (GEX libraries) or 10K reads per cell (CITE-seq, BCR and TCR). The sequencing format was Read 1: [28] cycles; i7 index: [10] cycles; i5 index: [10] cycles; Read 2: [90] cycles.

### scRNA-seq atlas construction

Raw sequencing files were aligned to GRCh38 using Cell Ranger v9.1 and denoised using CellBender v0.3.2^107^ to remove ambient RNA contamination. Cells were retained with >1,000 detected genes and <20% mitochondrial reads. Cell cycle phase was scored per cell^108^ and a transcriptional health score was computed from housekeeping gene expression as an orthogonal quality metric. Doublet scores were estimated per library using Scrublet^109^ and carried forward as metadata for cluster-level evaluation during annotation rather than applied as hard per-cell thresholds.

Integration was performed with scVI^110^ using the top 3,000 highly variable genes per population (Seurat_v3 method^108^), with dataset of origin as batch variable and donor identity and cell cycle phase as covariates (n_layers=1, n_latent=60, dispersion="gene-batch"). Leiden clustering (resolution=2; further readjusted manually) and UMAP visualisation were applied to the scVI latent representation using scvi-tools v1.4.2 and scanpy v1.12. Postnatal (paediatric, adult and postmenopausal) and prenatal cells were annotated independently. Integration proceeded hierarchically across three levels: broad lineage classification (epithelial, mesothelial, mesenchymal, endothelial, immune, granulosa, germ, and peripheral nervous system cells); per-lineage pan-organ sub-manifolds; and per-tissue sub-manifolds for epithelial, mesenchymal and gonadal supporting lineages. Fetal samples were integrated at the first two levels only. Germ cells were integrated across lifespan. Immune cells (pan-organ) were also integrated across lifespan (n_layers=2, dispersion="gene"), and macrophages, monocytes, and cDC2 were further integrated as a separate object to achieve fine grain annotation of cell states.

Cell type annotation was performed on all clusters passing quality control. Cluster-level artefacts – including doublets, low-quality cells, and donor-specific groupings – were identified and excluded based on QC metrics, Scrublet scores, co-expression of lineage-discordant markers, and absence of distinctive marker genes identified by TF-IDF (SoupX v1.5.0^111^). Cell type labels were assigned using published marker genes, de novo TF-IDF markers, donor metadata, and spatial localisation confirmed by Visium or Xenium spot deconvolution and smFISH for selected populations. We provide a detailed explanation in “Extended methods” (**Supplementary Information Note 1.1**).

### Spatial transcriptomics sample preparation

#### Tissue spatial embedding and sectioning

Fallopian tubes were embedded in cold OCT medium and flash-frozen in a dry ice-isopentane slurry. Tissues were sectioned at 10 µm thickness using a cryostat (Leica CM1950).

#### H&E staining and imaging

Fresh-frozen sections were air-dried before staining. Mayer’s haematoxylin solution was applied for 90 s, followed by 4-5 washes in deionised water, which also served to blue the haematoxylin. Sections were stained with 1% aqueous eosin applied manually by pipette and rinsed with deionised water after 1-3 s. Slides were dehydrated in absolute ethanol, cleared overnight in Neo-Clear (Sigma, 65351-M), coverslipped, air-dried, and imaged using a Hamamatsu NanoZoomer 2.0HT digital slide scanner.

### 10x Genomics Xenium Spatial Transcriptomics

Fallopian tube tissue cryosections (10 µM thickness) from the OCT blocks were processed using the 10X Genomics Xenium Spatial Transcriptomics platform, following the manufacturer’s protocols and using Xenium Prime 5K Human Pan Tissue & Pathways Panel + 100 add-on gene bespoke panel (design id: E2VR2Z). Briefly, samples were fixed with formaldehyde, permeabilised using SDS and methanol. Next, the tissue was incubated with target-specific probes for hybridisation, followed by ligation and rolling circle amplification for in situ signal enhancement. Prior to on-instrument analysis, autofluorescence quenching was performed. The slides were then stained with the Xenium Multi-Tissue Stain Mix as part of Multimodal Cell Segmentation workflow. The mix combines antibodies targeting plasma membrane (ATP1A1/E-cadherin), intracellular proteins (Vimentin) and a 18S ribosomal RNA probe to delineate cytoplasmic compartment, complementing DAPI nuclear staining. Finally, slides were then loaded on the Xenium Analyzer v2.0.1.0. Together with decoding reagents and consumables for iterative imaging, barcode decoding and multimodal cell segmentation. This multimodal staining strategy enhanced the identification of cell boundaries for segmentation.

### 10x Genomics Visium HD

Whole-transcriptome spatial gene expression profiling was performed on fresh frozen fallopian tube sections using the 10x Genomics Visium HD platform with CytAssist. Sections of 10 µm were cut with a cryostat, and tissue block preparation, sectioning, H&E staining, and imaging followed the Visium HD Fresh Frozen Tissue Preparation Handbook (CG000763). Probe hybridisation, probe ligation, slide preparation, probe release, extension, and library construction followed the Visium HD Spatial Gene Expression Reagent Kits User Guide (CG000685). Probe transfer from tissue slides to Visium HD capture slides was performed using the Visium CytAssist instrument (firmware v2.0.0 or higher), following the Tissue Slide Alignment Quick Reference Cards (CG000548). Libraries were sequenced on a NovaSeqX instrument, with a read configuration of 43 bp read 1, 50 bp read 2, and 10 bp dual indices. Spaceranger v4.1.0 was used to map reads to the reference, detect tissue, align data to the microscope and CytAssist images, and output feature-barcode matrices for further analysis.

### Spatial deconvolution of Visium HD data

Cell type deconvolution of Visium HD 8 µm bins for three reproductive tissues (uterus, fallopian tube and ovary) was performed using TACCO (v0.5.0)^112^. The HCA single-cell RNA-seq reference atlas was used as the annotation reference. For each tissue, only reference cells from the matching tissue were used to annotate the corresponding Visium HD sample, ensuring organ-specific deconvolution.

Raw count matrices were used as input for both the reference and query datasets. TACCO was run with default parameters (bisections=0), yielding a compositional score matrix of dimensions n bins × k cell types, where each row represents the fractional contribution of each cell type to a given spatial bin. To quantify spatial co-occurrence between cell type pairs, we computed soft co-occurrence matrices directly from the TACCO compositional scores using the TACCO co_occurrence_matrix function with max_distance=100. Unlike hard-label approaches, this method weights each spatial bin by its continuous cell type scores rather than assigning binary labels, thereby preserving compositional uncertainty and avoiding information loss from thresholding. The resulting co-occurrence score for each cell type pair reflects the observed frequency of spatial colocation relative to a random expectation, and was log_2_-transformed for visualisation and downstream analysis.

### Differential expression analysis between macrophage and ILC3 subsets

Pseudobulk differential expression analysis was performed using PyDESeq2^113,114^ (v0.5.4). Prior to each analysis, decoupler^115^ (v.2.1.6) was used to pseudobulk (sum) raw counts by donor and cell type and to filter low-quality pseudobulk samples and lowly expressed genes. Three comparisons were performed: 1) a one-versus-rest comparison across uftLAMs, LYVE1^hi^ macrophages, and oLAMs; 2) a pairwise comparison between uftLAMs and oLAMs; and 3) a pairwise comparison between adult NCR^hi^ ILC3 derived from the reproductive tract versus non-reproductive tract tissues (lung, oral cavity, colon). NCR^lo^ ILC3 were excluded from the last comparison due to their absence in the non-reproductive tract datasets. All comparisons used a Wald test with Benjamini-Hochberg FDR correction. Genes with an adjusted p-value < 0.05 and absolute log_2_ fold change > 1 were considered differentially expressed. The results for all comparisons are presented in **Supplementary Table 3.**

### Cell type enrichment for reproductive traits in the Human Female Reproductive System Cell Atlas v1

#### GWAS selection

To identify cell types enriched for gynaecological trait heritability, we first selected 7 GWAS focusing on the most recent, well-powered studies of common disorders, covering endometrial and myometrial disorders, heavy menstrual bleeding, polyendocrine metabolic ovarian syndrome (PMOS, formerly PCOS) and two high-powered reproductive timing traits, namely age at menarche and menopause **(Supplementary Table 4)**. Given the importance of linkage disequilibrium in downstream analyses, we prioritised European cohorts. Summary statistics for these GWAS were downloaded and lifted to GRCh38/hg38.

#### LDSC cell type enrichment

To identify cell types relevant for common gynaecological disorders using scRNA-seq, we used CELLECT-LDSC^116^, which integrates scRNA-seq with GWAS summary statistics. Briefly, CELLECT-LDSC is a pipeline which draws 100kb windows on either side of specifically expressed genes per cell type to perform stratified LD score regression (S-LDSC) partitioning and identify cell types enriched for trait heritability^117^.

To run LDSC on European (EUR) reference genome build GRCh38/hg38, we downloaded reference 1000 genomes data and baseline LD scores from https://zenodo.org/records/10515792. We computed unpartitioned LD scores from the 1000 genomes data to calculate observed scale heritability per GWAS. For S-LDSC, we adapted CELLECT-LDSC to use the GRCh38/hg38 inputs and used CELLEX scores as input. CELLEX scores^116^ are gene expression specificity scores which averages out different cell type expression specificity statistics to compute a consensus specificity score per gene per cell type. Fine-grain cell types were grouped into coarse categories and CELLEX was run on the whole 10x 5’/3’ scRNA-seq atlas in order to maximise cell type specificity across tissues^118^. Given that we were interested in ovarian and uterine traits, we subset CELLEX outputs to (a) common cell types in the uterus and ovaries, (b) cell types only in the ovaries and (c) cell types only in the uterus. CELLECT-LDSC was run per ovarian GWAS on CELLEX scores for (a)+(b) cell types and per uterine GWAS on CELLEX scores for (a)+(c) cell types. We included ADHD^119^ as a control phenotype in both analyses. We corrected p-values for multiple testing across all cell types within GWAS using the *q-value* package (v2.34.0).

To compare effect size (τ) between annotations, we calculated the standardised τ*, which is the per-SNP heritability change per standard deviation of the annotation^117^ as:

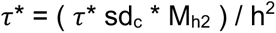

Where τ is the reported LDSC coefficient, sd_c_ is the standard deviation of annotation c (i.e. the per-SNP annotation output by CELLECT per cell type on which LDSC is run), h^2^ is the observed SNP heritability for the given GWAS and M_h2_ is the number of SNPs on which h^2^ was computed. We computed h^2^ per GWAS using the same hg38 LDSC baseline and weights files as were used to run CELLECT-LDSC.

To follow up on stromal cell enrichments in uterine traits, we subset to stromal fibroblasts only and reran S-LDSC to resolve fine-grain enrichment. To compute the linear association between endometriosis risk and LDSC, we used a weighted least squares regression on τ* values by cell state ranking across the menstrual cycle, weighted by the inverse of the squared τ* standard error. For these analyses, stromal fibroblasts were ranked by their enrichment in donors across the menstrual cycle, reflected in their HCA annotation.

To identify disease pathways driving cell type enrichment for a given cell-trait pairs, we selected all genes with nonzero CELLEX scores for that cell type and iteratively reran S-LDSC in a leave-one-out (LOO) approach, setting each gene’s CELLEX score to 0. We computed ΔZ_j_ for each “deleted” gene as:

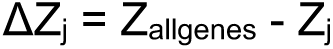

Where Z_allgenes_ is the Z-score of association between the trait and cell type including all genes and Z_j_ is the Z-score of association between the trait and cell type after setting the CELLEX score of gene j to 0.

In order to extract biological information from the LOO analysis without relying on individual genes whose association could be driven by non-coding SNPs which are difficult to assign to a gene, we performed gene-set enrichment analysis (fGSEA) within “deleted” genes per trait–cell type association using ΔZ_j_ as input to rank and score genes. Leading edge genes from nominally significant fGSEA gene sets with ΔZ_j_ > 0 were prioritised for biological interpretation. To add a layer of cell type specificity when interpreting our results within stromal fibroblasts, we subset the fGSEA prioritised genes to the top 100 with the highest ΔZ_j_ and identified those which were most specific to the enriched cell type, ranking them by CELLEX scores from our within-fibroblast analysis.

### Single-cell ATAC-seq atlas construction

scATAC-set datasets across the female reproductive system^15,120^ were collected and additional endometrial samples generated (detailed below) and uniformly reprocessed using cellranger-ATAC (v2.0.0). Similarly to the transcriptomic atlas, we harmonised metadata and processed all datasets in the same way to ensure consistency across the atlas.

#### Endometrial biopsies and single-nuclei extraction

Superficial endometrial biopsies have been collected through the “Sanger Human Cell Atlasing Project” study (Yorkshire & The Humber–Leeds East Research Ethics Committee (REC): 19/YH/0441). Written informed consent was obtained from study participants before tissue samples and phenotypic data were collected. Endometrial biopsy samples cryopreserved in CryoStor CS10 (STEMCELL Technologies; cat. no. 210502) were thawed at 37°C and immediately transferred to ice-cold RPMI 1640 medium (Gibco, cat. no. 21875-034). Tissue was centrifuged (500 × g, 5 min, 4°C) and resuspended in RPMI 1640 containing collagenase V (1 mg/mL; Sigma, cat. no. C9263), fetal bovine serum (FBS, 10% (v/v); Gibco, cat. no. A52567-01), and DNase I (0.1 mg/mL; Sigma, cat. no. 11284932001) for 15-60 min at 37°C with rotation. Following centrifugation (800 × g for 2 min), samples were washed in Dulbecco’s phosphate-buffered saline (PBS; Gibco, cat. no. 14190144) and incubated in red blood cell lysis buffer (Invitrogen, cat. no. 00-4333-57) for 5 min. Cells were washed in DPBS and resuspended in stromal fibroblast medium consisting of DMEM, high glucose (Gibco, cat. no. 41965-039) supplemented with FBS (10% (v/v)) and Primocin (0.1 mg/mL; InvivoGen, cat. no. ant-pm-1). Cell suspensions were passed through a 40 µm reversible strainer, and the flow-through was maintained on ice. Retained tissue fragments were further digested with TrypLE Select (Gibco, cat. no. 125630) for 15 min, followed by quenching with DMEM (high glucose) supplemented with FBS (10% (v/v)). Both fractions were centrifuged (800g, 2 min) and recombined in red blood cell lysis buffer for 5 min. Cells were washed in DPBS, resuspended in stromal fibroblast medium, and maintained on ice for downstream scATAC-seq. Nuclei were subsequently isolated using the Nuclei Isolation protocol (10x Genomics, CG000169) according to manufacturer instructions.

#### Library preparation and sequencing

Single-cell/nucleus ATAC libraries were prepared using the 10x Genomics Chromium Single Cell Next GEM ATAC v2, both according to the manufacturer’s instructions and with the aim to attain between 2,000 and 10,000 cells/nuclei per reaction. All libraries were sequenced on an Illumina-HTP NovaSeq 6000 or Illumina NovaSeq X platforms with S2, S4 or SP flow cells, using paired-end sequencing, and to target a minimum of 50K reads per cell/nucleus. The sequencing format was: Read 1: 50 cycles; i7 index: 8 cycles; i5 index: 16 cycles; Read 2: 50 cycles.

#### Read alignment, quantification and quality control

Raw snATAC-seq fragments were aligned by cellranger-ATAC (v2.0.0) to genome build GRCh38/hg38 and cells were filtered using EmptyDropsMultiome^121^ using the unfiltered peak matrices from cellranger-ATAC to retain real cells and maximise recovery of ovarian germ cells, which are otherwise difficult to recover. Cells with <10,000 fragments were removed to keep high-quality cells.

#### Integration and clustering

The following steps were performed by tissue and developmental stage (fetal or nonfetal). SnapATAC2 (v2.9.0)^122^ was used for dimensionality reduction using a Spectral embedding. Per-sample and per-dataset integration was performed using SnapATAC2’s built-in Harmony implementation. Scanpy nearest neighbours, UMAP generation and leiden clustering (at resolutions of 1 and 5) were performed on the Harmony latent space.

#### Cell type annotation, filtering and peak calling

We used ArchR (v1.0.3)^123^ in R (v4.3.1) for cell type annotation and peak calling. The snapATAC2 Harmony embedding was imported into R and ArchR gene scores were computed using a custom gene annotation from the 2020A 10x hg38 reference gene annotation, subset to 35,158 genes found in the scRNA-seq data. To set a reference for scATAC-seq datasets, the scRNA-seq atlas was downsampled to 10,000 cells per fine-grain cell type while maintaining sample proportions, and subset to each relevant tissue and life stage. We implemented ArchR’s built-in Seurat integration (addGeneIntegrationMatrix()) using the snapATAC2 Harmony embedding and restricting features to ArchR gene activity scores for the union of the top 100 marker genes per fine-grain cell type computed using a TF-IDF approach.

Resulting fine-grain annotations were grouped into higher confidence coarse annotations. Different cluster resolutions were used for different filtering steps to prioritise the cleanest final set of cells. Doublets were called where SnapATAC’s doublet probability > 0.75, and clusters (at resolution = 5) with over 10-20% of doublets were removed depending on the dataset, along with any remaining doublets. Clusters dominated by a single donor or with low transcription start site (TSS) enrichment without biological explanation were removed. Cells in the lowest 20% of annotation prediction scores per cell type were removed, except rare cell types like germ cells and ovarian surface epithelium. Clusters were assigned to a lineage by majority voting and cells whose lineage did not match their cluster were removed, whilst ensuring that clusters of rare cell types were not lost. Finally, we removed cell types with <10 cells if they did not form an independent cluster.

The annotated, high quality cells per tissue were concatenated in SnapATAC2 and integrated by donor and dataset to refine our final cell type annotation. The final snATAC-seq annotation was used to call reproducible peaks per cell type using ArchR’s MACS2 implementation.

To verify the predicted annotation, we used marker genes identified from the scRNA-seq atlas and the top genes per cell type. Using the new scATAC-seq coarse annotation groupings, the corresponding scRNA-seq cells (by tissue and developmental stage) were re-grouped to match the annotation and the top 20 marker genes per coarse annotation were identified using the TF-IDF approach. Mean gene activity scores per coarse annotation per gene were computed from the scATAC-seq and cross-cell type Z-scores were computed per gene to check the quality of our annotation, using both top 20 markers and canonical markers.

To confirm the functional relevance of our peaks, we used regioneR (v1.34.0) to randomly permute our peaks 1000 times across the genome and empirically derived enrichment for different ENCODE annotations in our peaks. This was performed for intergenic and all peaks to assess the P-values were corrected across tests using the qvalue package (v2.34.0).

#### Peak to gene links

We used ArchR to compute peak to gene associations. To avoid abundant cell types overpowering associations, we downsampled each to a maximum of 1000 cells. To account for data sparsity and imperfect snATAC-seq–scRNA-seq integration, ArchR groups cells into “metacells” of 200 nearest neighbours, which was more conservative than the ArchR defaults. Peak to gene links were then computed using metacells by correlation of peak accessibility and integrated scRNA-seq gene expression within 500kb of each gene. For plotting and prioritisation of top cell types, correlation was recomputed by pseudobulking by cell type.

To maximise confidence in our peak to gene links, we only retained those with FDR<1e-5 and asked whether peaks that were linked to genes were more likely to be enriched in functional ENCODE elements than peaks not linked to genes. We computed the enrichment of linked versus unlinked peaks for ENCODE annotations using Fisher’s exact test, both using all peaks and using only intergenic peaks (removing those overlapping with the gene body). To further check that our peak to gene links are robust, we downloaded experimentally tested peak to gene links across multiple cell lines (https://github.com/EngreitzLab/CRISPR_comparison/tree/main/resources/crispr_data)^124^ and subset both our peaks and the tested enhancers to the overlapping genomic ranges. We then tested for enrichment of validated enhancers to genes links in our peak–gene links using Fisher’s exact test.

#### GWAS locus overlap with peaks

To identify overlap between GWAS loci and open chromatin peaks and identify putative effector genes, we selected credible sets from Open Targets using the EFO ontology: all traits under HP_0000008 (Abnormal morphology of female internal genitalia) and EFO_0009549 (female reproductive system disease) terms were selected, and additional traits were selected from the female-specific non-developmental traits under HP_0000078 (Abnormality of the genital system), namely menstrual abnormalities (HP_0400007, HP_0100608, HP_0100607, HP_0000876, HP_0000858, HP_0000132) and premature ovarian insufficiency (HP_0008209). We also used index loci from GWAS which we used for cell type enrichment (see below) which were not in Open Targets, lifted to GRCh38/hg38 where necessary. We then intersected these loci with our peaks linked to genes with FDR<1e-5, excluding those located in the gene body of the gene they were linked to.

## Data and code availability

We provide user-friendly access to our annotated scRNA-seq resource and our and to our Spatial Transcriptomics samples via cellxgene at our https://www.reproductivecellatlas.org/HCAreproductive/v1/. All the raw and processed sequencing data generated in this study are currently being deposited to EGA and BioImageArchive (10x Xenium). The code used to perform the analyses presented in the manuscript can be found at https://github.com/ventolab/HCA_female_reproductive_system. Cell type annotations for scRNA-seq and scATAC-seq, CELLEX inputs for cell type enrichment, leave-one-out ΔZ scores per gene, all peaks and peak to gene links can also be found at https://github.com/ventolab/HCA_female_reproductive_system.

## Acknowledgments

We wish to acknowledge the donors, patients and relatives who provided tissues for this research. We also acknowledge Rosie Research midwives and gynaecology team at Cambridge University Hospitals NHS Foundation Trust for their support in donor recruitment and informed consent procedures; the transplant organ donors and their families for their generous tissue donations through the Cambridge Biorepository for Translational Medicine; Prof. John R.B. Perry and Dr. Felix Day for providing access to PCOS / PMOS summary statistics from their preprint (Moolhuijsen et al. 2026, now published); Prof. Triin Laisk for sharing the FGT polyps summary statistics whilst not yet available on GWAS Catalog; Dr. Tobi Alegbe for his advice on cell type enrichment analyses; Batu Cakir and Martin Prete for their help on the reproductivecellatlas.org web portal; Tarryn Porter, Heather Stanley, the Spatial Genomics Platform (SGP) and Sanger Core Sequencing pipeline for support with sample processing and sequencing library preparation; A. García from Bio-Graphics for scientific illustrations; and A. Maartens for his help on manuscript writing. This publication is part of the Human Cell Atlas – www.humancellatlas.org/publications/.

## Author contributions

C.E.C., L.G-A. and R.V-T. conceived and designed the study. A.P., I.K., J.M.A., J.G-E., and M.B. contributed to fallopian tube and uterine sample acquisition. A.P. and C.S-S. performed sample dissections and sample processing for single-cell and single-nuclei profiling. C.I.M. performed sample processing for the imaging experiments. L.G-A. coordinated data analysis, which was performed by C.E.C., A.P-L., L.R-M., C.E.K., M.M., A.Pr. and L.G-A. with contributions from R.V-T., A.P., V.L., R.V-B., E.A. and K.P. Cell type annotation was defined by L.G-A., C.E.K., V.L. and R.V-T. with oversight from The Human Cell Atlas Reproductive network coordinators A.S., S.S.H., C.L., and R.V-T. and reviewed through the HCA Reproduction Network workshops by M.Ma., L.M., N.D.U., D.N.A., L.F., T.S., F.W., G.E., T.T., J.M.A., M-T.B., N.K., V.P-G., F.C., C.B., K.T.Z., A.C., S.W., T.C-D., N.G-L., S.B., A.D.R., A.Ba., A.M., F.V., E.S-V., P.D., C.S., S.M.C.d.S.L. C.E.C. performed the genetic integration analyses, supervised by L.Fa., B.H and C.A.A.. C.E.C., A.P-L., L.R-M., C.E.K., L.G-A. and R.V-T. interpreted the data, with contributions from all remaining authors. C.E.C., L.R-M., C.E.K., L.G-A. and R.V-T. wrote the manuscript, with feedback from M.Ma., T.C-D., K.T.Z., N.G-L., F.V., E.S-V., P.D., C.S., S.M.C.d.S.L., C.L., A.S., S.S.H., L.Fa. and C.A.A. L.G-A. and R.V-T. supervised the work. All authors read and approved the manuscript.

## Table Legends

**Supplementary Table 1. Datasets and donors metadata in the Human Female Reproductive System Cell Atlas v1. A,** Per sample metadata. Table showing harmonised metadata for all samples and libraries included in the Human Female Reproductive System Cell Atlas v1, following Human Cell Atlas metadata conventions. Fields include organ and anatomical sampling site, dataset and library identifiers, donor-level demographic and developmental information, reproductive and menstrual status, disease or pathology annotations, tissue and specimen processing details, enrichment and sorting strategies, assay and sequencing metadata, and dataset-level batch and collection site information. Each row corresponds to a profiled library or sample (for multiplexed libraries). **B,** Novel samples profiled for scRNAseq. Table showing relevant metadata for novel fallopian tube samples profiled for 10x scRNAseq transcriptomics. **C,** Novel samples profiled for spatial transcriptomics Xenium and Visium HD. Table summarising the spatial transcriptomics samples from adult donors profiled using either Visium HD or Xenium with a bespoke 480-gene panel. These samples were used for spatial validation and anatomical mapping of atlas-defined cell populations across reproductive tissues. **D,** Per library QC as reported in Cell Ranger v9 web summaries

**Supplementary Table 2. Cell labels, taxonomic classification and markers defining the Human Female Reproductive System Cell Atlas v1. A,** HCA cell type classification system used in the atlas, in which each HCA cell type label is constructed hierarchically as: lineage + broad cell type + fine cell type. **B,** Structured cell classification developed within the HCA Reproduction Network for the Human Female Reproductive System Cell Atlas v1. For each atlas HCA cell type label defined in A, the table provides the corresponding classification in a four-level hierarchy, from lineage-level categories (L1) to increasingly resolved cell type and cell state annotations (L2–L4). The table also includes positive and, where applicable, negative marker genes supporting each annotation, a brief cell type description, and alternative cell type labels used in the literature or source datasets.

**Supplementary Table 3. Differentially expressed genes in immune cells. A,** Differentially expressed genes between uftLAM, Mac_LYVE1hi, and oLAM, identified via one-versus-rest method (see Methods). **B,** Differentially expressed genes between uftLAM and oLAM, identified via pairwise DEG analysis. **C,** Differentially expressed genes reproductive tract- and non-reproductive tract- (lung, oral cavity, colon) derived ILC3_NCRhi, identified via pairwise DEG analysis.

**Supplementary Table 4. Data and results for cell type enrichment analyses. A,** Gynaecological condition GWASs selected for cell type enrichment. **B,** Cross-atlas CELLECT-LDSC results. Columns include the output from CELLECT-LDSC (beta–pvalue), observed heritability per GWAS and associated SE (h2_obs, h2_se), number of SNPs on which heritability was computed (n_snps), standard deviation of the per-SNP annotation input for S-LDSC analysis, namely CELLEX scores, standardised effect and standard error (tau_star and tau_star_se – tau_star is the change in per-SNP heritability per standard deviation of the annotation), and cell lineage used for plotting. **C,** CELLECT-LDSC results within endometrial fibroblasts. Refer to Supp. Table 4.B. legend for column details. **D,** LOO-LDSC fGSEA results for top trait–cell type associations. Listed are the nominally significant fGSEA results with ES>0.

**Supplementary Table 5. Chromatin accessibility atlas metadata and peak–gene links. A,** Single-cell ATAC-seq sample metadata. **B,** Disease-overlapping peak–gene links where the top gene has FDR<1e-5. L2g information comes from Open Targets L2G effector gene prediction, which favours nearby genes, and Correlation, FDR and geneName are all the top associated gene at that peak. Genes may include multiple other genes with FDR-significant peak correlation and peak_info discriminates peaks linked with a single versus multiple genes. Loci from Open Targets are from credible sets and include l2g columns where non-Open Targets loci have NAs. Non-Open Targets loci are the top index SNP per locus as reported by each respective GWAS.

