## Supplementary Information for "An integrated multimodal pan-organ atlas of the female reproductive system across the lifespan contextualises gynaecological pathologies"

Supplementary notes

**Supplementary Note 1. Extended Methods.	2**

Note 1.1. Building the Human Female Reproductive System Cell Atlas v1	2

Data collection and inclusion criteria	2

Read alignment, quantification and ambient RNA removal	4

Per-cell quality control	4

Per-cell phenotypic scores	4

Data integration and clustering	5

Integration hierarchy	5

Cell type annotation	6

Nomenclature and Cell Classification mapping	7

Computationally friendly labels	8

Note 1.2. Integration with other organ datasets	9

Integration of reproductive tract-derived NCRhi ILC3s with those from other mucosal tissues.	9

**Supplementary Note 2. Extended results.	9**

Note 2.1 Novel insights from postnatal Human Female Reproductive System Cell Atlas v1	9

Organisation of the epithelium and mesothelium	10

Organisation of the mesenchyme	11

Organisation of the vascular endothelial cells	15

Organisation of the immune cells	15

Organisation of the peripheral nervous system cells	17

Organisation of ovarian follicular cells	17

Note 2.2. Expression of steroid receptors along all the cell types of the female reproductive HCA	19

Note 2.3. GWAS integration with the Human Female Reproductive System Cell Atlas v1	20

Note 2.4. snATAC-seq Human Female Reproductive System Cell Atlas	21

**Supplementary Notes References	21**

#### Supplementary Note 1. Extended Methods.

##### Note 1.1. Building the Human Female Reproductive System Cell Atlas v1

Here we describe the construction of the first version of the *Human Female Reproductive System Cell Atlas v1*.

Briefly, we systematically collected, reprocessed, integrated and annotated the available single-cell transcriptomic datasets from human ovary, fallopian tube, uterus, cervix and vagina profiling controls for which sequencing data was available. These datasets span prenatal development through postmenopause, including first- and second-trimester fetal samples, paediatric samples (ovary only), reproductive-age adults and postmenopausal donors, and include newly generated data from HCA network (fallopian tubes) and two manuscripts currently under revision (Lorenzi et al. 2026; Garcia-Alonso et al. 2026; under review, preprints in bioRxiv). All data were re-processed using a unified preprocessing, integration and re-annotation framework to generate a consensus cellular reference of the female reproductive system across the lifespan. This harmonisation is essential, particularly in reproductive tissues, where hormone-responsive states introduce substantial additional biological variation between donors.

Given the substantial transcriptional differences between fetal and postnatal biology, we annotated fetal (comprising first and second trimester) and postnatal (comprising pediatric and adult stages, including postmenopause) data individually and later combined both developmental stages into an across lifespan unified atlas.

###### Data collection and inclusion criteria

Single-cell RNA-sequencing datasets of the female reproductive system (ovary, fallopian tube, uterus, cervix and vagina), generated using 10x Genomics-based platforms, were systematically identified from public repositories and from studies under review within the HCA network (Lorenzi et al. 2026; Garcia-Alonso et al. 2026; under review, preprints in bioRxiv). Datasets profiling dissociated single cells were included if: (i) raw sequencing data (.fastq or .bam) were available, (ii) the tissue of interest was profiled, and (iii) samples were derived from donors without active disease affecting the profiled organ, except for the few cases noted below.

Datasets profiling overtly pathological tissue were excluded, including fallopian tube samples with confirmed hydrosalpinx, cervical cancer samples, or vaginal samples from donors with pelvic organ prolapse. Tissues were accepted if obtained from deceased organ donor programmes, from hysterectomy specimens, from patients undergoing biopsy procedures for conditions not directly affecting the sampled organ (for example, ovarian tissue from donors with non-gynaecological cancers), and menstrual fluid collection. Uterine samples from donors with endometriosis were included as an exception, given that endometriosis represents one of the most prevalent gynaecological conditions and its inclusion was necessary to achieve adequate representation of the endometrium across the menstrual cycle. We previously showed that endometriosis does not cause the emergence of novel endometrial cell types, and that gene-expression changes across endometrial cells in the presence of endometriosis are mild^1^. Endometriosis status was retained as a metadata covariate. Metadata including donor age, menopausal status, menstrual cycle phase where available, tissue type and sample provenance were harmonised across studies prior to integration.

While this *Human Female Reproductive System Cell Atlas v1* effort aimed to include both paediatric and adult tissues, paediatric sampling was limited to ovarian cortical tissue obtained through fertility preservation programmes. For completeness and to provide developmental context, the atlas also incorporated prenatal (1st and 2nd trimester) counterparts of these tissues, as well as external genitalia datasets from Lorenzi et al. (2026). A full list of datasets, accession numbers and donor metadata is provided in **Supplementary Table 1**. Table below provides an overall summary (**Supplementary Note 1.1. Table 1**).

| **Organ** | **Developmental stage** | **Dataset** | **Samples** |
| --- | --- | --- | --- |
| Ovary | Fetal | Lardenois 2026 | 16 |
| Ovary | Fetal | Taelman 2024 | 4 |
| Ovary | Fetal | Wamaitha 2023 | 5 |
| Ovary | Fetal | Garcia-Alonso 2022 | 55 |
| Ovary | Pediatric (postnatal) | Garcia-Alonso (under revision) | 38 |
| Ovary | Adult (postnatal) | Garcia-Alonso (under revision) | 18 |
| Ovary | Adult (postnatal) | Gaylord 2025 | 7 |
| Ovary | Adult (postnatal) | Jones 2024 | 3 |
| Ovary | Adult (postnatal) | Guahmich 2023 | 8 |
| Ovary | Adult (postnatal) | Wagner 2020 | 4 |
| Reproductive tract | Fetal | Lorenzi 2026 | 50 |
| Fallopian Tube | Adult (postnatal) | Unpublished (new) FallopianTube | 7 |
| Fallopian Tube | Adult (postnatal) | Weigert 2025 FallopianTube | 39 |
| Fallopian Tube | Adult (postnatal) | Ulrich 2022 FallopianTube | 10 |
| Uterus | Adult (postnatal) | Lorenzi 2026 (submitted) Endometrium | 38 |
| Uterus | Adult (postnatal) | Lorenzi 2026 (submitted) MenstrualFluid | 11 |
| Uterus | Adult Menopause (postnatal) | Lorenzi 2025 (submitted) Uterus | 11 |
| Uterus | Adult (postnatal) | Burns 2026 | 11 |
| Uterus | Adult (postnatal) | Liu 2025 | 10 |
| Uterus | Adult (postnatal) | Ulrich 2024 | 10 |
| Uterus | Adult (postnatal) | Mareckova 2024 | 33 |
| Uterus | Adult (postnatal) | Huang 2023 | 12 |
| Uterus | Adult (postnatal) | Tan 2022 | 12 |
| Uterus | Adult (postnatal) | Garcia-Alonso 2021 | 18 |
| Uterus | Adult (postnatal) | Wang 2020 | 10 |
| Cervix | Adult (postnatal) | Guo 2023 | 2 |
| Vagina | Adult Menopause (postnatal) | Li 2021 | 5 |

***Supplementary Note 1.1. Table 1. Summary of datasets included in the Human Female Reproductive System Cell Atlas v1*** *Each row indicates one dataset, with the organ profiled, the developmental stage, the reference manuscript and the number of samples.*

###### Read alignment, quantification and ambient RNA removal

Raw sequencing files were downloaded from the corresponding public repositories or source, and processed uniformly using a standardised pipeline. Reads were aligned to the GRCh38 human reference genome and transcripts quantified using Cell Ranger v9.1 (10x Genomics), applying default parameters. To account for ambient mRNA contamination (a systematic source of noise in droplet-based single-cell experiments), raw count matrices were denoised on a per-library basis using CellBender v0.3.2^2^ (remove-background module), which employs a deep generative model to distinguish true cell-associated counts from background ambient RNA. The CellBender-denoised count matrices were used for all downstream quality control and integration steps.

###### Per-cell quality control

Per-library quality control was applied to the CellBender-denoised count matrices to discard low quality cells prior integration. Cells were retained on the basis of a minimum number of detected genes (>1,000 genes per cell) and a maximum mitochondrial read fraction (<20%), with thresholds applied uniformly across libraries. Per-library cell counts and QC metrics are reported in **Supplementary Table 1d**.

###### Per-cell phenotypic scores

Beyond basic count-based filtering, we computed a set of per-cell phenotypic metrics to characterise each cell's technical state prior to integration. *Cell cycle phase* was assigned to each cell using the scoring approach proposed by Stuart et al., 2019^3^, based on the expression of established G1/S and G2/M phase marker gene sets, allowing cell cycle effects to be monitored during downstream analysis. Using a similar approach, we computed a per-cell *transcriptional health score* based on the aggregate expression of a curated panel of well-established housekeeping genes, providing an orthogonal measure of overall cell quality independent of mitochondrial fraction, particularly useful for identifying cells activating cell death programmes independent of cell stress, where mitochondrial read fractions may not reliably reflect cell integrity.

*Doublet* detection was performed on a per-library basis using Scrublet^4^. Scrublet simulates artificial doublets from the observed data to estimate a per-cell doublet probability score. While Scrublet provides valuable information, its accuracy is inherently limited by library-specific cell type composition and the clustering resolution used for score calibration, factors that are difficult to optimise uniformly across the large number of libraries integrated here. In particular, rare cell types present at low frequency within a given library may be poorly represented in the simulated doublet population, leading to underestimation of doublet scores for cells from those populations. For these reasons, doublet scores were not used as a hard filtering threshold at the per-library stage. Instead, doublet scores were retained as per-cell metadata and carried forward into the integrated analysis, where they were evaluated at the cluster level alongside marker gene specificity during the annotation process. This approach affords greater power to distinguish genuine rare cell types from artifactual doublets, and is described in detail in the Cell type annotation section below.

###### Data integration and clustering

Cells passing quality control were integrated using a standard analytical pipeline. Integration was performed with scVI^5^, a deep generative model that learns a shared low-dimensional representation of gene expression while accounting for technical variation across datasets.

The model was trained on the top 3,000 highly variable genes identified within each cell population being integrated, using the Seurat_v3 method^3^ on raw count data. To maximise biological resolution at each level of the integration hierarchy, highly variable genes were recalculated within the specific cell population being integrated; for example, immune cell integrations used highly variable genes identified exclusively within immune cells. Dataset of origin was specified as the batch variable, while donor identity and cell cycle phase were included as covariates to account for dataset-specific, donor-specific and cell cycle–associated variation. Model parameters (n_layers = 1, n_latent = 60, dispersion = “gene-batch”) were selected to minimise technical variation while preserving biological signal and biologically meaningful dataset structure.

Nearest-neighbour graphs were constructed from the resulting scVI latent representation. Uniform Manifold Approximation and Projection (UMAP) was used for visualisation, and Leiden clustering was applied to the neighbour graph (maximum resolution = 2). All analyses were implemented using scvi-tools^6^ (v1.4.2) and scanpy^7^ (v1.12).

To assess integration quality, we examined whether biologically coherent cell populations aligned across tissues and datasets while preserving expected biological differences. Cell types known to be transcriptionally conserved across tissues (for example, T cells and peripheral nervous system cells) clustered together irrespective of dataset, indicating effective correction of technical variation. In contrast, transcriptionally related but functionally distinct populations (for example, secretory epithelial cells from fallopian tube and endometrium) remained separated in the integrated space, confirming that biologically meaningful differences were preserved.

###### Integration hierarchy

Integration, clustering and annotation of the postnatal datasets were performed in a hierarchical manner across three levels of resolution:

- At the first level, cells in the main integrated manifold were classified into broad lineages (epithelial, mesothelial, mesenchymal, endothelial, immune, granulosa, germ cells and peripheral nervous system cells) using canonical lineage markers.
- At the second level, each lineage was extracted and re-analysed independently, generating per-lineage pan-organ sub-manifolds that offered greater resolution of closely related cell states than the main manifold alone.
- For the epithelial and mesenchymal lineages, which showed the greatest within-lineage diversity across the five tissues, a third level of analysis was performed in per-tissue sub-manifolds, capturing fine-grained, organ-specific cell states that would otherwise be obscured by the numerical dominance of more abundant populations.

Fetal integration followed the same two first levels of this hierarchy. A third level of per-tissue analysis was not performed because, unlike postnatal samples, fetal samples frequently contain cells from multiple organ primordia that cannot be reliably separated due to the difficulty of achieving tissue-specific dissections at early developmental stages, where the boundaries of the tissues are less well defined.

Cell type labels assigned at each level were subsequently projected back onto the main integrated manifold for visualisation and cross-tissue comparison. The annotation strategy applied at each level is described in detail in the Cell type annotation section below.

For clarity, we distinguish throughout this manuscript between two types of UMAP representations: *analytical manifolds*, on which clustering and cell type annotation were performed, and *visualisation manifolds*, which display curated subsets of cells to highlight specific populations or comparisons. Visualisation manifolds were generated for illustrative purposes only and were not used for annotation decisions; this distinction is indicated in the relevant figure legends.

###### Cell type annotation

We performed a comprehensive re-annotation of all clusters identified in the integrated *analytical manifolds*. This proceeded in two stages: first, removal of clusters likely to represent technical artefacts; and second, assignment of cell type labels to the remaining high-quality clusters.

Cluster-level quality control and artefact removal

Prior to annotation, we identified and excluded clusters representing unreliable cell identities, including doublets, low-quality cells (for example arising from cell damage, stress or apoptosis), and technically driven groupings lacking biological support. Distinctive marker genes for each cluster were identified using the TF-IDF approach implemented in SoupX^8^ (v1.5.0), and were central to all decisions.

- Clusters were flagged as **low quality** if they met all of the following criteria: (1a) lower-than-average total gene counts per cell; or (1b) lower-than-average UMI counts per cell; or (1c) higher-than-average expression of mitochondrial or stress-response genes (JUN family, FOS family, NR4A family); or (1d) higher-than-average expression of nuclear RNA suggestive of cell membrane rupture; or (1e) downregulation of housekeeping genes; and (2) absence of distinctive marker genes by TF-IDF.
- Clusters were flagged as **doublets** if they met the following criteria: (1) elevated mean Scrublet doublet score relative to the dataset distribution; (2) co-expression of marker genes from multiple distinct lineages or distal broad cell types, like simultaneous expression of epithelial and immune markers; and (3) absence of any distinctive marker gene by TF-IDF. This last criterion was weighted particularly strongly, as clusters lacking transcriptional features beyond the combined signatures of their putative singlets are difficult to distinguish from technical doublets using scRNA-seq data alone. Conversely, clusters with elevated doublet scores but coherent and distinctive marker programmes (for example, cells undergoing cell cycle) were retained, as these features support the presence of a genuine biological cell state rather than an artifactual mixture.
- Clusters dominated by cells from a **single dataset or donor,** without biological metadata supporting their separation, were additionally flagged for exclusion. For example, a cluster arising from a single endometrial donor in the proliferative phase was excluded on the grounds that it lacked distinctive marker genes despite the availability of matched biological replicates from the same menstrual stage. In contrast, clusters enriched in a specific dataset for biologically interpretable reasons were retained, like theca cells enriched in the Guahmich et al. ovarian dataset^9^, where tissue sampling and dissociation conditions favour recovery of this specialised population, and whose identity was supported by canonical marker expression.

Cell type label assignment

Cell type labels were assigned to all clusters passing quality control following the hierarchical clustering strategy described above (*“Integration hierarchy”* section). At each level of this hierarchy, annotation decisions drew on four sources of evidence applied in combination.

The primary evidence was the expression of marker genes previously described for cell types of the ovary, fallopian tube, uterus, cervix and vagina in the published literature. This was complemented by cluster-specific marker genes identified de novo by TF-IDF, which highlighted genes uniquely enriched in each cluster regardless of prior knowledge. A third source of evidence was biological and clinical metadata, including tissue sampling strategy (for example, whole ovary versus isolated cortex, or whole uterus versus endometrial biopsy), menstrual cycle phase, menopausal status and exogenous hormone use; these were used to distinguish genuine biological cell states from groupings reflecting donor- or condition-specific variation rather than independent cell identities. Finally, spatial localisation provided a final validation layer where matched Visium or Xenium spatial transcriptomics data were available, annotated cell types were confirmed to occupy expected tissue compartments by spot deconvolution; for selected populations of particular biological interest or ambiguity, smFISH was used for single-cell resolution spatial validation.

Cell state labels assigned at each level of the hierarchy were projected back onto the main integrated manifold for visualisation and cross-tissue comparison, as described above.

###### Nomenclature and Cell Classification mapping

We developed two complementary nomenclature systems for the human female reproductive tract, in consultation with the Human Cell Atlas Reproduction Network and international teams of organ experts, including developmental biologists, clinicians and pathologist and bioinformaticians.

###### Computationally friendly labels

The first is a computationally oriented cell type labelling system designed for programmatic use. Cell type labels follow a hierarchical structure:

lineage + _ + broad cell type + _ + cell state details.

This enables users to recover broad cell categories by trimming labels at any delimiter “_” and to trace each annotation back to its lineage of origin and broad category.

Immune cells are an exception to this scheme, instead following the simpler structure of lineage + _ + cell state details, and different broad categories are assigned to more closely align with conventional immune cell type nomenclature.

| **Label name** | **Label formula** |
| --- | --- |
| *celltype_HCA_lineage* | lineage |
| *celltype_HCA_broad* | lineage_broadcelltype |
| *celltype_HCA_fine* | Lineage_broadcelltype_cellstate  + _furtherdetails (optional) |

***Supplementary Note 1.1. Table 2. celltype labelling system in the Human Female Reproductive System Cell Atlas v1.***

In the integrated object, these labels are called celltype_HCA_lineage, celltype_HCA_broad and celltype_HCA_fine.

**Human-readable names and descriptions**

Each cell type label in the *Human Female Reproductive System Cell Atlas v1* (aka, *celltype_HCA_fine*) is linked to an entry in our cell type classification, designed to link cell type names with histopathological terminology, the broader scientific literature and Cell Ontology terms.

This cell type classification was organised using a four-level hierarchical framework (L1: lineage, L2: cell broad type, L3: cell fine type and L4: cell state), in which tissue-agnostic identity is defined at the upper L1-L2 levels and additional biological context (such as anatomical location, temporal state or function) is introduced progressively at lower levels L3-L4. For tissue-restricted cell types, such as epithelial populations, lower levels capture anatomical location and cycle-dependent states. For broadly distributed cell types, such as immune and endothelial cells, lower levels instead capture cell state, activation status and functional subtype independently of tissue of origin.

Marker genes supporting each annotation, relevant synonyms from the literature and tissue distributions are provided in **Supplementary Table 2**, together with mappings to equivalent terms used in previously published single-organ atlases. Additionally, for some cell types, we provide notes with extra information about the cell type and alternative population names agreed by the HCA Reproductive Network.

##### Note 1.2. Integration with other organ datasets

###### Integration of reproductive tract-derived NCR^hi^ ILC3s with those from other mucosal tissues.

To understand whether the NCR^hi^ ILC3s exhibited tissue-adapted characteristics in the reproductive tract, we leveraged the availability of scRNA-seq datasets of other mucosal tissues, including the colon, lung, and oral cavity^10–13^. Each individual public dataset was preprocessed analogously to what is described in the **“scRNA-seq atlas construction”** section in **Methods,** then manually re-annotated to identify immune cells (i.e., expressing *PTPRC*) and associated broad cell types.

The innate lymphocytes (NK cells and ILCs) from each dataset were then integrated with those from the Human Female Reproductive System Cell Atlas v1. Integration was performed with scVI^5^ using the top 3,000 highly variable genes per population (Seurat_v3 method^3^), with dataset of origin as batch variable and donor identity and cell cycle phase as covariates (n_layers=2, n_latent=60, dispersion="gene"). Leiden clustering (resolution=1) and UMAP visualisation were applied to the scVI latent representation using scvi-tools^6^ (v[1.4.2]) and scanpy^7^ (v[1.12]). A final round of cell type annotation and cluster-level quality control were performed analogously to what is described in the “*scRNA-seq atlas construction*” section in **Methods**.

#### Supplementary Note 2. Extended results.

##### Note 2.1 Novel insights from postnatal Human Female Reproductive System Cell Atlas v1

This section focuses on findings from the postnatal pan-organ integration. While we include a harmonised fetal counterpart within this atlas to enable cross-stage comparisons, the sections below primarily discuss observations emerging from the postnatal integration.

Pan-organ integration of the fetal female reproductive tract, including external genitalia, which are not covered in the postnatal component of the Human Female Reproductive System Cell Atlas v1, was previously described in Lorenzi et al 2026^14^. Given the limited availability of comparable reproductive tract fetal datasets and the extensive validation of the proposed cell types using matched high-resolution spatial transcriptomics, we refer readers to the original study for detailed characterisation of the cell types and observations in the fetal atlas.

###### Organisation of the epithelium and mesothelium

Integration of epithelial and mesothelial compartments from ovaries to vagina revealed a structured hierarchy of cell types with characteristic tissue distributions, a subset of which were only resolvable through cross-organ analysis (**Figure 4a,b**).

**Mesothelial cells**

Mesothelial cells, defined by expression of *LNRR4* and *UPK3B* and embryologically distinct from conventional epithelia (*EPCAM*^+^, *KRT17*^+^), formed a coherent population spanning the ovarian surface epithelium and the serosal surfaces of the reproductive tract, separating cleanly from all conventional epithelial clusters in integrated space (**Figure 4c**).

**Conventional epithelial cells**

Beyond mesothelial cells, cross-organ integration identified four subtypes of conventional epithelium: mucinous, squamous/basal, ciliated and non-ciliated secretory cells. Each population had a characteristic tissue distribution that broadly reflected the known biology of the organs, but with organ-specific transcriptomic signatures. Notably, the integration revealed novel rare populations in unexpected locations that were only detectable at this pan-organ scale.

Mucinous epithelial cells (*MUC5AC*^+^/*MUC5B*^+^) and squamous basal cells (*TP63*^+^/*KRT5*^+^) were most abundant in the cervix and vagina, consistent with the specialised barrier function of the mechanically and chemically exposed distal reproductive tract (**Figure 4c**). Cross-organ integration resolved the identity of a *MUC5B*^+^ population recurrently observed in endometrial datasets as endocervical mucinous epithelium. High-resolution Xenium imaging confirmed that the full mucinous programme (*MUC5B*^+^/*BPIFB1*^+^/*LTF*^+^) was confined to the endocervix, while a subset of endometrial glands (predominantly in the basalis, but also seen in lumen and functionalis) showed partial activation of this programme (*MUC5B*^+^/*LTF*^+^ but *BPIFB1*-negative), indicating that mucinous identity extends across the reproductive tract with organ-specific depth of programme activation (**Figure 4d**)

Ciliated (*FOXJ1*^+^) and non-ciliated secretory epithelial cells were all sampled in the fallopian tube and endometrium, consistent with their established roles in gamete transport and uterine receptivity, respectively (**Figure 4b**). Both cell types displayed clear organ-specific transcriptional signatures (**Figure 4a**).

**Paratubal inclusion cysts**

Squamous basal cells were identified in the fallopian tubes of n = 9 donors. Spatial transcriptomics localised these cells to *paratubal inclusions*, and their expression of *GATA3* (a marker associated with urothelial differentiation) is consistent with Walthard cell rests, benign transitional cell rests of uncertain significance occasionally identified in the broad ligament and fallopian tube (**Figure 5e,f**). This interpretation was independently confirmed by a specialist gynaecological pathologist.

**Epithelial cells of ovarian inclusion cysts**

Unexpectedly, we identified n = 220 cells in ovarian samples that clustered with conventional ciliated and secretory epithelial populations of the reproductive tract, rather than with ovarian surface epithelium or granulosa cells, in the integrated scRNA-seq space. Marker analysis identified two distinct secretory epithelia within these cells. One population expressed *PAX8* together with *OVGP1*, consistent with secretory fallopian tube epithelium, whereas a second secretory population expressed *PAX8* and *WNT7A*, consistent with endometrial glandular epithelium (**Figure 5a-d**). Spatial transcriptomics subsequently localised these populations to discrete intraovarian inclusion cysts, confirming that these cells represent bona fide tissue-resident structures rather than dissociation or sampling artefacts. Independent histopathological review identified the structure containing secretory *PAX8*⁺/*OVGP1*⁺ and ciliated *PAX8*⁺/*FOXJ1*⁺ in an adult donor (28 years old) as consistent with ovarian cortical inclusion cysts lined by tubal-type epithelium (**Figure 5a,b**). In contrast, the the structure containing *PAX8*⁺/*WNT7A*⁺ epithelium in a paediatric donor (14 years old) was interpreted as ovarian endometriosis. Supporting this interpretation, spatial transcriptomics deconvolution showed that the stromal compartment surrounding the *WNT7A*⁺ glands expressed *HOXA9* and *HOXA10*, transcription factors characteristic of endometrial-type fibroblasts, consistent with the presence of endometrial-type stroma alongside the glandular epithelium (**Figure 5c,d**).

These observations are relevant in the context of ovarian pathology. To our knowledge, this atlas provides the first single-cell transcriptomic characterisation of ovarian cortical inclusion cysts and ovarian endometriosis in donors without a prior diagnosis of either condition, establishing a molecular reference for ectopic epithelial populations in the otherwise healthy ovary. Altogether, these findings demonstrate how pan-reproductive single-cell integration can reveal rare epithelial populations linked to early pathological processes that would likely remain undetected in single-tissue analyses.

###### Organisation of the mesenchyme

Cross-organ integration of postnatal and adult mesenchymal cells resolved interstitial fibroblasts alongside three additional transcriptionally distinct classes (adventitial fibroblasts, perivascular mural cells and smooth muscle cells) each with characteristic marker profiles and tissue distributions not resolved in prior single-tissue analyses (**Figure 2**).

**Interstitial stromal fibroblasts**

Interstitial stromal fibroblasts were the most abundant mesenchymal population in the ovary, endometrium and cervix, consistent with the predominantly fibroblastic structure of these tissues. In contrast, their relative abundance was lower in the fallopian tube, whole uterus and vagina, where smooth muscle cells and adventitial fibroblasts contribute more substantially to the mesenchymal compartment (**Figure 2**). Across tissues, interstitial fibroblasts retained distinct organ-specific transcriptional signatures, including developmental HOX patterns, indicating stable positional identity in adult tissues.

Annotation of interstitial stromal fibroblasts presents specific challenges in dissociated scRNA-seq data. Previous studies in the endometrium and ovary have shown that fibroblast subpopulations occupy distinct anatomical niches that are difficult to capture after dissociation and conventional scRNA-seq analysis, but are readily observed in intact tissue by spatial transcriptomics. Examples include the basalis and functionalis layers of the endometrium, and the outer cortex, inner cortex, medulla and perifollicular regions of the ovary. Fibroblasts are particularly sensitive to tissue dissociation: disruption of extracellular matrix interactions during enzymatic digestion induces stress and inflammatory gene programmes, leading to measurable transcriptional changes relative to their in situ state. In addition, as the dominant population, these contribute disproportionately to overall transcriptional variation and often shape the structure of the integrated embedding. As a result, most clusters reflect dissociation-induced signatures (like stress, inflammatory and programmed cell death) rather than spatially or functionally defined fibroblast states.

Altogether, the diversity of fibroblasts is more clearly resolved in spatial transcriptomics data than in dissociated scRNA-seq. Consistent with their abundance, fibroblasts dominate the spatial patterns recovered by classical clustering of Visium spots, or Xenium or Visium HD cells, making spatial approaches more informative for resolving their organisation and diversity. Here, the availability of matched spatial transcriptomics across organs enabled the annotation and spatial localisation of interstitial fibroblast subpopulations, providing a spatially grounded reference that can be used to orient and interpret fibroblast clusters in newly generated single-cell datasets from these tissues.

**Adventitial fibroblasts**

Cross-organ integration revealed a population of adventitial fibroblasts spanning the ovary, fallopian tube, uterus, cervix and vagina that had not been recognised as a shared compartment in prior single-tissue analyses, where organ-enriched populations were often annotated as stromal fibroblasts.

This pan-reproductive adventitial fibroblast population was defined by co-expression of *DPT*, *SFRP2* and *C3* (**Figure 2c**), a module absent from interstitial fibroblasts, perivascular cells and smooth muscle cells, which unified otherwise heterogeneous, organ-specific populations into a coherent compartment. Within this compartment, *PI16* and *C7* were mutually exclusive, defining a gradient from a *PI16*^hi^/*CD34*^+^/*C7*^−^ to a *PI16*^−^/*CD34*^–^/*C7*^hi^ state, with *COL15A1* increasing modestly along the same axis. This organisation is consistent with the cross-tissue fibroblast hierarchy defined in mouse, in which *PI16*^+^ and *COL15A1*^+^ mark two universal fibroblast subtypes, with *PI16*^+^ cells positioned as the root progenitor state from which more specialised identities emerge^15^.

Adventitial fibroblasts were most abundant in the fallopian tube and vagina, consistent with the tissue organisation of these organs. The *PI16*^hi^ state was enriched in the fallopian tube, whereas the remaining tissues were dominated by *PI16*^lo^/*C7*^hi^ states (**Extended Data Figure 3b**). The *C7*^hi^ populations from each organ correspond to fibroblast populations previously described in single-tissue analyses: the *C7*^+^ fibroblast of the endometrium^16^, spatially mapped to the basalis fibroblasts^1^; the *C3*^+^ fibroblasts of the paediatric ovary, localised to perivascular and subepithelial niches beneath the ovarian surface epithelium (Garcia-Alonso 2026; under review; preprint bioRxiv) and the population labelled as fibroblasts in the fallopian tube by Ulrich et al^17^ (**Extended Figure 3**).

The functional role of these populations remains to be fully established. The pan-adventitial expression of *C3* is consistent with complement-competent, immune-modulatory fibroblasts described in pulmonary vascular contexts^18^, while *SFRP2* expression implicates these cells in local WNT signalling modulation at the adventitial niches. Spatial transcriptomics deconvolution confirmed that these adventitial fibroblasts (*DPT*^+^, *SFRP2*^+^) occupy perivascular and subepithelial positions, with organ-specific distributions: in the uterus, this population was restricted to the basalis and myometrium and excluded from the functionalis; in the ovary, cells localised beneath the surface epithelium and around cortical vessels; and in the fallopian tube, adventitial *PI16*^hi^ fibroblasts were localised to the subserosal connective tissue; while the adventitial *PI16*^lo^ where in the interstitium of the mucosa and muscularis (**Figure 2d**).

**Perivascular fibroblasts**

We next defined the perivascular compartment, marked by the absence of *DPT*, *C7* and *PI16* expression (**Figure 2c**), as a heterogeneous group that ranges from highly contractile vascular smooth muscle cells (vSMCs; *RERGL*^+^) surrounding larger vessels, to capillary-associated pericytes (*RGS5*^+^) that play a crucial role in supporting the microvasculature.

Within the vSMC compartment, we identified two subpopulations corresponding to the tunica media of large vessels and small arteries. Large vessels vSMCs were present in all tissues except the endometrium and ovarian cortex, whereas small artery/arteriolar vSMCs were detected in all tissues (**Extended Data Figure 3b,c**).

Within the pericytes compartment, general capillary pericytes (*RGS5*^+^, *STEAP4*^+^) were present across all tissues and formed a transcriptionally coherent population, while exhibiting organ-specific transcription factor signatures: *GATA4* in the ovary, *GATA2* and *HOXA9* in the fallopian tube, and *HOXA10*/*HOXA11* in the uterus. These signatures mirror those of the surrounding stromal compartments, suggesting that pericyte transcriptional identity reflects that of their tissue location.

In addition, we identified a distinct perivascular population displaying features intermediate between pericytes and vSMCs, co-expressing canonical smooth muscle markers (*MYH11*^+^, *CNN1*^+^) together with pericyte-associated genes such as *STEAP4*. This population also expressed unique markers like *CLSTN2*, involved in calcium signaling, as previously described by Barnett et al.^19^.

**Uterine-specific perivascular populations**

Profiling of both endometrial and myometrial compartments resolved three uterine pericyte (*RGS5*^+^) populations consistent with previous work^1^. Pan-organ integration further showed that two of these populations were uterus-specific and were not detected in other reproductive tissues.

The first was a pericyte population enriched in the endometrium, but also present in the myometrium, that we called “*pericyte of the uterine spiral arteries*" (short label Mesen_Pericyte_EndoSpiralArt). Beyond canonical pericyte markers, this population uniquely expressed the markers *SLC38A11* and *STC2* alongside other hypoxia-induced genes (*FLT1*) and genes associated with stiffness (*AOC3*), contractility (*MYOM1*, *CTNNA3*), and SMCs (*ACTG2*, *KCNMB1*; **Figure 2c**), while lacking expression of *RERGL* (characteristic of vSMCs). Spatial transcriptomics confirmed that *SLC38A11*^+^ cells co-localised with arterial endothelial cells (*SEMA3G*^+^/*CDH5*^+^) in both the endometrium and myometrium; histological inspection of matched H&E sections identified the corresponding vessel structures as spiral arteries, supporting the identity of this population as spiral artery pericytes (**Figure 2d**).

The second uterine-specific population was restricted to the endometrium and not detected in myometrium, which we called “*endometrial stromal-like pericytes*”. It co-expressed the pericyte marker *RGS5* and endometrial interstitial stromal markers *PDGFRA* and *MMP11*, was enriched in the proliferative phase while also present in the secretory phase, and was absent from postmenopausal samples, consistent with dependence on hormone-driven endometrial regeneration. This population occupied an intermediate position between perivascular and stromal clusters in the integrated scRNA-seq manifold (**Figure 2a,b**) and localised to the perivascular regions between the pericytes of the uterine spiral arteries and the interstitial stroma^20^, arguing against a pericyte-stromal doublet origin. We interpret this population as a perivascular-stromal transitional state.

Previous studies used *SUSD2* to enrich clonogenic endometrial stromal cells, usually referred as to endometrial mesenchymal stem/stromal cells (eMSCs), which have been proposed to replenish the endometrial interstitial stromal fibroblast compartment^21^. In our atlas, *SUSD2* was expressed across all uterine perivascular populations, with highest expression in spiral artery-associated pericytes (**Figure 2c**). This suggests that the *SUSD2*^+^ fraction enriched in previous studies may capture a broader uterine perivascular compartment than previously appreciated, and that the clonogenic cells identified in those studies may reside within these transcriptionally distinct perivascular populations.

Barnett et al.^19^ identified uterine-specific perivascular transcriptional signature (including *APCDD1* and *HOPX*) by comparing mural cells across 19 human organs. In that study, the uterus was the only reproductive organ included. Cross-referencing their uterine perivascular signature with our atlas reveals that the same markers are expressed across perivascular cells of the fallopian tube, uterus, cervix and vagina (spanning canonical pericytes, vSMCs, and the two uterine-specific pericyte populations described here). This suggests that what Barnett et al. identified as uterine-specific perivascular signature in a broad cross-body comparison may more accurately reflect a shared perivascular identity across the female reproductive tract.

**SMCs**

The integration also revealed a muscularis SMC population distinct from vascular vSMCs, characterised by the absence of *RERGL* and *KCNAB1* expression and enrichment of contractile organ-wall markers (e.g. *ACTG2*, *DES*). These cells were localised to the muscularis layers of the fallopian tube, vagina, myometrium and ovarian medulla, and were absent from the endometrium and ovarian cortex, consistent with an organ-wall contractile role rather than a vascular identity.

###### Organisation of the vascular endothelial cells

The endothelial compartment resolved into 11 populations spanning arterial, venous, capillary and lymphatic lineages, consistent with the vascular organisation described in cross-body atlases. Arterial endothelial cells (*GJA5*^+^/*GJA4*^+^/*SEMA3G*^+^) were detected across all reproductive tissues. Venous endothelial cells showed greater diversity and included canonical venous cells, post-capillary venous endothelial cells (*ACKR1*^+^/*SELP*^+^/*TNFRSF1A*^+^), activated post-capillary venous endothelial cells (*SELE*^+^/*ICAM1*^+^/*AQP1*^+^/*PLVAP*^+^), and high endothelial venule cells (*TIMP1*^+^/*OSBPL1A*^+^).

Capillary endothelial cells comprised general capillaries (*RGCC*^+^/*CAV1*^+^/*VWF*^+^), angiogenic tip cells (*ESM1*^+^/*CXCR4*^+^/*PGF*^+^) and cycling capillaries (*MKI67*^+^/*CDK1*^+^), indicating ongoing vascular turnover. We also identified an endometrium-enriched capillary endothelial population marked by *APCDD1*^+^, a negative regulator of WNT signalling. This population aligns with the organotypic uterine vascular signature described by Barnett et al.^19^. However, comparison across the reproductive tract showed that *APCDD1* expression was not restricted to the uterus, as it was also present in the cervix. Lymphatic endothelial cells (*PROX1*^+^/*LYVE1*^+^/*CCL21*^+^) were detected across tissues.

Thus, the reproductive endothelial atlas recapitulates the major vascular bed identities described across the body. These include immune-interactive venous states, cycling and angiogenic capillary populations, and an *APCDD1*-associated vascular programme shared across the uterus-cervix but absent from the ovary and fallopian tubes.

###### Organisation of the immune cells

Cross-organ integration of prenatal, postnatal and adult immune cells from individuals without exogenous hormone treatment resolved multiple lymphoid, myeloid, and progenitor cell states across 13 broad compartments (**Extended Data Fig. 4a**): haematopoietic progenitor cells (HPC), myeloid progenitors, megakaryocytes/platelets, erythroid cells, granulocytes, monocytes, macrophages, conventional and non-conventional dendritic cells (DC), B cells, T cells, innate lymphoid cells (ILC) and natural killer (NK) cells. We defined specific markers for all populations (**Extended Data Fig. 4b**), and several had tissue-residency characteristics not appreciated in prior single-organ analyses (**Figure 3b** and **Extended Data Fig. 4c**), as described in the main text. A summary of cell taxonomy and comprehensive markers per cell state is presented in **Supplementary Table 2.**

**Lymphocytes**

Integrating immune cells across human development enabled the detection of prenatal-specific cell states spanning B lymphopoiesis, including pro-B cells, large pre-B cells, small pre-B cells and immature B cells (**Figure 3a** and **Extended Data Fig. 4a,b**). Naive and memory B cells were predominantly derived from adult samples (**Extended Data Fig. 4a**). Plasma cells separated on the basis of κ- or λ-restricted subsets (**Figure 3a** and **Extended Data Fig. 4b**).

The T cell compartment was also primarily derived from adult samples, with the exception of a Vγ9/Vδ2^+^ γδT cell population specific to the fetal gonads (**Figure 3a** and **Extended Data Fig. 4a**). The remaining T cell populations comprised CD8^+^ T cells, CD4^+^ T cells and innate-like T cells. CD8^+^ T cells included naive, effector memory (T_EM_) and tissue-resident memory (T_RM_) subsets, with two T_RM_ states distinguished on the basis of *GZMK* expression (**Extended Data Fig. 4b**). Marker expression of CD8^+^ T_EM_ cells is consistent with terminally differentiated effector CD45RA^+^ T_EMRA_ cells^22^ (**Supplementary Table 2b**), however we have refrained from annotating them as such due to the limitations of scRNA-seq in profiling CD45 isoforms. CD4^+^ T cells primarily comprised naive, regulatory (T_reg_) and T_H17_ subsets, including a stem cell memory-like T_H17_ state, which expressed both naive and memory T cell markers^23^. The innate-like T cell compartment consisted of MAIT cells, NKT and Vδ1^+^ γδT cells. Modular nomenclature for all T cell subsets, consistent with the recent consensus statement published by Masopust et al.^22^, are presented in **Supplementary Table 2b.**

NK cells were predominantly derived from adult samples and comprised three main subsets on the basis of *CD56* (*NCAM1*) and *CD16* expression (**Figure 3a** and **Extended Data Fig. 4a,b**). CD56^hi^ NK cells were enriched in the endometrium and exhibited three distinct states in line with what we have previously described^1,24^: uNK1 (*B4GALNT1*^hi^, *KIR2DL1*^hi^), uNK2 (*XCL1*^hi^, *CDHR1*^hi^) and uNK3 (*CD160*^hi^, *LDB2*^hi^). CD16^hi^ (and *CD56*^dim^) NK cells were detected across all reproductive tissues regardless of developmental stage (**Extended Data Fig. 4c**), whereas a rare CD56^dim^/CD16^dim^ circulating (*SELL*^hi^) NK subset upregulating *SPTSSB*, a regulator of sphingolipid biosynthesis, was only found in the adult reproductive tract (**Extended Data Fig. 4c**). We also identified two subsets of ILC3 enriched in the uterus: NCR^hi^ and NCR^lo^, as expanded upon in the main text.

**Myeloid**

Among the myeloid cells, monocytes (*CDA*^hi^) and macrophages (*C1QB*^hi^) were the most prevalent (**Extended Data Fig. 4a,b**). Neutrophils, basophils/eosinophils and mast cells were enriched in postnatal samples (**Figure 3a** and **Extended Data Fig. 4a**). We also identified multiple subsets of dendritic cells (DCs) (**Figure 3a**), including cDC1, cDC2, CD207^hi^ cDC2, plasmacytoid DC (pDC), AXL^+^/SIGLEC^+^ DC (ASDC)^25^, mature regulatory DC (mregDC)^26^ and tolerizing DC (tolDC)^27^. To enable the identification of cell states, monocytes, macrophages and cDC2 subsets were clustered and re-annotated separately.

We detected classical (*CD14*^hi^, *FCGR3A*^lo^) and non-classical (*FCGR3A*^hi^, *CD14*^lo^) monocytes across nearly all organs (**Extended Data Fig. 5d,f** and **Extended Data Fig. 4c**), alongside a broadly distributed macrophage population that shares expression of certain monocyte-associated markers (i.e., *S100A9*) but lacks others (i.e., *S100A12*) (**Extended Data Fig. 4b**). These macrophages did not express markers associated with more differentiated macrophages subsets (i.e., *LYVE1*, *FCGBP*) or inflammatory markers (i.e., *EREG*), and were therefore named “*transitional*” macrophages. Prenatal reproductive tract-derived macrophages, besides those that expressed *LYVE1*, were present in this subset. The remaining macrophage subsets separated on the basis of *LYVE1* expression, where LYVE1^lo^ subsets included uterine and fallopian tube lipid-associated macrophages (uftLAMs), ovary LAMs (oLAMs) and inflammatory uterine macrophages (called “*uMac_Inf*”). uftLAMs and oLAMs express markers consistent with a canonical lipid-associated macrophage (LAM) programme (*TREM2*, *CD9*, *SPP1*) and lipid metabolism signature (*ALDH2*, *APOE*, *APOC1*, *APOC2*, *LPL*), and were characterised further in the main text. *uMac_Inf* were characterised in a parallel study (Lorenzi et al. 2026; under review; preprint in bioRxiv).

**LYVE1^hi^ macrophages**

LYVE1^hi^ macrophages, which were shared across all organs (**Extended Data Fig. 4c**), expressed a suite of genes consistent with a perivascular identity (**Extended Data Fig. 5e**). These included *LYVE1* and *CEMIP2*, together regulate the ECM niche: LYVE1 binds hyaluronan, a glycosaminoglycan enriched in the perivascular niche^28^, while CEMIP2 mediates its degradation^29^. Additional markers also suggest perivascular residency, including *ANTXR2*, which contributes to extracellular matrix remodelling in the mouse reproductive tract^30^, and *ITGA9*, which pairs with β1 to form the α9β1 integrin heterodimer, serving as a receptor for VEGF-C and VEGF-D^31^.

Beyond their putative perivascular residency, these macrophages actively promote angiogenesis and tissue homeostasis through expression of several genes. *NRP1* encodes a co-receptor for VEGF ligands, through which *NRP1*^+^ macrophages promote neovascularisation in the mouse central nervous system, adipose tissue, and tumours^32–34^; *PDGFC* encodes a secreted ligand for PDGFR-α expressed by pericytes and stromal fibroblasts (**Figure 2c**), a signalling axis that drives alveogenesis in mice^35^; *GAS6* encodes a ligand for TAM receptors TYRO3, AXL and MERTK, promoting survival and efferocytosis; and *ENPP2* encodes an enzyme that produces lysophosphatidic acid, which exhibits pro-angiogenic effects in mice^36^. LYVE1^hi^ macrophages also express additional markers characteristic of anti-inflammatory, homeostatic macrophages, including *MRC1* (**Figure 3d**).

###### Organisation of the peripheral nervous system cells

The peripheral nervous system (PNS) compartment was dominated by non-myelinating Schwann cells, with myelinating Schwann cells comprising a smaller fraction (**Figure 1e**). Both populations were detected in the ovary, fallopian tube and uterus, and were most abundant in the vagina, consistent with its dense autonomic and sensory innervation. PNS cells were not detected in the cervix, likely reflecting limited cell sampling depth rather than a true absence of innervation.

###### Organisation of ovarian follicular cells

The atlas captures the three core follicular cell compartments of the ovary (germ cells, granulosa cells and theca cells) drawing primarily from the paediatric ovarian dataset of Garcia-Alonso et al. (2026, under review, preprint in bioRxiv), which specifically targeted the ovarian cortex where primordial and early growing follicles are enriched. Theca cells were predominantly contributed by the dataset of Guahmich et al.^9^, whose sampling strategy focused on growing follicles and enabled deeper recovery of these populations than is typically achieved from standard cortical biopsies. Datasets profiling whole ovaries from aged adult donors contributed relatively few follicular cells, reflecting both the age-related depletion of the primordial follicle pool and the dilution of cortical regions within the full ovarian volume.

Germ cells were identified as oocytes associated predominantly with primordial to preantral follicles, defined by expression of *ZP3*, *GDF9* and *DDX4*, and absence of *BMP15*, a marker of more advanced antral-stage oocytes. Consistent with the progressive decline of the follicle pool across the reproductive lifespan, oocytes were sparse in adult samples and most abundant in paediatric tissue.

Granulosa cells spanned a continuum of transcriptional states across early follicle development, as defined by Garcia-Alonso et al. (2026; under review, preprint in bioRxiv). These included AMH^–^ granulosa in primordial and transitioning follicles (*RDH10*⁺/*WNT6*⁺/*AMH*^–^), *AMH*^+^ granulosa in primary follicles (*RDH10*⁺/*WNT6*⁺/*PTPRZ1*⁺/*AMH*⁺), *AMH*^+^ granulosa non steroidogenic in small multilayered follicles (*AMH*⁺/*GSTA1*⁺), *AMH*^+^ granulosa in multilayered follicles emerging steroidogenic activity (*AMH*⁺/*CYP19A1*⁺/*IHH*⁺). We also identified a population of *AMH*^+^ granulosa that is cycling (*MKI67*⁺) and two atretic granulosa populations downregulating housekeeping genes.

Theca cells comprised three transcriptionally and functionally distinct populations associated with growing follicles, originally defined by Guahmich et al.^9^. Perifollicular theca interna (*THBD*⁺/*PIEZO2*⁺) localised adjacent to the granulosa layer; androgenic theca interna (*CYP17A1*⁺/*ANPEP*⁺) exhibited a steroidogenic programme; and theca externa (*PTCH1*⁺/*ACTA2*^hi^) formed a contractile outer layer. These populations were primarily associated with preantral and antral follicles, as well as atretic remnants, consistent with the emergence of theca cells during follicle growth beyond the primordial stage.

For a detailed characterisation of follicular cell states and their spatial organisation within the ovarian cortex, we refer to Garcia-Alonso et al. (2026; under review, preprint in bioRxiv).

##### Note 2.2. Expression of steroid receptors along all the cell types of the female reproductive HCA

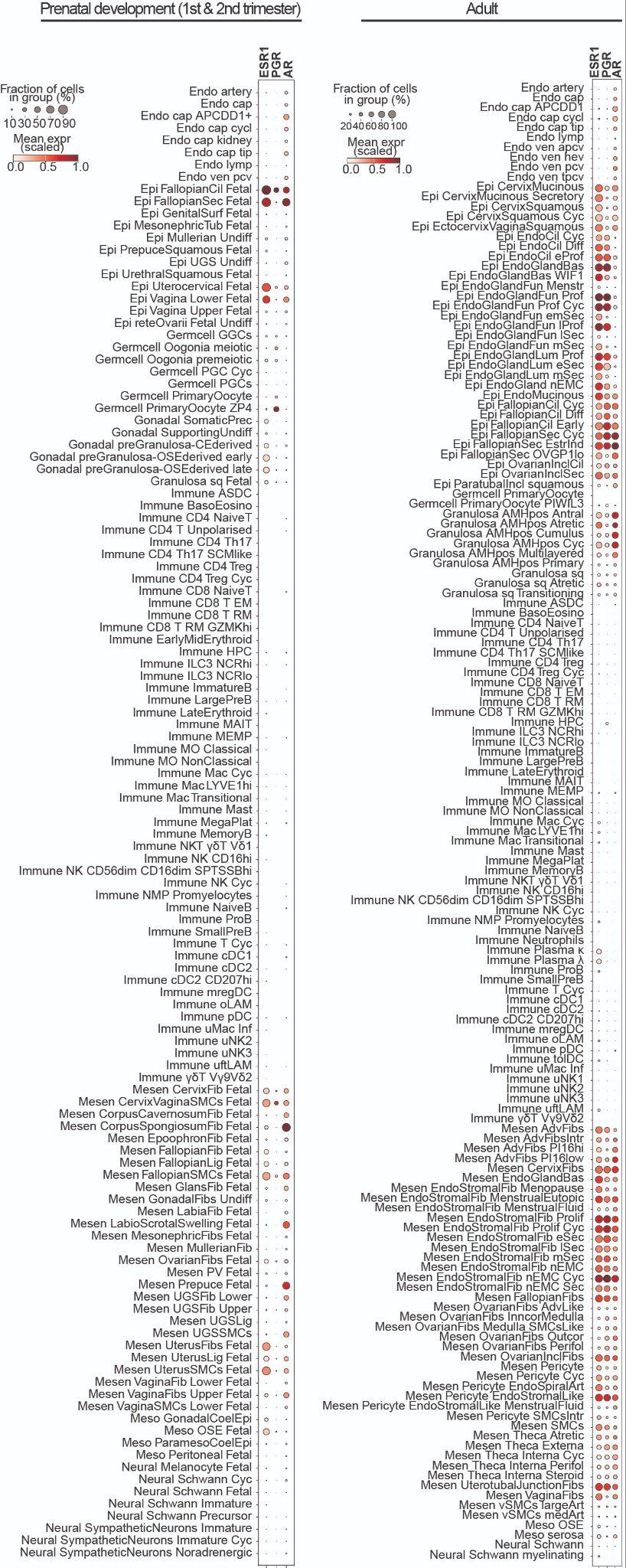

***Supplementary Note 2.2. Table 1. Gene expression distribution of steroid receptors in the Human Female Reproductive System Cell Atlas v1*** *Dot plot showing log-transformed, min–max-normalised expression of selected marker genes (x-axis) across all HCA v1 cell type categories (y-axis). Labels correspond to HCA fine_celltype.*

##### Note 2.3. GWAS integration with the Human Female Reproductive System Cell Atlas v1

We computed cell type enrichment using S-LDSC and identified genes driving each association using our leave-one-out framework.

Age at menopause heritability was enriched in germ cells in the ovaries **(Figure 6a** and **Extended Data Fig. 7a, Supplementary Table 4b)**, consistent with the role of the ovarian reserve in determining menopause onset. This was driven by DNA damage repair genes **(Extended Data Fig. 7b, Supplementary Table 4d)**, consistent with oocyte maintenance to stabilise the ovarian reserve.

Uterine fibroids heritability was enriched in smooth muscle cells **(Figure 6a, Supplementary Table 4b)**. Hormone receptors (*ESR1*, *PGR, THRB, PRLR*) appeared across multiple pathways to drive the enrichment in uterine smooth muscle cells **(Extended Data Fig. 7b, Supplementary Table 4d)**. Top pathways included chromatin and transcriptional machinery, including major transcription factors involved in cell fate and development (*ALX3*, *SOX15*, *BNC2*, *FOXP2*) and genes directly involved in chromatin remodelling (*E2F6*, *PCGF6*), suggesting that transcriptional rewiring through chromatin remodelling is a major driver of uterine fibroids in smooth muscle cells. Whilst chromatin alterations have previously been observed in uterine fibroids^37–39^, these results support that genes involved in chromatin remodelling drive uterine fibroids in smooth muscle cells. We also find tissue remodelling (*TGFBR3*, *FOXO1*) and lipid metabolism-associated genes (*ACAT1, PCK1, DGKB*), consistent with tumour growth and previous associations suggested between lipid levels and uterine fibroids^40,41^. Endometrial stromal fibroblasts also showed broad suggestive enrichment, supporting their potential role in uterine fibroid pathogenesis^42^.

Heavy menstrual bleeding heritability was enriched in basal endometrial fibroblasts, detailed in the main text. To check that our ΔZ scores per gene were not biased by gene size, we computed the Spearman’s rank correlation between ΔZ and gene size for the heavy menstrual bleeding in basal fibroblasts ΔZ scores – given the extent of our follow up on this trait in the main text – and did not find a significant correlation (⍴=-0.014, *p*=0.32). Here we detail basal-fibroblast-specific genes driving enrichment. BMP2 regulates endometrial wound healing by promoting re-epithelialisation and induces expression of the transcription factor KLF10 (TIEG1)^43^, which modulates cell proliferation, is induced by estrogen and impaired by glucocorticoids^44^ and inhibits proliferation of cultured ectopic endometrial stromal cells^45^. CCDC117 drives proliferation and DNA synthesis in the developing heart^46^ and DNAJA1 (Hdj2) knockdown upregulates multiple matrix metalloproteinases (MMPs) in glioblastoma cell culture^47^. MMPs were also among genes driving the trait–cell type enrichment **(Extended Data Fig. 7b, Supplementary Table 4d)**, though their expression is not specific to basal fibroblasts. Given this, *DNAJA1* may regulate MMP expression to limit tissue breakdown in the basalis, thereby preventing excessive shedding – a process that, when dysregulated, could contribute to heavy menstrual bleeding. Other specific genes, including PI3K/Akt pathway activators *PIKB**^48^*, *SNCA**^49^*, *SERPINE3**^50^* and *KLF16**^51^*, also showed some albeit weaker genetic signal reflective of the polygenic architecture of HMB **(Extended Data Fig. 7c)**. Collectively, these signals suggest that regulation of steroid-responsive cell proliferation and MMP expression may contribute to the specific role of basal stromal fibroblasts in HMB. HMB heritability enrichment in basal fibroblasts is therefore driven by the intersection of pathways relevant across stromal fibroblasts and a cell-type specific polygenic signal, pointing to multiple genes with individually weaker signal which we may be underpowered to detect in current GWAS.

##### Note 2.4. snATAC-seq Human Female Reproductive System Cell Atlas

We integrated the scATAC-seq and scRNA-seq atlases to annotate cells and generate pseudo-scRNA-seq expression profiles per cell. Cell type annotations were derived from our transcriptomic cell type definitions established in the single-cell transcriptomics HCA. Gene activity scores derived from chromatin accessibility in our cell types were consistent with transcriptomics-derived markers **(Extended Data Fig. 8a-c)** and we recapitulated subtle cell state changes, notably in endometrial cells across the menstrual cycle **(Extended Data Fig. 8d)**.

We called peaks and describe enrichment for ENCODE functional annotations in the main text **(Figure 7b)**. Called peaks were enriched to a lesser extent in less functionally relevant annotations including the intersection of chromatin accessibility and transcription factor binding motifs (CA-TF), but depleted of these annotations alone. Removing peaks overlapping gene bodies increased enrichment in distal annotations but decreased enrichment in proximal annotations, likely from removing promoter-associated peaks with some gene body overlap. Together this suggests that our peaks recapitulate known functional annotations derived from multiple regulatory ENCODE annotations whilst being distinct from ENCODE scATAC-seq peaks, likely due to higher tissue and cell type specificity.

When assessing enrichment of peaks linked to genes for functional annotations compared with unlinked peaks, we found that enrichment increased for distal enhancer-like signatures (dELS) after removing peaks linked in gene bodies **(Extended Data Fig. 8d)**, which could be a consequence of transcription rather than indicating a regulatory relationship. Together with enrichment described in the main text, these results demonstrate that the identified peak–gene links reflect genuine cis-regulatory relationships.

### Supplementary Notes References
