## Extended Data Figures for "An integrated multimodal pan-organ atlas of the female reproductive system across the lifespan contextualises gynaecological pathologies"

Extended Data Figure 1

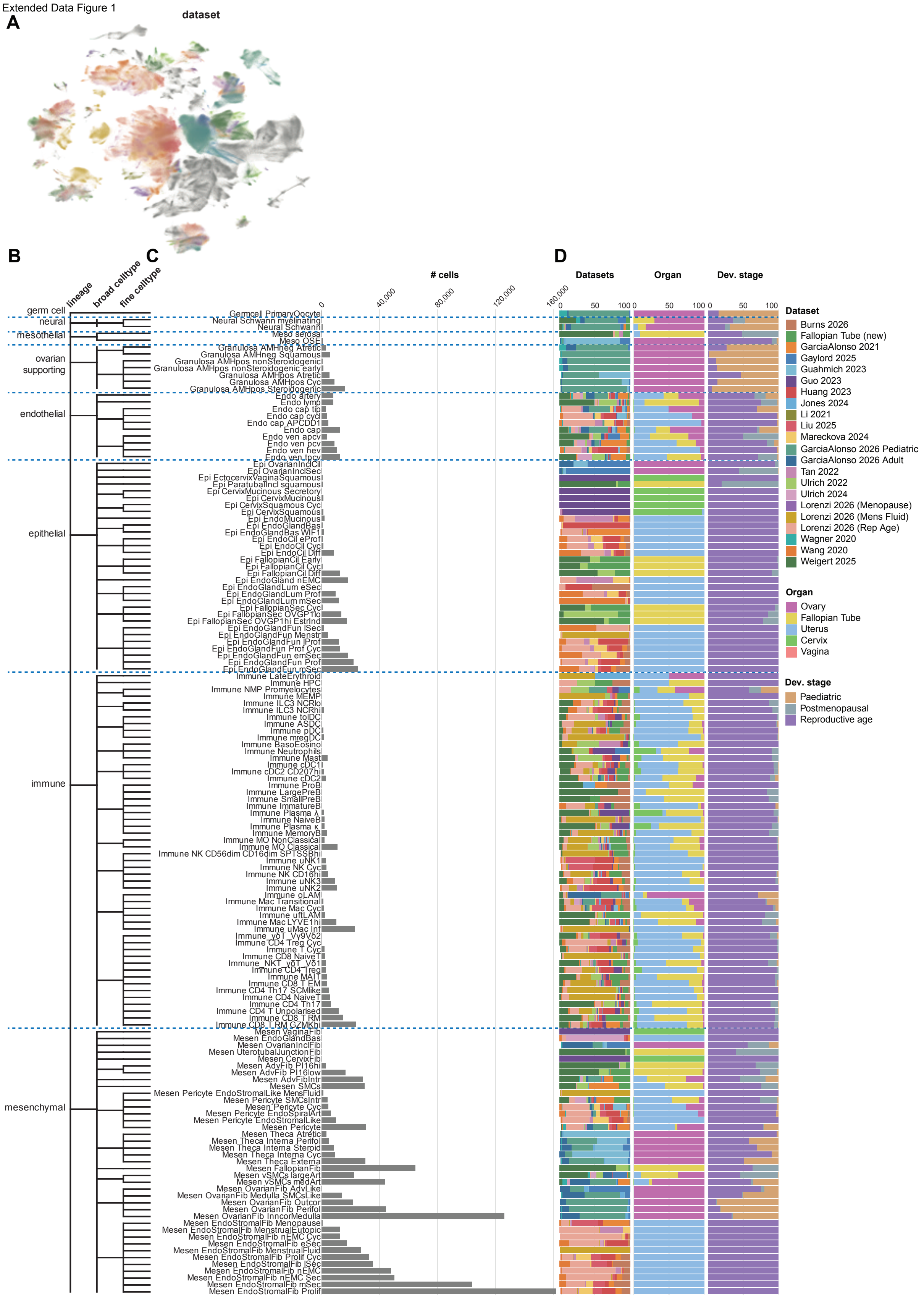

**Extended Data Fig. 1. Fine cell type taxonomy and metadata composition of the Human Female Reproductive System Cell Atlas v1 in the postnatal (pediatric & adult) dataset. A,** Batch-corrected Uniform Manifold Approximation and Projection (UMAP) visualisation manifold of the scRNA-seq dataset (n = 2,235,448 cells; n = 291 donors), coloured by dataset related to Figure 1c-e. Datasets color legend depicted in D. **B,** Dendrogram showing the three-level cell type hierarchy encoded in the `fine_celltype` annotation labels (structured label encoding `lineage_broad_fine`). Lineages are shown on the left, broad cell types in the middle and fine cell types as leaves on the right. Fine cell types are ordered by lineage and ranked by relative abundance within each broad cell type. Each fine cell type is assigned to its broad cell type, and each broad cell type to its lineage, based on the components of the structured label. **C,** Total number of cells per `fine_celltype` in the Human Female Reproductive System Cell Atlas v1. **D,** Bar plots showing, for each `fine_celltype`, the proportional contribution of each dataset (n = 22 datasets; left barplot); contribution of each organ of origin (ovary, fallopian tube, uterus, cervix, vagina; middle barplot); and contribution of developmental stage at time of sample collection: paediatric (postpubertal), reproductive age (adult), and postmenopausal (including perimenopausal). Prenatal-specific cell labels are shown in Extended Data Fig. 2. Please refer to **Supplementary Table 2b** for all cell annotation-associated abbreviations. Dev, developmental.

Extended Data Figure 2

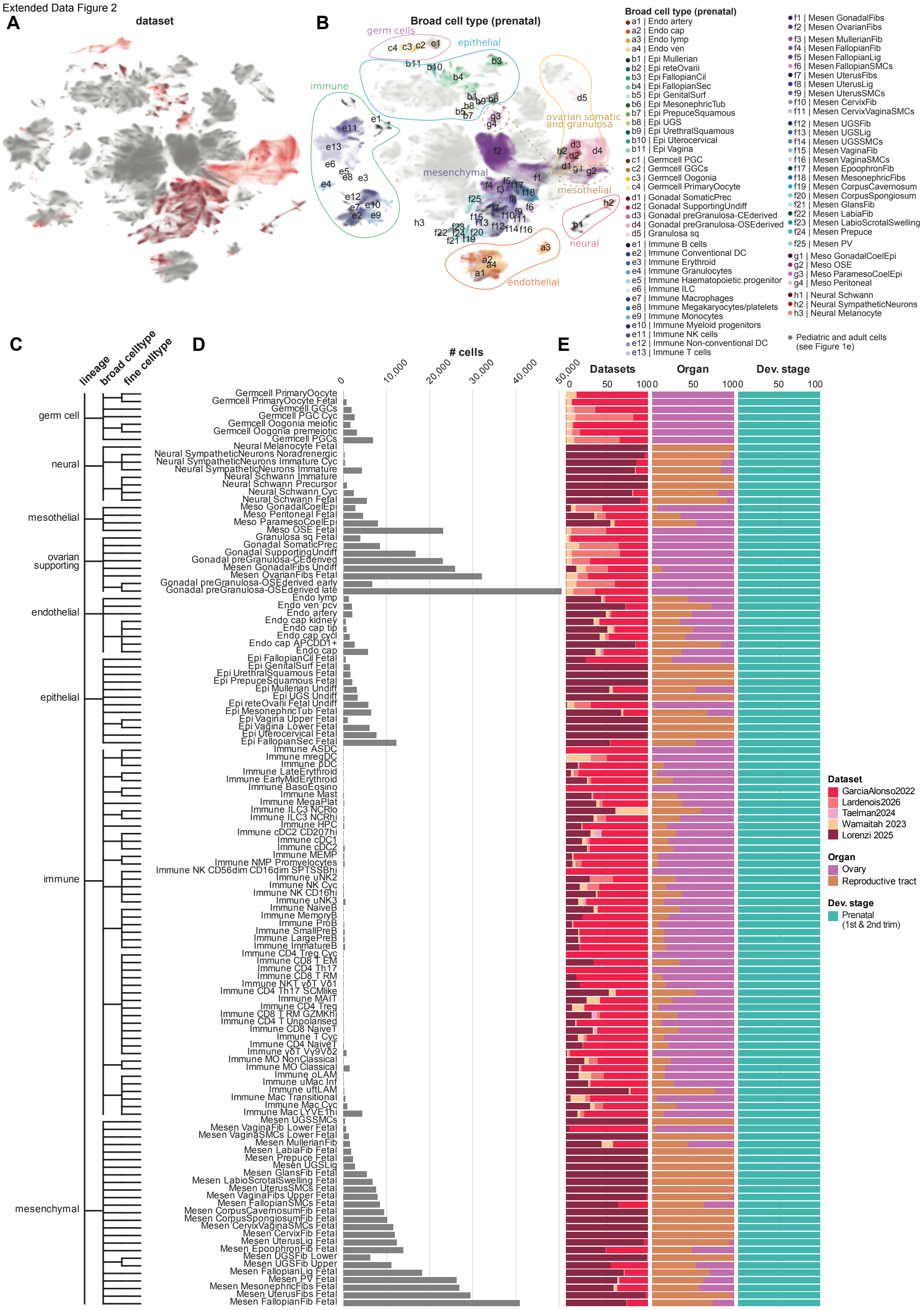


**Extended Data Fig. 2. Fine cell type taxonomy and metadata composition of the Human Female Reproductive System Cell Atlas v1 in the prenatal dataset. A,** Batch-corrected UMAP visualisation manifold of the scRNA-seq dataset (n = 2,235,448 cells; n = 291 donors), coloured by prenatal dataset, related to Figure 1c-e. Dataset color legend depicted in E. **B.**

Batch-corrected UMAP visualisation manifold of the scRNA-seq dataset, coloured by `broad_celltype` annotation in prenatal donors. Postnatal `broad_celltype` categories are shown in Figure 1e. **C.** Dendrogram showing the three-level cell type hierarchy encoded in the `fine_celltype` annotation labels (structured label encoding `lineage_broad_fine`). Lineages are shown on the left, broad cell types in the middle and fine cell types as leaves on the right. Fine cell types are ordered by lineage and ranked by relative abundance within each broad cell type. Each fine cell type is assigned to its broad cell type, and each broad cell type to its lineage, based on the components of the structured label. **D.** Total number of cells per `fine_celltype` in the Human Female Reproductive System Cell Atlas v1, prenatal dataset subset. **E.** Bar plots showing, for each `fine_celltype`, the proportional contribution of each dataset (n = 4 datasets; left bar plot); organ of origin (ovary, or reproductive tract; middle bar plot); and developmental stage at sample collection (all prenatal; right bar plot). Please refer to **Supplementary Table 2b** for all cell annotation-associated abbreviations. Dev, developmental.

Extended Data Figure 3

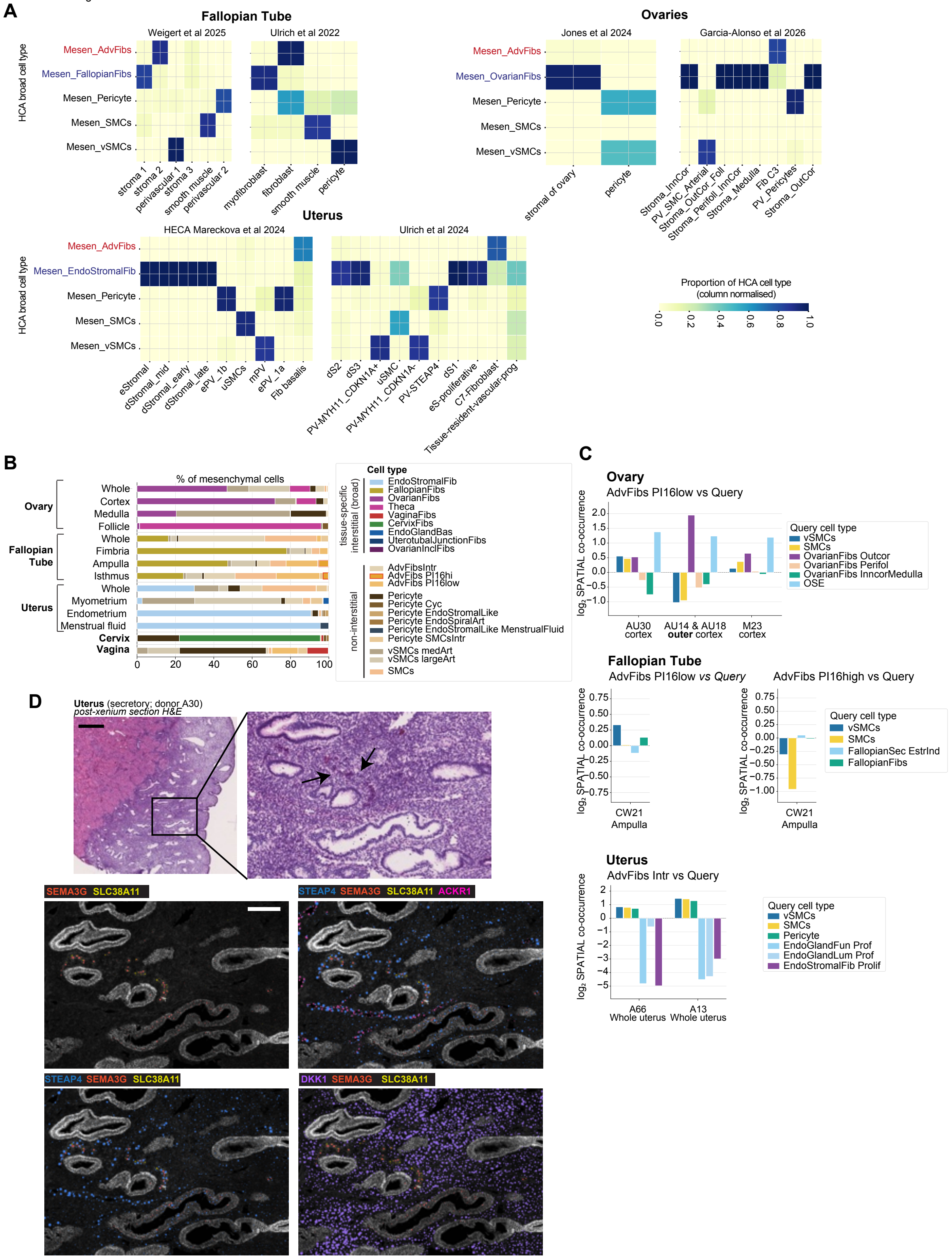


**Extended Data Fig. 3. Overview of mesenchymal cells in the Human Female Reproductive System Cell Atlas v1. A.** Confusion matrices comparing mesenchymal broad categories in the Human Female Reproductive System Cell Atlas v1 – adventitial, interstitial, pericyte, smooth muscle cells (SMCs) and vascular smooth muscle cells (vSMCs) – with analogous author-defined labels from the original published datasets, including two fallopian tubes, two ovarian and two uterine datasets. **B.** Bar plot showing the cellular composition of mesenchymal populations according to sampled organ and organ region. Colors indicate the different populations: interstitial populations are shown at the broad_celltype level, while non-interstitial populations are shown at the fine_celltype level. Population “AdvFibs PI16hi” highlighted in red. Organs shown include ovary, fallopian tube and uterus. **C.** Bar plots showing log2 spatial co-occurrence, predicted using TACCO, between adventitial populations and selected representative anatomical populations, including smooth muscle cells (SMCs), vascular smooth muscle cells (vSMCs), interstitial fibroblasts and epithelium. Co-occurrence was assessed in three ovarian sections representing the cortex from three donors, one fallopian tube section representing the ampulla, and two whole-uterus samples, both proliferative stages of the menstrual cycle, profiled using Visium HD. **D.** Visualisation of selected vascular and perivascular marker transcripts in one representative whole uterine section from adult donor A30 (reproductive age, mid-secretory phase), profiled using a bespoke Xenium 480-gene panel. The top panels show the corresponding haematoxylin and eosin (H&E) histology and a magnified view of the inset, alongside Xenium visualisation (bottom panels) of marker genes for arterial endothelium (*SEMA3G*; red), spiral artery pericytes (*SLC38A11*; yellow), venous/vessel endothelium (*ACKR1*; pink), decidualised interstitial fibroblast of the secretory phase (DKK1; purple) and classical pericytes (*STEAP4*; blue). Arrows point to spiral arteries in the tissues. Scale bar = 500μm in full thickness H&E images and 100μm in magnified insets. Adv, adventitial; Fib, fibroblast; SMC, smooth muscle cell; vSMC, vascular smooth muscle cell; InnCor, inner cortex; OutCor, outer cortex; Foll, follicle; Perifoll, perifollicular; PV, perivascular; eStromal/eS, endometrial stromal; dStromal/dS, decidualized stromal; ePV, endometrial perivascular; mPV, myometrial perivascular; uSMC, uterine smooth muscle cell; prog, progenitor; Incl, inclusion; Intr, interstitial; Cyc, cycling; Art, arteries; OSE, ovarian surface epithelium; Sec, secretory; EstrInd, estrogen induced; Fun, functionalis; Lum, luminal; Prof/Prolif, proliferative.

Extended Data Figure 4

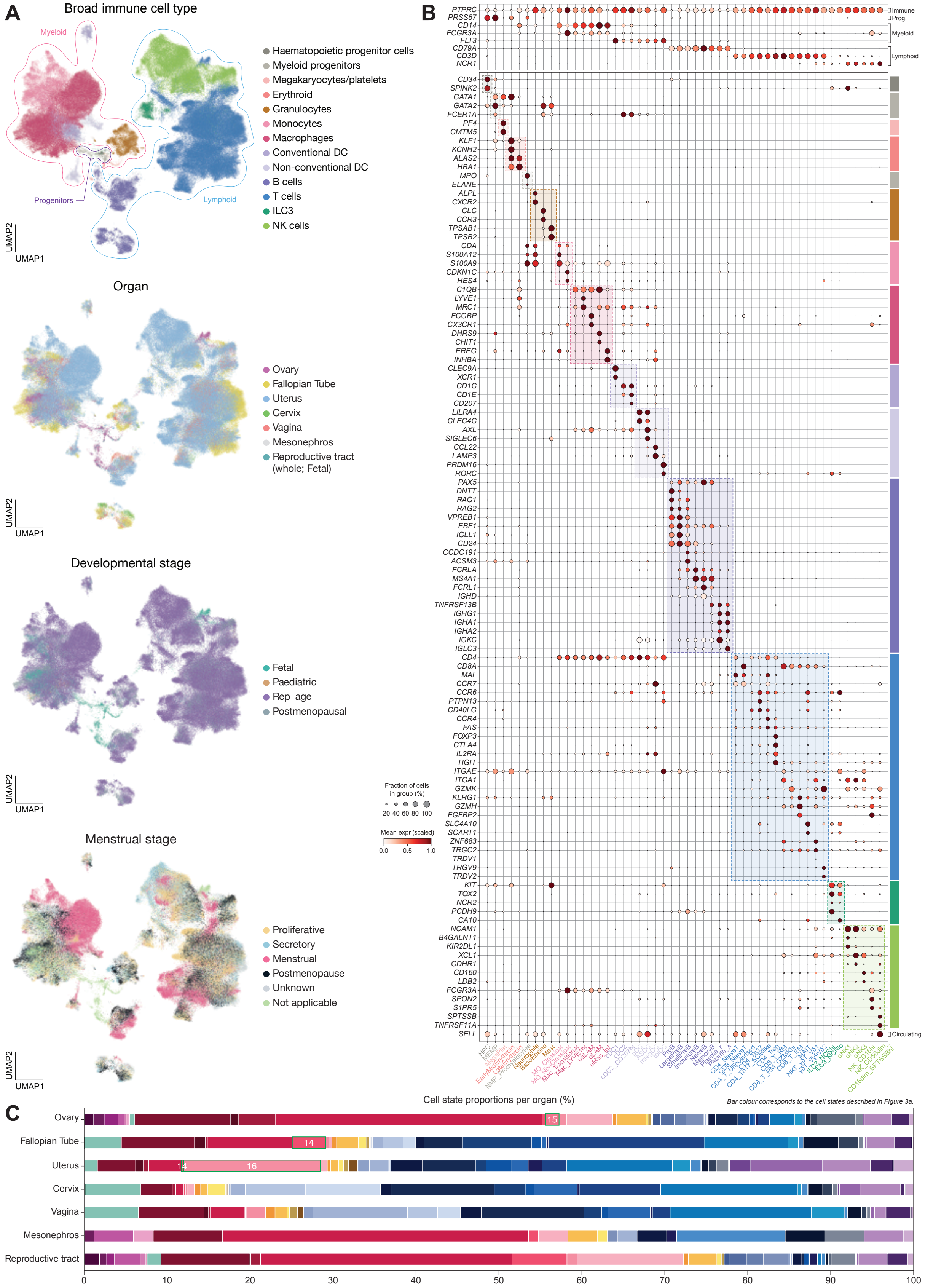


**Extended Data Fig. 4. Overview of immune cells in the Human Female Reproductive System Cell Atlas v1. A,** UMAP visualisation manifolds of scRNA-seq data for all 220,867 immune cells coloured by broad cell type, organ, developmental stage, and menstrual stage. **B,** Dot plot showing the log-transformed, min-max normalised expression of selected marker genes (x-axis) for all immune cell states (y-axis), excluding cycling cells. The colours of the bars, dotted squares, and cell state names correspond to the broad immune cell types depicted in **A. C,** Bar plot displaying the proportion of all immune cell states within each organ. Both prenatal and postnatal samples contribute to the ovary. Bars outlined in green and numbered with 14 (uftLAM), 15 (oLAM), or 16 (uMac_Inf) indicate macrophage subsets enriched in certain organs. Bar colour to cell states depicted in Figure 3a. DC, dendritic cell; ILC3, type 3 innate lymphoid cell; NK, natural killer; Prog, progenitor; HPC, haematopoietic progenitor cell; MEMP, megakaryocyte-erythroid-mast progenitor; NMP, neutrophil-myeloid progenitor; Baso, basophil; Eosino, eosinophil; MegaPlat, megakaryocyte/platelet; MO, monocyte; Mac, macrophage; uftLAM, uterine and fallopian tube lipid-associated macrophage; oLAM, ovary lipid-associated macrophage; Inf, inflammatory; Cyc, cycling; cDC, conventional dendritic cell; pDC, plasmacytoid dendritic cell; ASDC, AXL/SIGLEC6+ dendritic cell; mregDC, mature regulatory dendritic cell; tolDC, tolerizing dendritic cell; SCM, stem cell memory; Treg, regulatory T cell; MAIT, mucosal-associated invariant T cell; RA, retinoic acid. Rep_age, reproductive age; expr, expression.

Extended Data Figure 5

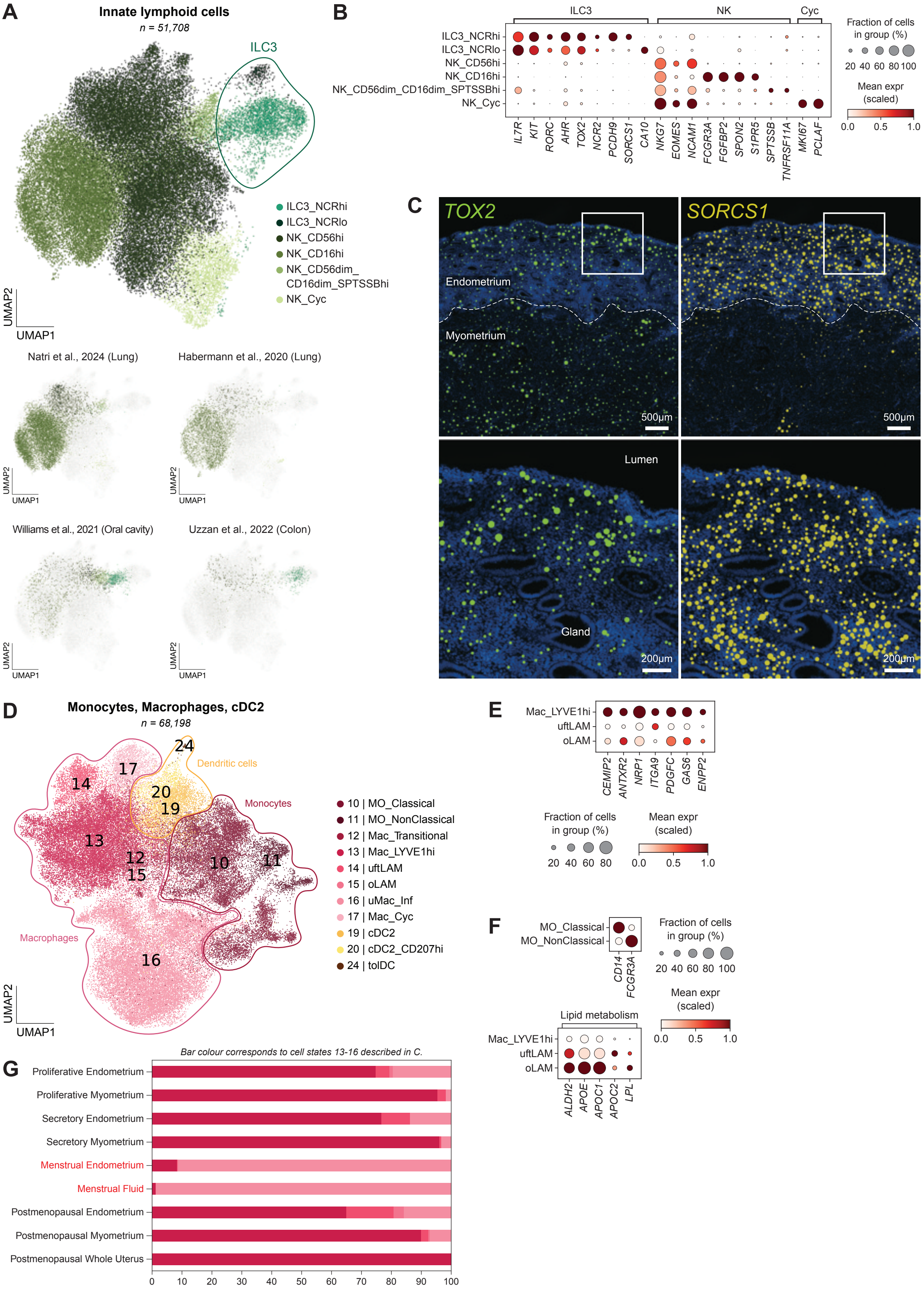


**Extended Data Fig. 5. Characterisation of ILC3 and macrophage subsets. A,** (Left) UMAP visualisation manifold of integrated scRNA-seq data of 51,708 innate lymphoid cells (ILCs), both from the Human Female Reproductive System Cell Atlas v1 (ILC3, NK cells) and from single-cell studies of diseases related to the lung [1,2], oral cavity [3], and colon [4]. Integration was performed with healthy control cells from each study. Colours indicate cell type or state, which was determined via manual annotation. (Right) Each public dataset's contribution to the integrated object. **B,** Dot plot showing the log-transformed, min-max normalised expression of selected marker genes (x-axis) for the ILC cell states (y-axis) depicted in **A. C,** Spatial distribution of transcripts for *TOX2* (left column, pan-ILC3 marker) and *SORCS1* (right column, NCR$^{hi}$ ILC3 marker) within an endometrial tissue section, taken during day 5 of menstruation, stained by the Xenium 5k panel. Bottom two images are a zoom in of the inset indicated by the white square. Dotted line indicates the endometrial-myometrial border. Generated by 10x Genomics Xenium Explorer 4.1.1. **D,** UMAP visualisation manifold of scRNA-seq data of 68,198 monocytes, macrophages, and cDC2 from the Human Female Reproductive System Cell Atlas v1. Cell state colours and numbers correspond to those depicted in Figure 3a. **E,** Dot plot showing the log-transformed, min-max normalised expression of selected genes upregulated in LYVE1$^{hi}$ macrophages compared to uftLAMs and oLAMs. **F,** Dot plot showing the log-transformed, min-max normalised expression of (top) *CD14* and *CD16* (*FCGR3A*), which differentiate classical and non-classical monocytes, and (bottom) lipid metabolism genes that are upregulated in LAMs. **G,** Bar plot displaying the proportion of LYVE1$^{hi}$ macrophages, uftLAMs, oLAMs, and inflammatory macrophages within uterine tissues (endometrium, myometrium, menstrual fluid, whole uterus) across menstrual cycle stage (proliferative, secretory, menstrual, postmenopausal). Bar colors correspond to the cell states depicted in **D.** ILC3, type 3 innate lymphoid cell; NK, natural killer; Cyc, cycling; expr, expression; MO, monocyte; Mac, macrophage; uftLAM, uterine and fallopian tube lipid-associated macrophage; oLAM, ovary lipid-associated macrophage; Inf, inflammatory; cDC, conventional dendritic cell; tolDC, tolerizing dendritic cell.

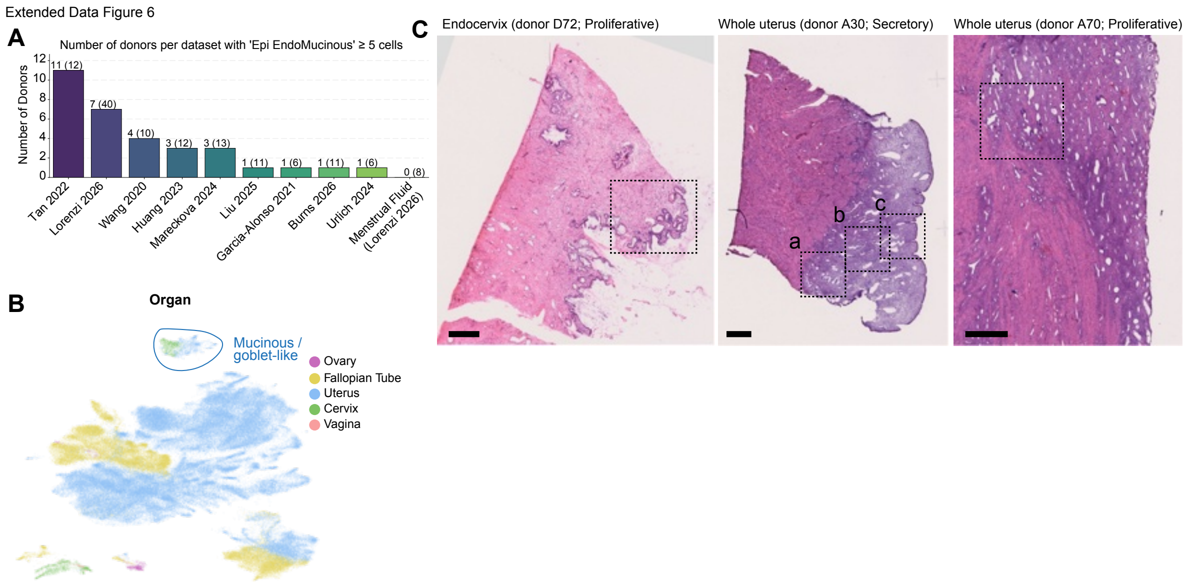


**Extended Data Fig. 6. Overview of endometrial/endocervical mucinous epithelial cells in the Human Female Reproductive System Cell Atlas v1. A.** Bar plot showing the number of donors with at least five cells classified as "EndoMucinous" epithelial cells per dataset. Numbers above the bars indicate donors passing this threshold, while numbers in parentheses indicate the total number of donors in each dataset. **B.** Batch-corrected Uniform Manifold Approximation and Projection (UMAP) embedding of the scRNA-seq dataset of all epithelial cells (n=205,208; "broad_celltype" labels) from the postnatal donors in the Human Female Reproductive System Cell Atlas v1, coloured by organ (matching **Figure 4a**). **C.** Visualisation of haematoxylin and eosin (H&E) histology and the magnified view of the inset (matching **Figure 4D**) for one endocervical section from donor D72 (reproductive age, proliferative phase) with a magnified view of the endocervical glands (left column); one whole uterine section from donor A30 (reproductive age, secretory phase) with three magnified regions “a-c” showing *MUC5B* detection in the basalis, functionalis and lumen, respectively (middle columns); and one whole uterine section from donor A66 (reproductive age, proliferative phase) with one magnified region showing *MUC5B* detection in the basalis (right column). Epi, epithelial.

Extended Data Figure 7

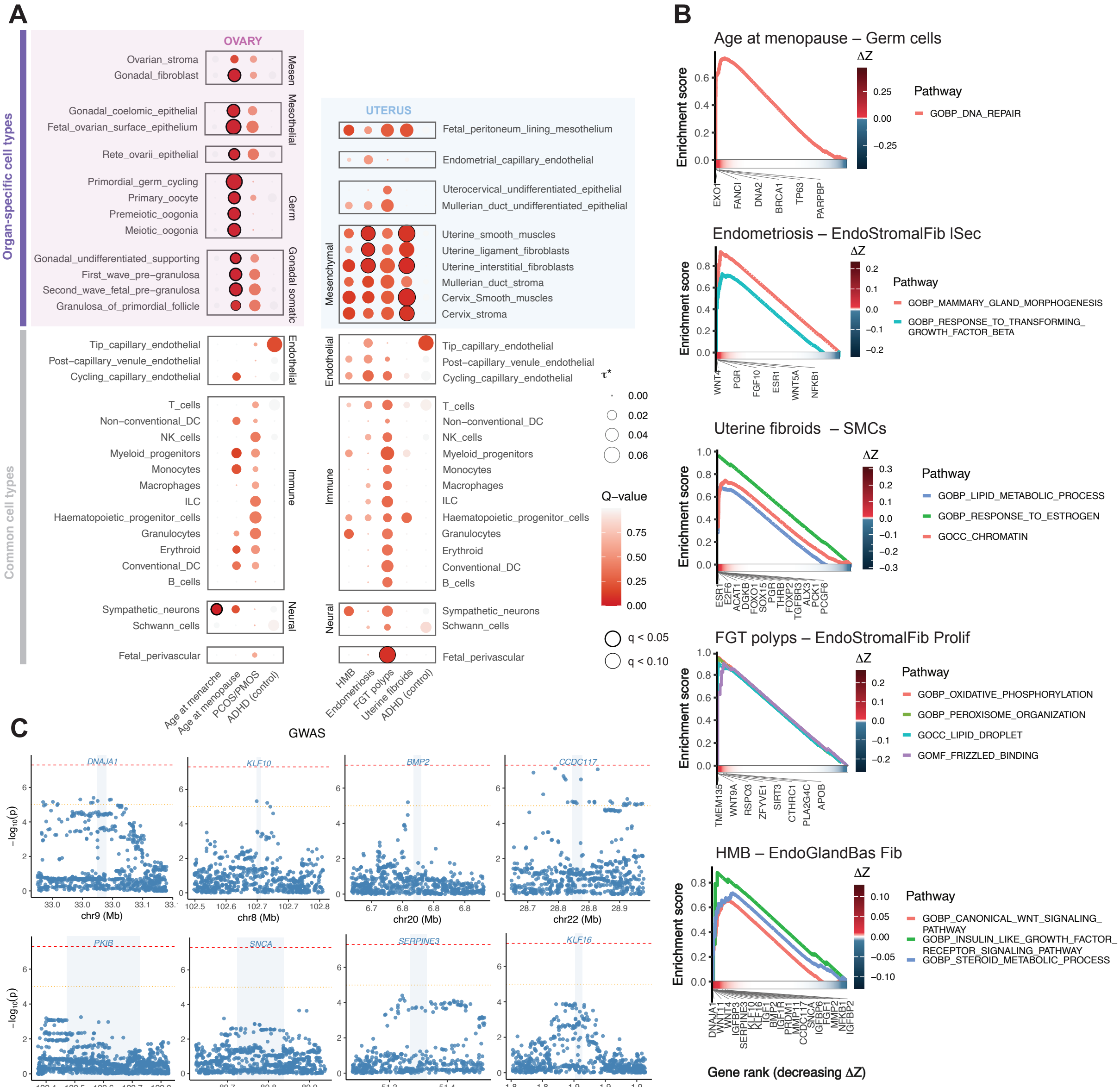


**Extended Data Fig. 7. Integration of GWAS with the Human Female Reproductive System Cell Atlas v1 and leave-one-out gene decomposition analysis. A,** Cell type enrichment of disease heritability in the specifically expressed genes per cell type in the fetal Human Female Reproductive System Cell Atlas v1. The dot plots show enrichment for both uterine and ovary specific traits (x-axis) computed using LDSC, with ovary-specific and uterus-specific cell types respectively highlighted in pink and blue and non-specific cell types not highlighted. Cell types (y-axis) are grouped by lineages and enrichment is shown in dots where the size represents the standardised effect $\boldsymbol{\tau}^*$ – the additive change in heritability explained by a 1 standard deviation increase in the gene expression specificity (CELLEX) annotation – and the colour represents significance q-value as -log10(q), corrected for multiple testing. Bold black circles indicate enrichment at $q<0.05$ and light black circles indicate enrichment at $q<0.10$. Data points with $\boldsymbol{\tau}^*<0$ were set to 0 and corresponding q-values set to 1. **B,** fGSEA results from the leave-one-out gene decomposition analysis from the adult HCA enrichment analysis. For each top trait–cell type association per trait, we computed the contribution $\Delta Z$ of each gene to the enrichment. Given that Gene Ontology pathways are often redundant and not transferrable to uterine or ovary-specific processes, we highlight specific pathways among the top nominally significant gene sets **(Supplementary Table 4d)** and leading edge genes relevant to known biological processes in the female reproductive system. **C,** Scatter plots of GWAS associations with HMB in genomic windows around *BMP2*, *KLF10*, *CCDC117*, *DNAJA1*, *KLF16*, *PKIB*, *SERPINE3*, *SNCA*. The x-axis shows genomic coordinates in megabases across the specified chromosome, extended by 100kb windows from either end of each gene body which is highlighted and labelled in blue. The y-axis shows -log10 p-values for association between each SNP and HMB, with genome-wide significance (5e-8) as a red dotted line and suggestive significance (1e-5) as an orange dotted line. DC, dendritic cell; NK, natural killer; ILC, innate lymphoid cell; PCOS, polycystic ovary syndrome; PMOS, polyendocrine metabolic ovarian syndrome; ADHD, attention deficit hyperactivity disorder; HMB, heavy menstrual bleeding; FGT, female genital tract; GOBP, Gene Ontology biological processes; GOCC, Gene Ontology cellular components; GOMF, Gene Ontology molecular function; GWAS, genome-wide association study.

Extended Data Figure 8

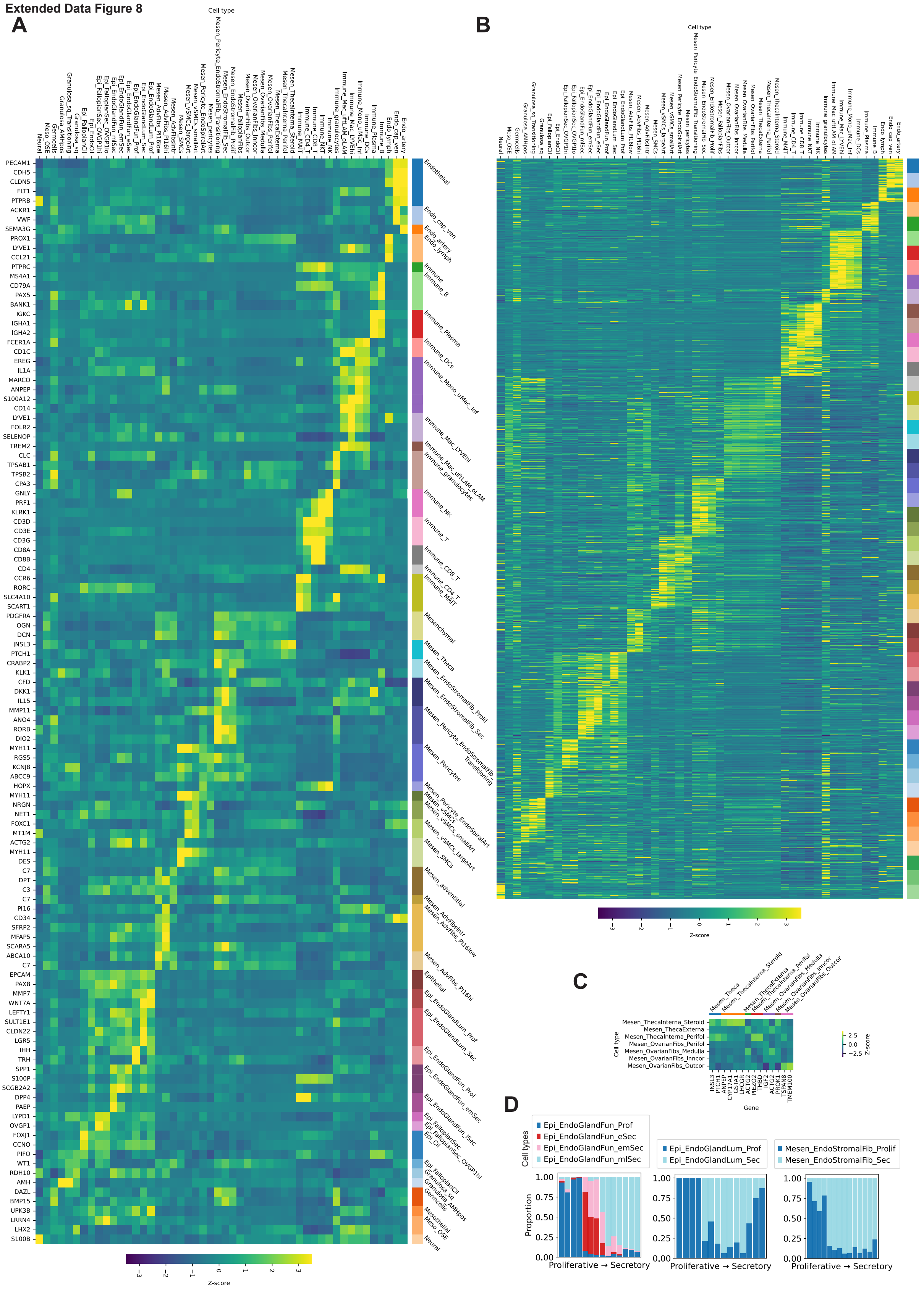


**Extended Data Fig. 8. Chromatin accessibility atlas cell type annotation. A,** Heatmap of marker gene activity scores in annotated cell types from the Human Female Reproductive System Cell Atlas v1. Gene activity scores were computed using chromatin accessibility in and around each gene body in ArchR. The colour of each square is the row-wise Z-score of gene activity for each given gene (y-axis) in a given cell type (x-axis). Marker genes are grouped by relevant cell type or lineage on the right-hand y-axis. **B,** Heatmap of HCA top 20 marker genes gene activity scores in annotated cell types. The Human Female Reproductive System Cell Atlas v1 transcriptomic atlas was regrouped according to the scATAC-seq cell types and top 20 marker genes per cell type computed using a TF-IDF approach. The colour of each square is the row-wise Z-score of gene activity for each given gene (y-axis) in a given cell type (x-axis). The genes are grouped by cell type, where the order of the colour bars corresponding to 20 genes for each cell type is the same as the cell types on the x-axis. **C,** Ovarian mesenchymal cell markers in annotated cell types. Due to the difficulty of characterising ovarian fibroblasts and marker genes distinguishing between ovarian fibroblasts not being specific to ovarian fibroblasts, these markers were not included in **A**. Here we replotted **A** with cell types on the y-axis and genes on the x-axis to show distinction between different annotated ovarian fibroblast cell types consistent with RNA-derived markers. **D,** Proportion of cells from different menstrual stages in donors across the menstrual cycle. Within each group of cell types (Epi_EndoGlandFun – epithelial endometrial glandular functional, Epi_EndoGlandLum – epithelial endometrial glandular luminal, Mesen_EndoStromalFib – mesenchymal endometrial stromal fibroblasts), we show the proportion of cells per donor annotated as different cell states associated with menstrual cycle progression, ordering donors by menstrual cycle day. The two proliferative donors did not have menstrual cycle day information so were randomly added in the middle. Endo, endothelial/endometrial; Mesen, mesenchymal; Epi, epithelial; Meso, mesothelial; cap, capillary; ven, venule; lymph, lymphatic; DC, dendritic cell; Mono, monocyte; Mac, macrophage; Inf, inflammatory; uftLAM, uterine and fallopian tube lipid-associated macrophage; oLAM, ovary lipid-associated macrophage; NK, natural killer; NKT, natural killer T cell; MAIT, mucosal-associated invariant T cell; Perifol, perifollicular; Fib, fibroblast; Inncor, inner cortex; Outcor, outer cortex; Prolif/Prof, proliferative; Sec, secretory; Art, arteries; vSMC, vascular smooth muscle cell; SMC, smooth muscle cell; Adv, adventitial; Intr, interstitial; Lum, luminal; Fun, functionalis; eSec, early secretory; emSec, early/mid secretory; mlSec, mid/late secretory; Cil, ciliated; sq, squamous; OSE, ovarian surface epithelium.

**Extended Data Figure 9**

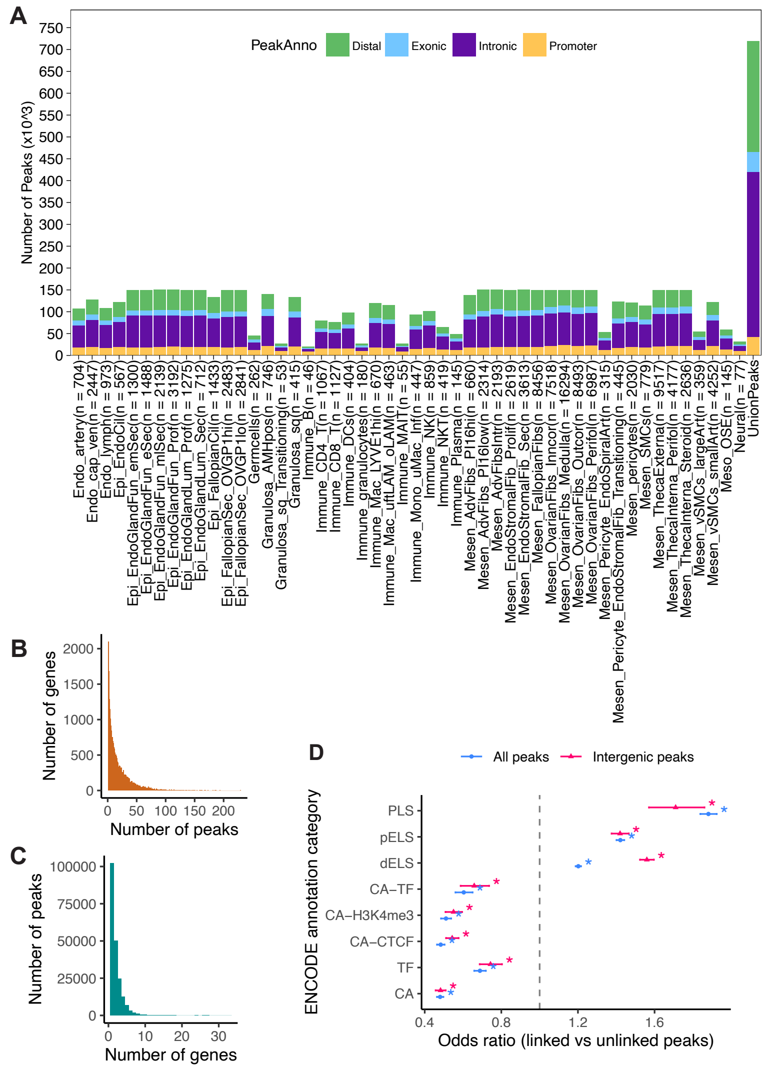


**Extended Data Fig. 9. Peak calling and peak–gene linkage. A,** Summary of peak calling in the chromatin accessibility atlas. Peaks (y-axis) were called iteratively per cell type (x-axis) then overlapped to find consensus peaks (“UnionPeaks” on the x-axis) to avoid losing cell type specific peaks during peak calling. Peaks are classified into distal peaks (>1kb from a gene body), exonic, intronic, or promoter (within 1kb upstream of the transcription start site), differentiated by colour. **B,** Histogram of the number of genes (y-axis) linked to different numbers of peaks (x-axis), showing a right tail of genes associated with many peaks. **C,** Histogram of the number of peaks (y-axis) linked to different numbers of genes (x-axis), showing a much smaller right tail of peaks associated with many genes than **B**. **D,** Relative enrichment of peaks with a gene linked versus unlinked peaks for different ENCODE annotations, considering all peaks (blue) and removing peaks within gene bodies (intergenic peaks in pink). The annotations are promoter-like signatures (PLS), proximal enhancer-like signatures (pELS), distal enhancer-like signatures (dELS), the intersection of ENCODE chromatin accessibility and predicted transcription factor binding sites (CA-TF), the intersection of ENCODE chromatin accessibility and H3K4me3 sites (CA-H3K4me3), the intersection of ENCODE chromatin accessibility and CTCF sites (CA-CTCF), predicted transcription factor binding sites (TF) and ENCODE chromatin accessibility (CA). All enrichments (>0) and depletions (<0) shown were significant after multiple testing, shown with an asterisk. Endo, endothelial/endometrial; Epi, epithelial; Mesen, mesenchymal; Meso, mesothelial; cap, capillary; ven, venule; lymph, lymphatic; Cil, ciliated; Fun, functionalis; emSec, early/mid secretory; eSec, early secretory; mlSec, mid/late secretory; Prof/Prolif, proliferative; Lum, luminal; Sec, secretory; sq, squamous; DC, dendritic cell; Mac, macrophage; uftLAM, uterine and fallopian tube lipid-associated macrophage; oLAM, ovary lipid-associated macrophage; MAIT, mucosal-associated invariant T cell; Mono, monocyte; Inf, inflammatory; NK, natural killer; NKT, natural killer T cell; Adv, adventitial; Fib, fibroblast; Intr, interstitial; Inncor, inner cortex; Outcor, outer cortex; Perifol, perifollicular; Art, arteries; SMC, smooth muscle cell; vSMC, vascular smooth muscle cell; OSE, ovarian surface epithelium.
